# Principles of human pre-60*S* biogenesis

**DOI:** 10.1101/2023.03.14.532478

**Authors:** Arnaud Vanden Broeck, Sebastian Klinge

## Abstract

During early stages of human large ribosomal subunit (60*S*) biogenesis, an ensemble of assembly factors establishes and fine-tunes the essential RNA functional centers of pre-60*S* particles by an unknown mechanism. Here, we report a series of cryo-electron microscopy structures of human nucleolar and nuclear pre-60*S* assembly intermediates at resolutions of 2.5-3.2 Å. These structures show how protein interaction hubs tether assembly factor complexes to nucleolar particles and how GTPases and ATPases couple irreversible nucleotide hydrolysis steps to the installation of functional centers. Nuclear stages highlight how a conserved RNA processing complex, the rixosome, couples large-scale RNA conformational changes to pre-rRNA processing by the RNA degradation machinery. Our ensemble of human pre-60*S* particles provides a rich foundation to elucidate the molecular principles of ribosome formation.

**One-Sentence Summary:** High-resolution cryo-EM structures of human pre-60S particles reveal new principles of eukaryotic ribosome assembly.

## Main Text

Protein synthesis in all cells requires ribosomes, two-subunit RNA-protein molecular machines in which the RNA component is responsible for both decoding of messenger RNA within the ribosomal small subunit (40*S*) as well as peptide bond formation within the ribosomal large subunit (60*S*). Since the discovery of ribosome assembly in human cells, more than 200 ribosome assembly factors have been identified that catalyze the modification, processing and folding of ribosomal RNA, initially in the nucleolus and subsequently in the nucleus and cytoplasm (*1*–*3*). The large ribosomal subunit rRNAs (5*S*, 5.8*S* and 28*S* rRNA) are generated from two transcripts: a transcript containing the 5*S* rRNA and a separate 47*S* pre-rRNA that contains the 5.8*S* and 28*S* rRNA in addition to the rRNA of the small subunit. Following the transcription of the 47*S* pre-rRNA in the nucleolus and a cleavage reaction separating precursors of both subunits, the first stable nucleolar large subunit pre-rRNA is a 32*S* precursor in which the 5.8*S* and 28*S* rRNA are joined by internal transcribed spacer 2 (ITS2).

Traditionally, eukaryotic 60*S* assembly has been studied in the yeast system ((*4*) and references therein), (*5*–*10*), which has offered several molecular snapshots of early nucleolar assembly intermediates from *Saccharomyces cerevisiae* (*11*–*13*). However, the mechanisms that drive early assembly forward remain unknown. The absence of key transition states has so far precluded a mechanistic understanding of how essential energy consuming enzymes such as ATPases, including DEAD/H-box helicases and AAA+-ATPases, as well as GTPases can bring about irreversible transitions and the formation of catalytic centers of the eukaryotic large ribosomal subunit, such as the peptidyl transferase center (PTC) (*14, 15*). Thus, the mechanisms that enable early eukaryotic large ribosomal subunit assembly to proceed in a controlled and unidirectional manner remain poorly understood.

Towards the end of nucleolar maturation, pre-60*S* assembly intermediates already contain functional centers, including the peptidyl transferase center (PTC), the GTPase-associated center (GAC), the L1 stalk, as well as a central protuberance (CP) containing an unrotated 5*S* RNP (*16*). However, these functional centers have not yet reached their mature conformations, requiring an ensemble of nuclear ribosome assembly factors for their completion. Although several structures covering nuclear stages of large ribosomal subunit assembly in yeast have been described (*5*–*7, 16, 17*), major gaps in our mechanistic understanding of this process remain. An essential transition during nuclear pre-60*S* maturation is the rotation of the 5*S* RNP during conversion of a Nog2-bound particle (*16*) to a Rix1-Midasin bound particle (*7*), which is catalyzed by Midasin (Mdn1). Importantly, only after the rotation of the 5*S* RNP does ITS2 processing result in the mature 5.8*S* rRNA, but how these two irreversible events are linked has remained unclear. Consistent with functional coupling between 5*S* RNP rotation and ITS2 processing, mutations or depletions of assembly factors involved in 5*S* RNP rotation or ITS2 processing result in the accumulation of either unprocessed pre-ribosomal RNA or 5.8*S* precursors containing parts of ITS2 (*18*–*22*). Interestingly, the accumulation of the same pre-RNA species is also observed upon depletion of the endonuclease cleaving ITS2 (yeast Las1, human LAS1L) and components of the RNA exosome, respectively (*23, 24*). While these data suggest coupling of 5*S* RNP rotation with ITS2 cleavage and processing by Las1/LAS1L and the RNA exosome, the molecular basis of this mechanism remains elusive.

Insights into human large ribosomal subunit assembly are still in their infancy as only a few structures of very late stages of pre-60*S* maturation have been obtained through the ectopic expression of an affinity-tagged nuclear export factor (*25*). Due to the biochemical properties of the human nucleolus, an inability to obtain endogenous early pre-60*S* assembly intermediates has so far limited our understanding of the molecular mechanisms responsible for their unidirectional assembly and maturation. Furthermore, within the human nuclear ribosome assembly pathway, it is unclear how an evolutionarily expanded multiprotein complex - the rixosome (*26*) - can integrate different functional activities to ensure that cleavage of ITS2 by LAS1L (*27*) and the subsequent RNA exosome-mediated processing of ITS2 are efficiently coupled.

From an evolutionary perspective, the 32*S* pre-rRNA of the late nucleolar intermediates in humans is much larger than its yeast 27*S* counterpart due to the presence of several expansion segments in ITS2 and the 28*S* rRNA and it is currently unknown if these expansions are required for ribosome assembly.

In this study, we have combined human genome editing and biochemistry to permeabilize human nucleoli and nuclei to determine high-resolution cryo-EM structures of 24 maturing human pre-60*S* particles at resolutions of 2.5-3.2 Å. Of these, eight main nucleolar structures elucidate how multiple levels of regulation are employed to drive unidirectional 60*S* maturation by integrating tethered multiprotein complexes with ATPases, such as DEAD-box RNA helicases, and GTPases that catalyze key transitions during the assembly of functional centers. Four main nuclear structures reveal the mechanism by which 5*S* RNP rotation is coupled to ITS2 processing and how the human rixosome plays a central role in mediating these steps. By employing engineered rRNAs in human cells, we have identified elements of the human ITS2 and 28*S* rRNA that are critical for ribosome assembly and nuclear export.

### Cryo-EM structures of maturing human pre-60*S* particles

To obtain structural insights into early stages of human large ribosomal subunit assembly, we have biallelically affinity tagged the endogenous ITS2-associated assembly factor MK67I (**fig. S1A-D**). The biochemical affinity purification of MK67I enabled the direct isolation of human pre-60*S* particles containing ITS2 for cryo-EM analysis. These nucleolar and nuclear stages of assembly are associated with the expected nucleolar (32*S*) and nuclear (28*S*, 12*S*, and 8*S*) pre-rRNAs (**figs. S1E-I, S2)**. A total of 172,699 cryo-electron micrographs were collected on a Titan Krios microscope equipped with a K3 detector. After initial data curation and extensive 3D classification, we identified 12 nucleolar stages of assembly, of which the main eight states are here referred to as A to H, and 12 nuclear stages of assembly, of which the four main states are referred to as I to L (**figs. S3-5**). Focused classifications and refinements were used to obtain high-resolution composite maps (2.5-3.2 Å) (**figs. S6-15, tables S1**,**2**). These reconstructions enabled the assignment and precise model building of human pre-ribosomal RNAs (5.8*S*, ITS2, 28*S* and 5*S* rRNA), 41 ribosomal proteins and 47 nucleolar/nuclear ribosome assembly factors (**tables S3**,**4**). Differences in the pre-rRNA folding states and the presence or absence of distinct assembly factors enabled the assignment of states A-H and states I-L into nucleolar and nuclear assembly pathways, respectively (**Fig. 1, fig. S16-S19**).

**Fig. 1.**
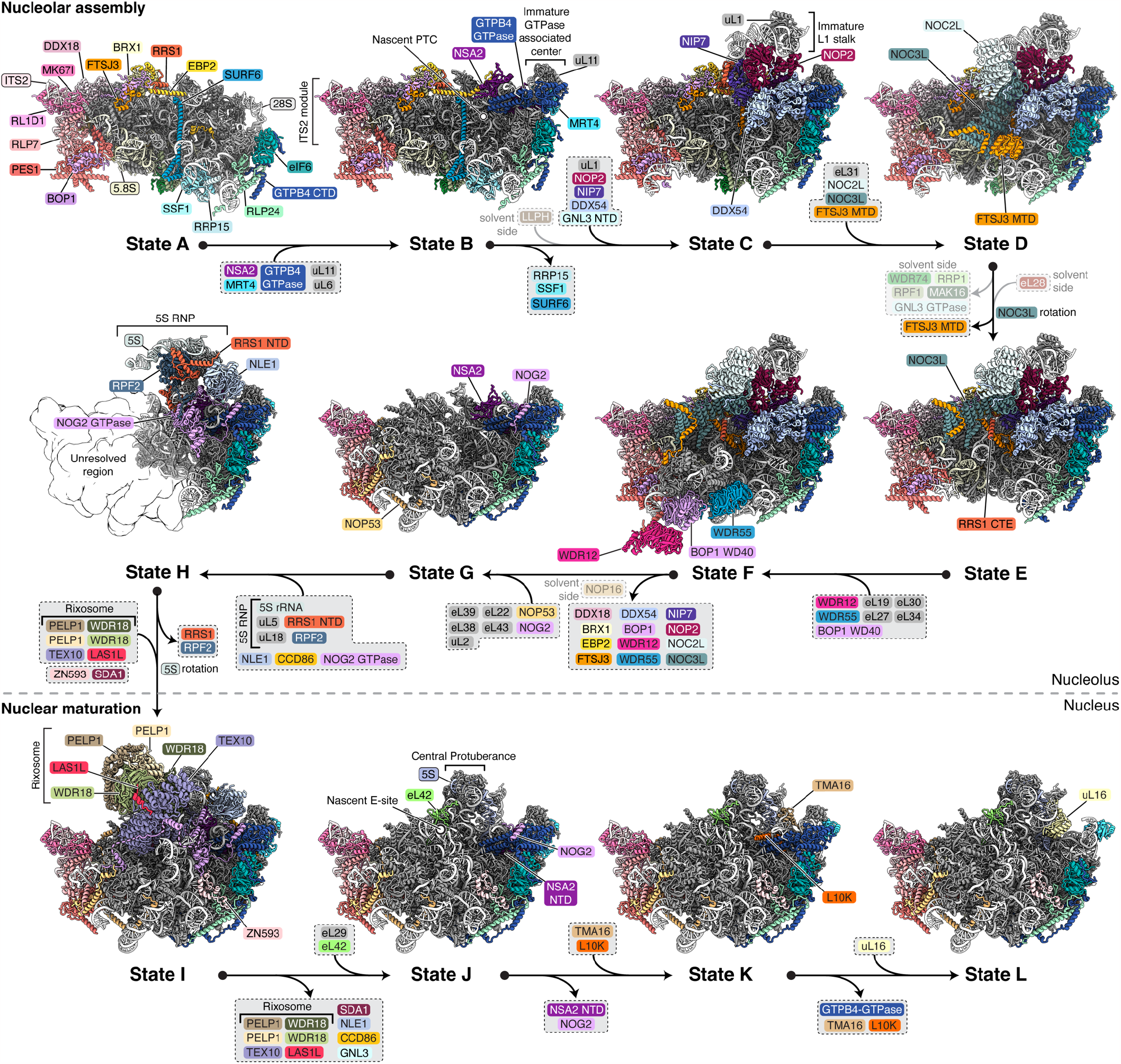
Nucleolar assembly and nuclear maturation of human pre-60*S* particles. Simplified human pre-60*S* assembly pathway with crown view of atomic models of states A to L with ribosomal proteins in grey and color-coded pre-rRNAs and ribosome assembly factors. Proteins that become ordered or depart between different assembly states are indicated on arrows.

We have identified parallel assembly pathways at the beginning of the nucleolar assembly (states A-D), characterized by the presence or absence of the MAK16-associated assembly factor complex (MAK16, WDR74 (yeast Nsa1), RPF1, RRP1 and the GTP-binding domain of GNL3 (yeast Nug1)) that are removed by NVL2 (*15*) (**fig. S16**,**17**). We also observe parallel maturation pathways with the stepwise maturation of ITS2 elements from state I to L during nuclear stages of assembly (**figs. S18**,**19**). Importantly, these ITS2-centered events can occur concurrently with other maturation events that focus on the fine-tuning of functional centers of progressing nuclear pre-60*S* particles. These include the cooperative formation of the E-site (state J), the progressive fine-tuning of the PTC through irreversible exchange of assembly factors (state K), and the maturation of the P-site with the installation of uL16 prior to nuclear export (state L).

The unprecedented scale of our cryo-EM dataset allowed us to observe the progression of the human large ribosomal subunit assembly as a fine-grained series of molecular snapshots that enable a precise mechanistic explanation of how ribosome assembly factors act in concert, the identification of human-specific adaptations and the visualization of covalent modifications of pre-ribosomal RNA as a function of ribosome assembly. This ensemble of states now provides a high-resolution perspective on nucleolar assembly and nuclear maturation of the human large ribosomal subunit as described below.

### High-level organization and hierarchy of pre-rRNA and protein incorporation

A unique principle of human pre-60*S* assembly is the early establishment of hubs that organize architectural pre-rRNA elements and tether ribosome assembly factor complexes to coordinate different steps of assembly (**Fig. 2**). Ribosomal RNA root helices, which form the origin of each of the six subdomains (I-VI) of the 28*S* rRNA, are already positioned in the first visualized post-transcriptional assembly state (state A). The early establishment of root helices around the assembly factor complex SURF6-SSF1-RRP15 assists in the correct positioning of each rRNA domain, thereby facilitating subsequent folding events that occur either within a domain or across domains (**Fig. 2A**). Ribosome assembly factors are not only responsible for the correct placement of central architectural rRNA elements within state A, but also prevent premature folding of key rRNA elements (**figs. S20**,**21**).

**Fig. 2.**
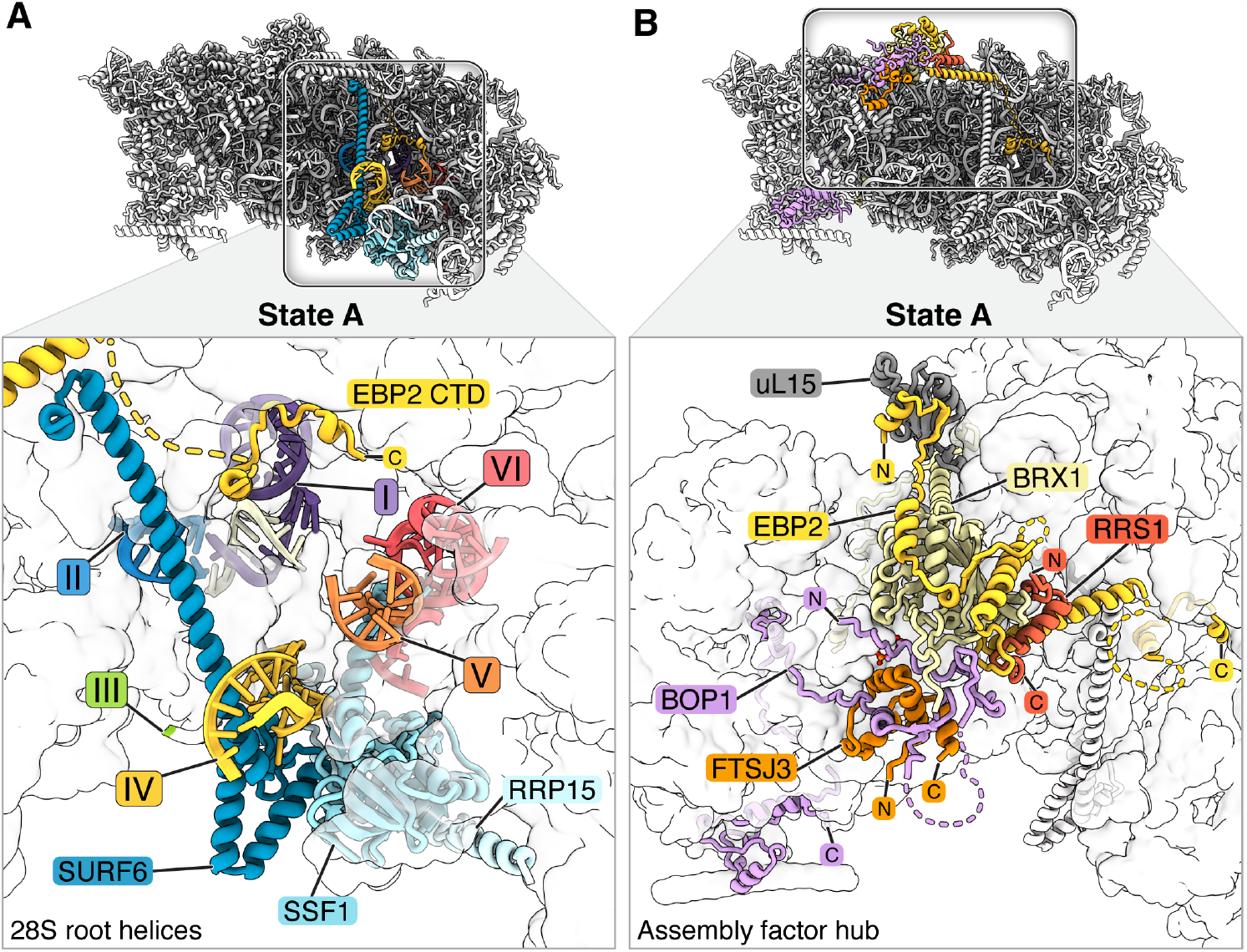
Early clustering of pre-rRNA and assembly factor modules. (**A**) Crown view of state A showing color-coded root helices of domains I-VI and assembly factors. (**B**) Crown view of state A highlighting a protein-protein interaction hub with color-coded assembly factors.

Comparable to the organization of central elements of pre-rRNA, a protein-protein interaction hub is established in state A near domain I, which tethers ribosome assembly factor complexes to the maturing particles, thereby allowing for progression through subsequent stages of assembly (**Fig. 2B**). This protein-protein interaction hub is formed around an N-terminal region of BOP1 to which an extended BRX1-EBP2 complex and peptide elements of FTSJ3 and RRS1 are bound.

Previous studies have highlighted that eukaryotic ribosome assembly is controlled through both the TOR and casein kinase II pathways (*28, 29*). Interestingly, we have identified two mammalian-specific post-translational modifications (PTMs) in the N-terminal segment of BOP1 (phosphoserines 126 and 127), which are present within a sequence motif resembling casein kinase II substrates (*30*). The central location of these modified residues, which are present in all states containing BOP1 (states A-F), raises the possibility that this set of PTMs enables modulation of human ribosome biogenesis within cellular metabolism through signaling pathways (**fig. S21B**).

During nucleolar assembly stages covering states A to F, the protein-protein interaction hub (**Fig. 2B**) performs three major organizing functions: first, the presence of a peptide of the RNA 2’-O-methyltransferase FTSJ3 (yeast Spb1) suggests that tethering enables 2’-O-methylation of 28*S* rRNA as well as recruitment of the NOC2L-NOC3L complex and ribosomal proteins at different stages of assembly (**fig. S21B**). By interacting with all these proteins, tethered ribosome assembly factors and ribosomal proteins are already present before their involvement in subsequent stages of assembly. Second, the presence of a C-terminal extension of RRS1 provides a mechanism for recruitment and later incorporation of the L1 stalk and the 5*S* RNP via the RRS1-RPF2 complex that binds to 5*S* rRNA and ribosomal proteins uL18 (RPL5) and uL5 (RPL11) (**Fig. 1, fig. S21D**). Third, EBP2 has dual functions that include both the prevention of premature rRNA folding as well as the stabilization of the NSA2-GTPB4 complex in state B (**Figs. 1, 2, figs. S20B, S21C**).

Collectively, the organization of pre-rRNA elements and ribosome assembly factor complexes in state A lays the foundation for many of the subsequent irreversible transitions that require the presence of already tethered assembly factors to efficiently stabilize new pre-rRNA folding states.

### GTPB4 catalyzes the initial installation of the peptidyl transferase center

Key transitions during ribosome assembly are controlled through irreversible steps that involve nucleotide hydrolysis, RNA cleavage or the generation of force (*4, 31*). An essential family of GTPases conserved in all domains of life that acts as a central regulator of large ribosomal subunit assembly includes the human protein GTPB4 (Nog1 in yeast, ObgE in *E*.*coli*) (*16, 32*–*34*). So far, the limiting resolution of endogenous assembly intermediates obtained from any species and the lack of transition states of maturing particles has precluded a precise mechanistic understanding of the GTPase cycle of GTPB4 and its homologues during assembly.

While the C-terminus of GTPB4 is already bound to eIF6 and RLP24 in state A (**Fig. 3A**), the GDP-bound GTPB4 GTPase domain first appears in state B and remains GDP-bound in all subsequent nucleolar and nuclear states that we observe (**Figs. 1, 3B, figs. S22, S23**). As state B is the first state in which parts of the nascent Peptidyl Transferase Center (PTC; helices 89 & 91) and the GTPase Associated Center (GAC; remodeled ES13 & helices 42-44) appear, these data indicate that the incorporation of these RNA elements is coupled to the GTPase activity of GTPB4. The incorporation of GTPB4 after GTP hydrolysis is therefore similar to other enzymes with ATPase or methyltransferase activity (*10, 35*). Consistent with this model, conserved N-terminal residues of NSA2 that have been implicated in activating GTPase activity in yeast (*36*) are positioned near the GTPase active site (**fig. S22B**) and dominant negative mutations near the GTPB4 active site result in nucleolar accumulation of GTPB4/Nog1 in mouse and yeast, consistent with trapped particles equivalent to state A in which only the C-terminal fragment of GTPB4 is bound (*32, 33*).

**Fig. 3.**
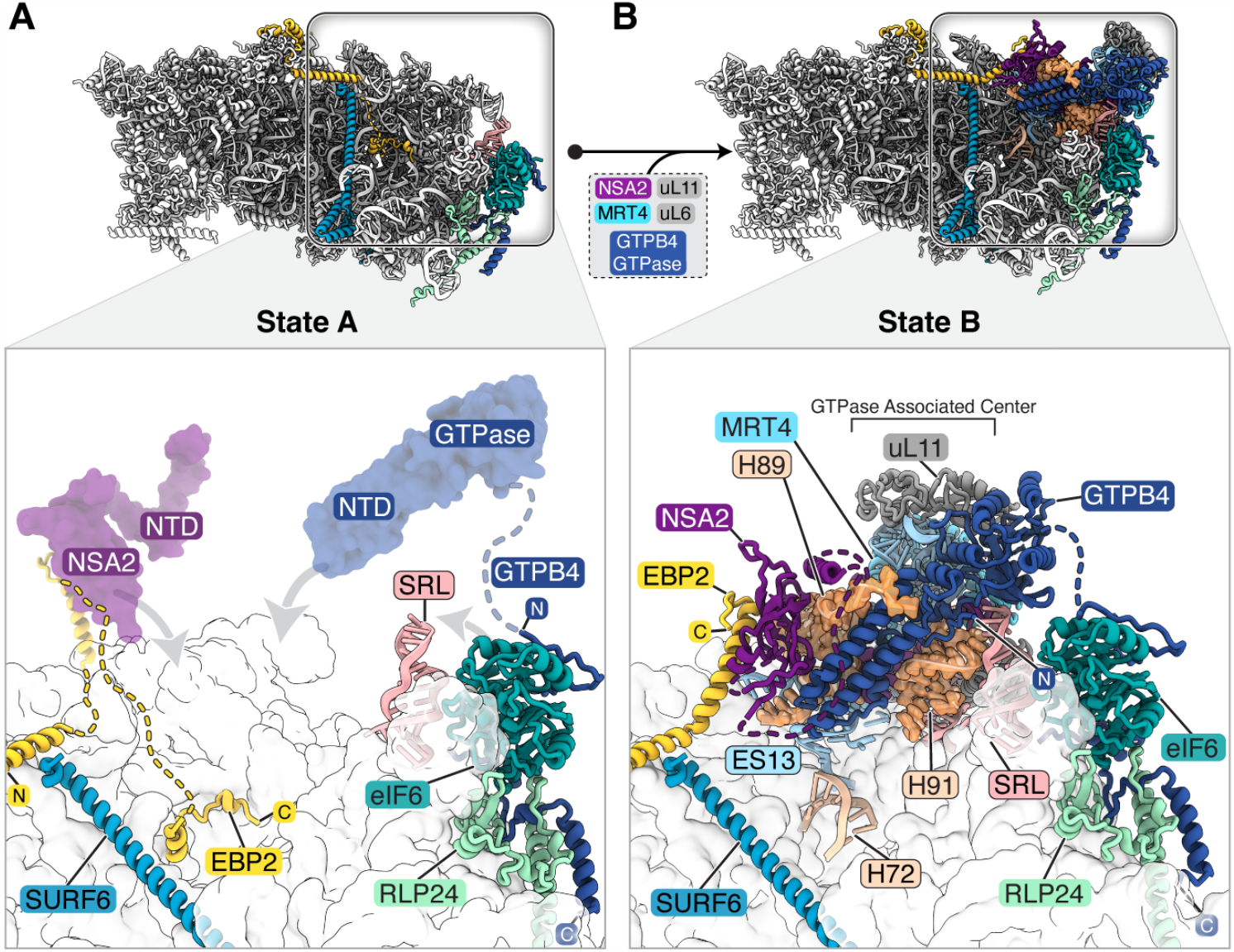
GTPB4 drives early PTC installation. (**A, B**) Crown views of states A (A) and B (B) showing selected assembly factors. Lower panels show a zoomed view in which arrows show flexible elements of GTPB4, NSA2 and EBP2 in state A (A) before these are incorporated in state B (B). Assembly factors are color-coded and newly integrated PTC rRNA are represented as orange surface.

Importantly, state B represents a key transition state as early assembly factors bridging between domains I, II and VI (SURF6-SSF1-RRP15) are still present while EBP2 has been remodeled to accommodate helix 72 and stabilize NSA2 and the GTPase domain of GTPB4 in the post-hydrolysis state (GDP-bound) (**Figs. 3B, fig. S22B**). In state B, crucial rRNA elements around GTPB4, such as expansion segment 13 (ES13) and parts of helix 89 (H89), adopt immature conformations compared to the subsequent state C, in which the L1 stalk is incorporated (**fig. S22C**,**D**).

Together, these data allow us to postulate a universal mechanism of early GTPase-mediated installation of the PTC that rationalizes existing data from bacterial and eukaryotic systems. In contrast to the prevalent model in which GTP hydrolysis triggers dissociation of GTPB4 and its homologues in yeast and *E*.*coli* during late stages of assembly, our model postulates that GTP hydrolysis is used to dock GTPB4 and its homologues early as a quality control step to ensure correct PTC installation.

### DDX54 couples irreversible rRNA remodeling to PTC assembly

Following the GTPase-mediated partial installation of the PTC, the presence of GTPB4 in state B provides a unique binding site for the conserved DEAD-box RNA helicase DDX54 (Dbp10 in yeast) that has been implicated in PTC maturation downstream of GTPB4 (*37*–*39*). In addition to the two characteristic RecA lobes shared by all DEAD-box RNA helicases, DDX54 contains a structured C-terminal region (Elbow), N- and C-terminal extensions (NTE, CTE), as well as a C-terminal tail (CTT), providing substrate specificity as observed in other DEAD-box enzymes (*9*). We find that DDX54 is incorporated into state C in a post-hydrolysis state and remains ADP-bound in states C-F (**Figs. 1, 4, fig. S23**).

**Fig. 4.**
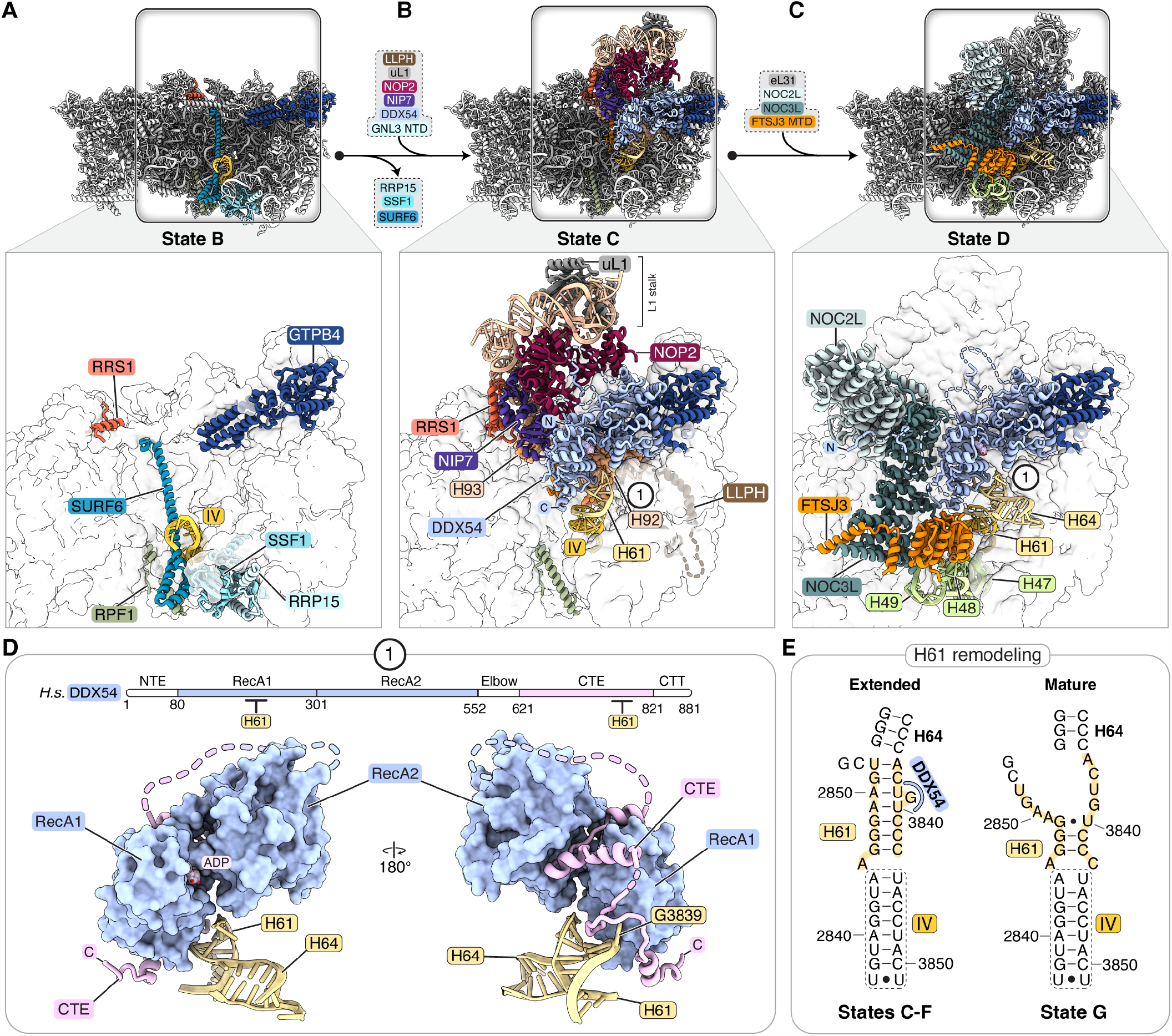
DDX54 integrates key transitions during early nucleolar assembly. (**A-C**) Crown views of states B (A), C (B) and D (C) showing selected assembly factors as they enter the respective states. Zoomed views below provide more detailed views of the environment that is present before (A), during (B) and after the installation of DDX54 (C). (**D**) A schematic indicates the domain organization of DDX54 and interactions with helix 61 (H61) of the 28*S* rRNA. The cartoon representation of DDX54 below shows two views of its interactions with a remodeled H61. A flipped-out base (G3839) is indicated. (**E**) Schematics of remodeled H61 as seen in states C-F (left) and the mature form of H61 as seen in state G and subsequent states (right).

Our structures illustrate that DDX54 has a pivotal role during nucleolar large ribosomal subunit assembly by performing three essential functions: first, DDX54 is responsible for the formation of functional centers such as the PTC and the L1 stalk. Second, DDX54 provides directionality to 60*S* maturation through remodeling of the pre-rRNA root helices such that early assembly factors can dissociate. Third, DDX54 acts as a recruitment platform for new assembly factors that catalyze downstream events (**Fig. 4, fig. S23**).

In its first role during the formation of functional centers, the association of DDX54 with GTPB4 limits conformational freedom of the remaining elements of the PTC (H90,92,93), and RNA elements that accompany the NOP2/NIP7 complex, including the L1 stalk and part of the E site. By acting as a clamp, DDX54 is thus able to facilitate PTC formation, remaining bound on top of H90 and H92, and fixing the L1 stalk in an immature conformation, consistent with prior biochemical data (*38*) (**Fig. 4A,B, fig. S23E**). The installation of elements of the PTC is therefore catalyzed through checkpoints that are controlled by energy-consuming enzymes in two steps with GTPB4 installing H89 and H91 and DDX54 stabilizing H90, 92 and 93.

In its second role, DDX54 remodels the root helices, first of domain IV (H61) and subsequently domain III (H47), triggering a series of irreversible conformational changes that provide directionality to 60*S* assembly (**Fig. 4A,B, fig. S23D**). In state B, root helices of domains III and IV are trapped by the SURF6-RRP15-SSF1 complex (**Fig. 4A**). In state C, DDX54 has successfully remodeled H61, stabilizing an extended conformation of H61 that abolishes the binding site for the SURF6-RRP15-SSF1 complex (**Fig. 4B, fig. S23D**). The unidirectionality of the removal of these proteins is further guaranteed by a FTSJ3 C-terminal segment which binds the site previously occupied by SURF6, thereby preventing re-association of the states A/B-specific SURF6-RRP15-SSF1 complex whilst additionally blocking the nascent polypeptide exit tunnel together with RPF1 (**Fig. 4A,B, fig. S23D**). As a result of the release of the SURF6-RRP15-SSF1 complex, which previously maintained domainVI in an immature conformation, a compaction occurs between domains V and VI, creating a binding site for LLPH at the solvent side (**Fig. 4B**). The remodeling of H61 further involves alternate base pairing in the extension of the root helix of domain IV, thereby repositioning H64 and accommodating the newly integrated PTC elements H90 and H92. The specific alternate base pairing on top of H61 is further interrogated by DDX54 via a flipped-out nucleotide (G3839) so that only the correct substrate can be accommodated near the PTC (**Fig. 4D, E, fig. S23E**).

In its third role as a recruitment platform, the C- and N-terminal extensions of DDX54 selectively recruit NOP2-NIP7 in state C and the FTSJ3-NOC2L-NOC3L assembly factor complex in state D, respectively (**Fig. 4B,C, fig. S23E,F**). In state D, the NOC2L-NOC3L complex is positioned in a downward orientation, thereby preventing premature maturation of domain III (**Figs. 1, 4C**).

The ability of DDX54 to interact with NOC2L-NOC3L and the methyltransferase domain (MTD) of FTSJ3 via the DDX54 C-terminal tail (CTT), allows DDX54 to organize elements of domain III that include the domain III root helix H47 as well as helices H48 and H49 (**Fig. 4C, fig. S23E,F**). In this context, the MTD of FTSJ3 performs a secondary structural role while its substrate for 2’-O-methylation (nucleotide 4499 within the A-loop/H92) is already deeply buried underneath DDX54 (**fig. S23E**). Since the position of H92 does not change after state C to expose nucleotide 4499, we believe that FTSJ3 modifies nucleotide 4499 before the PTC is stabilized by DDX54 in state C. The ability to tether FTSJ3 already in state A further supports this model (**Fig. 2B**). The structural changes observed during the transitions from state B to D highlight how an ensemble of factors centered around the DEAD-box RNA helicase DDX54 remodels important segments of pre-rRNA. Beyond enzymes such as DDX54 and FTSJ3 that directly act upon a substrate, these also include proteins that fulfil either architectural roles such as NOC2L-NOC3L or stabilizing roles such as the N-terminus of GNL3 that stabilizes ES13 in state C specifically (**Fig. 4, fig. S22C**). These findings support a model for the cooperative actions of these factors that is required to advance 60*S* assembly, further rationalizing prior genetic interactions observed in yeast among Dbp10, Noc3 and Nug1 (*39*)(**Fig. 4**).

### Transitions towards nuclear maturation

Following a rotation of the NOC2L-NOC3L complex from a downward (state D) into an upward position that is accompanied by the removal of MAK16-assembly factors at the solvent exposed side (state E; **Fig. 1, fig. S24**), we observe that a trio of WD40 repeat proteins (BOP1, WDR12 and WDR55 (yeast Erb1, Ytm1, and Jip5)) chaperone the formation of domain III in state F (**figs. S24,25**). Subsequently, the transition from state F to G is marked by the MDN1-mediated removal of a nucleolar protein interaction network. In state G, DDX54 is replaced by a peptide of the GTPase NOG2 that binds to GTPB4 (**Fig. 1, fig. S26**). The initial tethering of NOG2 is followed by the installation of the 5*S* RNP in a non-rotated state in state H, for which we have obtained a partial reconstruction (**Fig. 1, fig. S26**). For a more detailed description of these transitions, see **Supplementary Text**.

### The rixosome and NOP53 couple 5*S* RNP rotation to ITS2 processing

Following nucleolar assembly of the human pre-60*S*, a critical event during the subsequent nuclear maturation of pre-60*S* precursors is the MDN1-catalyzed rotation of the 5*S* RNP from the unrotated to the rotated state, which results in the installation of the central protuberance (CP) (*7, 15, 16, 18, 40*–*43*).

The structure of state I shows that the formation of this intermediate, in which the 5*S* RNP is rotated, is the product of several hierarchical and coupled steps (**Fig. 5**). Only upon correct rotation of the 5*S* RNP and the departure of early assembly factors (RPF2 and RRS1) can the assembly factor SDA1 and the rixosome stably associate, thereby stabilizing the L1 stalk in a bent conformation (**Fig. 5**). Association of the rixosome is required for ITS2 processing on two levels, as the rixosome-associated endonuclease LAS1L (*16, 22*) initiates cleavage of ITS2 whilst the bent conformation of the L1 stalk – induced by rixosome stabilization – serves as a binding site for a peptide of NOP53 that brings the RNA exosome into close proximity to its ITS2 substrate (**Fig. 5, fig. S27**). We note that while components of the RNA exosome and the rixosome (MDN1, SENP3, NOL9) are present in our mass spectrometry data, they could not be visualized in our reconstructions, most likely due to flexibility (**Data S1**).

**Fig. 5.**
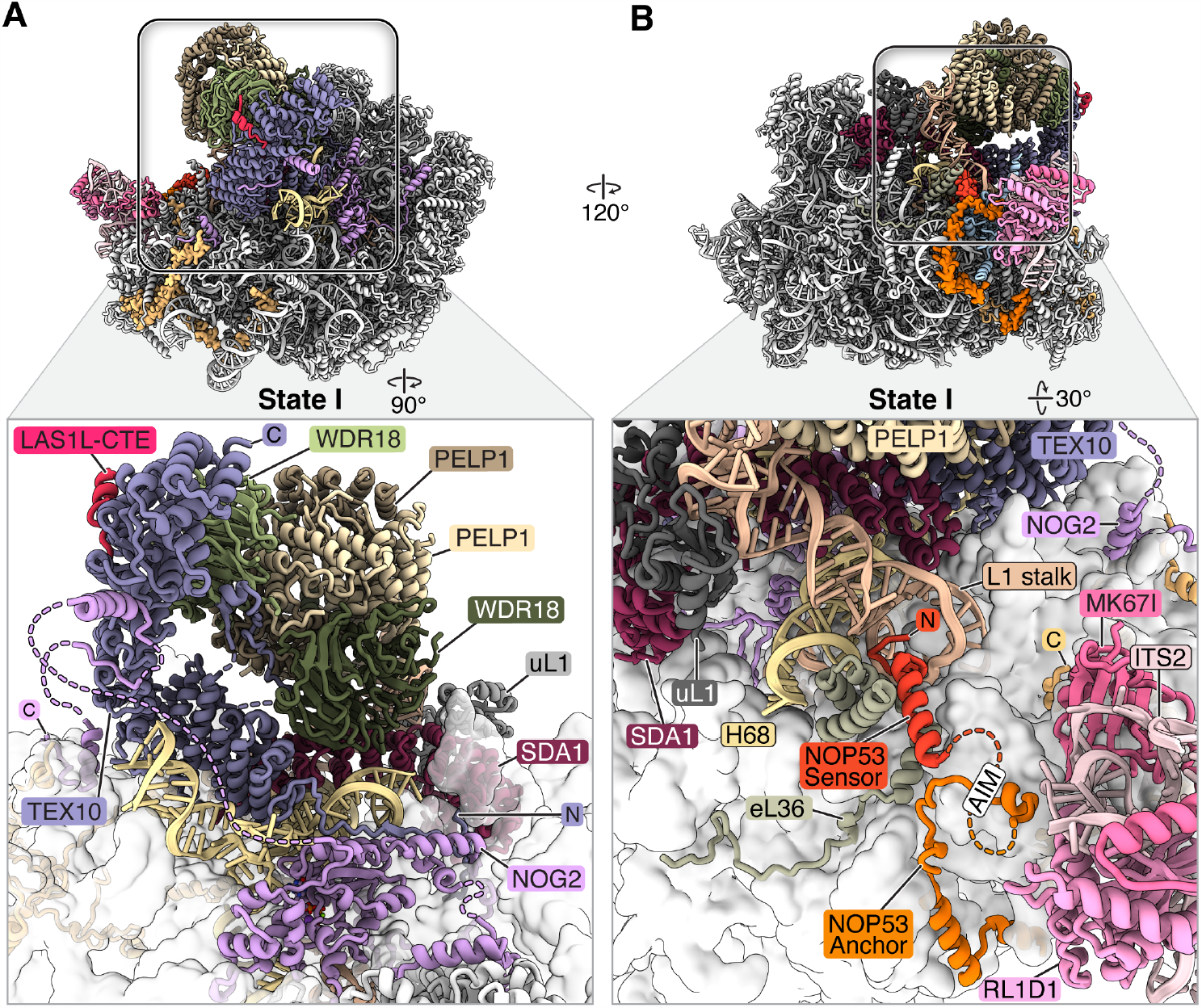
State-specific processing of ITS2. (**A, B**) Two views of state I showing selected assembly factors involved in ITS2 processing. Lower panels show a zoomed view of the rixosome and its interactions with the pre-60*S* particle (A) and the NOP53 N-terminus near the L1 stalk (B). The NOP53 sensor and anchor domains (red and orange respectively) are shown in proximity to the AIM motif.

While 5*S* RNP rotation and bending of the L1 stalk have been observed in the yeast system, limiting resolution near ITS2 has so far precluded a mechanistic understanding of how these large-scale conformational changes are coupled to processing of ITS2 by the RNA exosome (*7, 17, 40*). Our high-resolution structures of state I explain how the bent conformation of the L1 stalk and the presence of ITS2 are specifically interrogated by 200 amino acids of a universally conserved N-terminal segment of NOP53 (sensor and anchor domains), which had not previously been visualized (**Fig. 5B, fig. S27**). The involvement of the NOP53 sensor domain is supported by biochemical data showing that N-terminal truncations of this domain result in ITS2 processing defects (*44*).

The NOP53 sensor domain is only able to bind the L1 stalk in the bent conformation; binding in such a way that, in the presence of ITS2 and associated assembly factors, the NOP53 anchor domain can additionally wrap around ITS2. The joint binding of sensor and anchor domains positions a recruitment peptide of the RNA exosome, the NOP53 AIM motif (*45*), in very close proximity to ITS2. As the AIM motif has been shown to interact with the RNA exosome-associated RNA helicase MTR4 (*46, 47*), our structural data now provide key insights into the coupling of 5*S* RNP rotation, L1 stalk conformation and ITS2 processing.

The structure of state I now rationalizes how defects or depletions of assembly factors involved in 5*S* RNP rotation and the installation of the rixosome result in the accumulation of unprocessed pre-rRNA species (*18*–*22*). In the absence of a stabilized bent L1 stalk either no cleavage of ITS2 occurs, or in cases where ITS2 cleavage does occur, the ITS2 precursor cannot be efficiently processed by the RNA exosome as it is not positioned close to its substrate by NOP53 (*28*). This model also explains how the NOP53 anchor domain can remain associated with the ITS2 region during exosome-mediated ITS2 processing (*17*).

### Functional integration of the human rixosome

State I further illustrates how the human rixosome and NOG2 function through an evolutionarily expanded protein-protein interaction network that is formed through C-terminal extensions (CTEs) of assembly factors NOG2, TEX10 and LAS1L that have no counterparts in yeast (**Fig. 5A, fig. S28**). These CTEs facilitate three integrated functions during nuclear pre-60*S* maturation.

First, extensive interactions of CTEs with NOG2 allow the human rixosome to already associate with late nucleolar assembly via a peptide of NOG2 as revealed in states G and H (**Fig. 1, fig. S26**) (*10*). Second, the CTE of NOG2 is also ensuring directionality during the transition from nucleolar to nuclear assembly by probing the environment around eL34 where it has a mutually exclusive binding site with the nucleolar assembly factor FTSJ3 (yeast Spb1) (**Fig. 5A, figs. S21B, S28D)**. Third, the large C-terminal extension of TEX10 provides binding sites for NOG2 as well as several rixosome components known as the RIX1 subcomplex (comprising two copies each of PELP1 and WDR18) (*48*) as well as the C-terminal extension of the endonuclease LAS1L, thereby directly recruiting the ITS2 endonuclease to state I (**Fig. 5A, fig. S28D**). In the context of the human rixosome, CTEs are used to flexibly tether different enzymatic activities with LAS1L, NOL9, and SENP3 acting as functional modules with endonuclease, 5’ polynucleotide kinase and SUMO-specific protease activity, respectively (**fig. S28C,D**) (*27, 48, 49*).

Together, our structural data on state I highlight specific aspects of human ribosome assembly where the rixosome has evolved into a higher order protein interaction hub that directly couples 5*S* RNP rotation and L1 stalk bending to the recruitment of the endonuclease LASL1 to initiate cleavage of ITS2 for subsequent exosome-mediated RNA processing (**Fig. 5, fig. S27,28**).

### Processing of recombinant pre-ribosomal RNA in human cells

In comparison with yeast, human ITS2 is five-fold larger. While available biochemical data suggests that there are more pre-rRNA processing sites (*50*), it remains unclear how ITS2 processing is achieved and which of its RNA elements are essential for cleavage and ribosome assembly. Similarly, it is unclear if large RNA expansion segments of the human 28*S* rRNA are also required for ribosome assembly. By combining our structural data with a human large ribosomal subunit assembly assay, we have determined the order of ITS2 removal and identified key regions of pre-ribosomal RNA that are required for LSU maturation (**Fig. 6**).

**Fig. 6.**
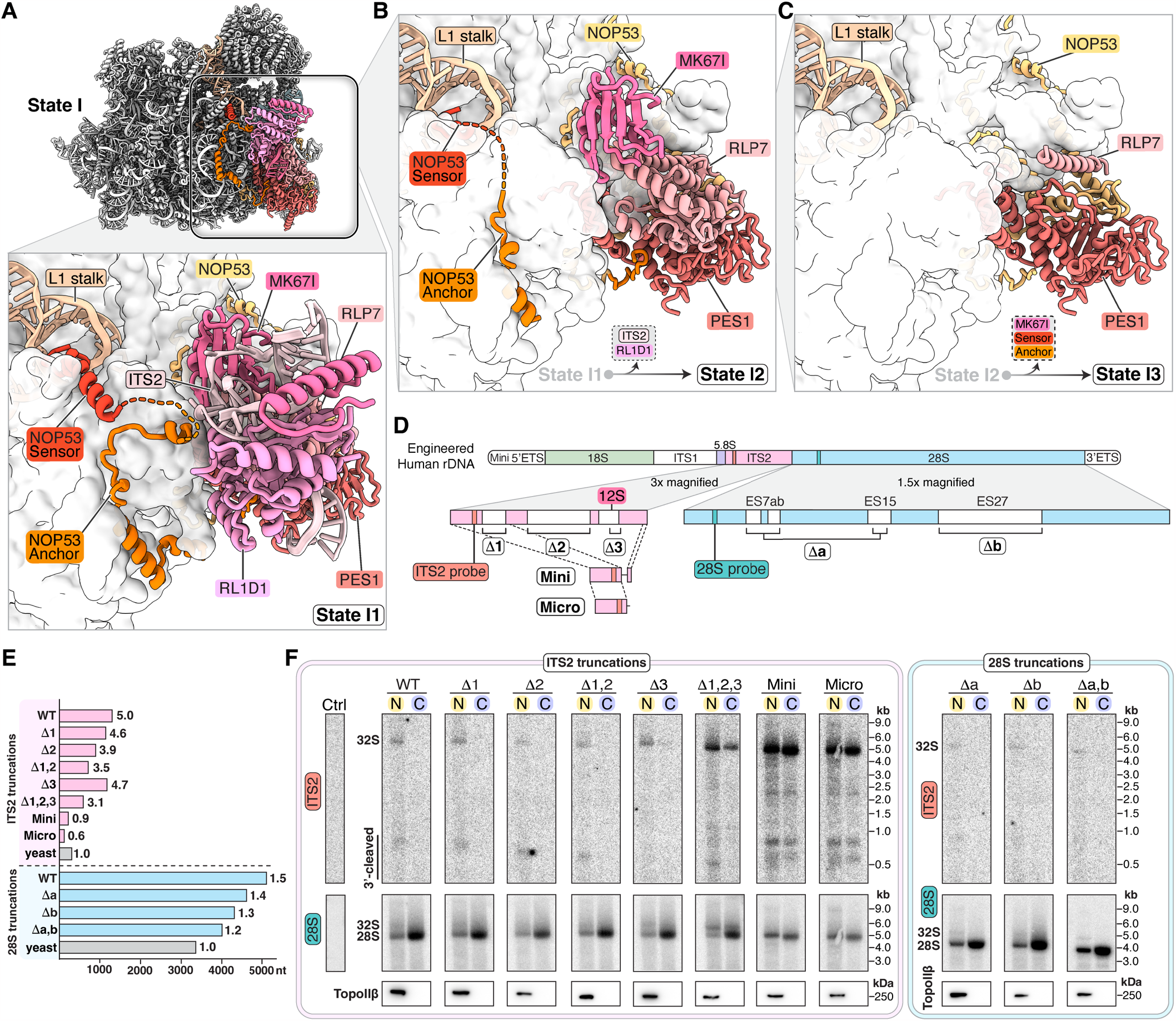
Processing of synthetic human 28*S in vivo*. (**A**) View of state I with color coded assembly factors near ITS2. The zoomed view below shows domains of NOP53 around ITS2-associated assembly factors and ITS2 (light pink). (**B,C**) Comparative zoomed views of the ITS2 region in sub-states I2 and I3. (**D**) Schematic representation of designed ITS2 variants in relation to an engineered human rDNA locus. Zoomed sections indicate the position of ITS2 variants (bottom left) and designed truncations of expansion segments ES7ab & ES15 (Da) or ES27 (Db) (bottom right). The locations of specific probes for ITS2 and 28*S* are highlighted in light red and cyan, respectively. (**E**) Bar graph indicating the size of human ITS2 (pink, top) and 28*S* (light blue) variants relative to their yeast counterparts (grey). (**F**) Northern blots of indicated synthetic human ribosomes containing truncations in ITS2 (left) and 28*S* (right) using probes for ITS2 (red, top) and 28*S* (cyan, bottom) for both nuclear (N, yellow) and cytoplasmic (C, blue) fractions. Western blots for TopoII β highlight the separation of nuclear and cytoplasmic extracts (bottom).

Similar to what has been observed for the 5’ external transcribed spacer in the context of the human SSU processome (*51*), only a very small portion of ITS2 can be visualized in all pre-60*S* assembly intermediates (**fig. S29**). In state I1, only helices 1 and 2 of ITS2 are visible, and are positioned close to the ITS2-associated assembly factors MK67I, RLP7, PES1 and RL1D1 (yeast Nop15, Rlp7, Nop7 and Cic1, respectively) (**Fig. 6A, fig. S29**). States I1-I3 represent different stages of ITS2 processing and disassembly of the associated protein factors with elements of ITS2, RL1D1 and the NOP53 anchor domain disappearing in state I2 (**Fig. 6B**). The transition from state I2 to I3 is marked by the disordering of the remaining N-terminal segments of NOP53 (sensor and anchor domains) and of the structured domain of RLP7 that is associated with MK67I (**Fig. 6B,C**).

The minimal visualized region of ITS2 within the pre-60*S* particles prompted us to investigate if disordered elements of ITS2 and expansion segments within human 28*S* rRNA are required for ribosome assembly and pre-ribosomal RNA processing by designing truncations in these regions and testing for successful ribosome assembly (**Fig. 6D-F, fig. S29B,C**). While truncations of ITS2 were based on its predicted secondary structure (**fig. S29B)**, 28*S* rRNA truncations were focused on large expansion segments that were not visualized in any of our human pre-60*S* assembly intermediates (**Fig. 6D**). All truncations were inserted into a plasmid containing an engineered human rDNA that, in addition to the indicated truncations, includes unique 28*S* and ITS2 probes for Northern blotting, specifically detecting the formation of mature recombinant large ribosomal subunits as well as its ITS2-containing precursors (**Fig. 6D, fig. S29B,C**).

Within ITS2, long helices were selected for truncations that either lacked known cleavage sites (Δ1 and Δ2) or contained the predicted LAS1L cleavage site responsible for generating a 12*S* species (*52*) (Δ3) as well as combinations of these (Δ1,2 & Δ1,2,3). We further designed ITS2 variants that lacked the LAS1L cleavage site and contained RNA segments just exceeding what is visible in our structures (Mini, Micro, **Fig. 6D, fig. S29B,C**). Within the 28*S*, rRNA truncations were designed to eliminate disordered expansion segments ES7ab and ES15 (Δa), ES27 (Δb) or all three of these (Δa,b). The resulting truncations reduced ITS2 from five times the length of yeast ITS2 to approximately 60% of the yeast ITS2 length. Similarly, the combined truncations of expansion segments of the 28*S* rRNA resulted in a 28*S* rRNA that is only 20% larger than its yeast counterpart (**Fig. 6E**).

To investigate the effects of these truncations, the designed variants were transfected into HEK293F cells from which cytoplasmic and nuclear extracts were obtained to identify engineered mature ribosomes that were exported into the cytoplasm as well as their corresponding nucleolar and nuclear precursors (**Fig. 6F, fig. S30**). Interestingly, ITS2 truncations that removed long helices without the predicted LAS1L cleavage site (Δ1, Δ2 and Δ1,2) were processed similarly to the wild-type without significant accumulation of a nucleolar precursor species (32*S*). By contrast, the elimination of the predicted LAS1L cleavage site (Δ3 or Δ1,2,3) resulted in the accumulation of a 32*S* species in the nucleus. Despite this, mature ribosomes containing either 28*S* or 32*S* pre-rRNA were detected in the cytoplasm for both Δ3 or Δ1,2,3 truncations. These data suggest that either redundant enzymes or cleavage sites exist to trigger ITS2 removal and that despite the absence of a regular LAS1L cleavage site, some precursors are exported. These observations are in line with what has been observed in yeast in the absence of Las1 where ribosomes containing ITS2 are exported into the cytoplasm and turned over (*53*–*55*) (**Fig. 6F**). The strongest accumulation of precursors was observed with minimal variants of ITS2 (Mini and Micro truncations) for which no processing of ITS2 was observed and the resulting cytoplasmic ribosomes contained ITS2 (**Fig. 6F**). For truncations of expansion segments within the 28*S* pre-rRNA, no processing defects were observed when compared to the wild-type (**Fig. 6F**). Therefore, large parts of ITS2 and the 28*S* expansion segments ES7ab, ES15 and ES27 are dispensable for ribosome assembly in *cis* in human cells, with the exception of the regions of ITS2 including the pre-60*S* associated helices and the LAS1L cleavage site whose removal results in partial processing defects.

### Transitions preceding nuclear export

Following the departure of the human rixosome, large-scale RNA conformational changes together with eukaryote-specific rRNA and protein elements drive the maturation of the central protuberance and the formation of the human E-site (**figs. S31,32**). Subsequently, we observe how the late nuclear assembly factor TMA16 together with the newly identified assembly factor L10K triggers the removal of PTC-proximal assembly factors such as GTPB4, NOG2 and NSA2 (**fig. S33**). For a more detailed description of these transitions, see **Supplementary Text**.

### Chemical modifications of pre-RNA during human pre-60*S* assembly

Beyond our insights into the role of numerous human nuclear ribosome assembly factors, the high resolution of our cryo-EM reconstructions of nuclear pre-60*S* assembly intermediates (states I-L) also allowed us to introduce and confirm the positions of many previously mapped covalent modifications of the human 28*S* rRNA (*65*).

These include modifications introduced through proteins that are guided by snoRNAs base-pairing with RNA regions surrounding their substrates (2’-O-methylations and pseudouridylations) as well as modifications introduced via protein factors directly (m^1^A, m^3^U, m^5^C, and m^6^A nucleobase modifications and a 2’-*O*-methylation) (*66, 67*). We note that during the assembly of the human pre-60*S*, the introduction of every chemical modification will require local chaperoning of substrate RNA so that any introduced modification is a direct representation of a prior RNA chaperoning event. By mapping the observed chemical modifications onto key nucleolar (states A-H) and nuclear assembly intermediates (states I-L) of our human pre-60*S* structures, we can now observe the order in which RNA elements appear and place chemical modifications in the context of ribosome assembly (**Fig. 7**). This analysis highlights four central concepts for human pre-*60S* assembly:

**Fig. 7.**
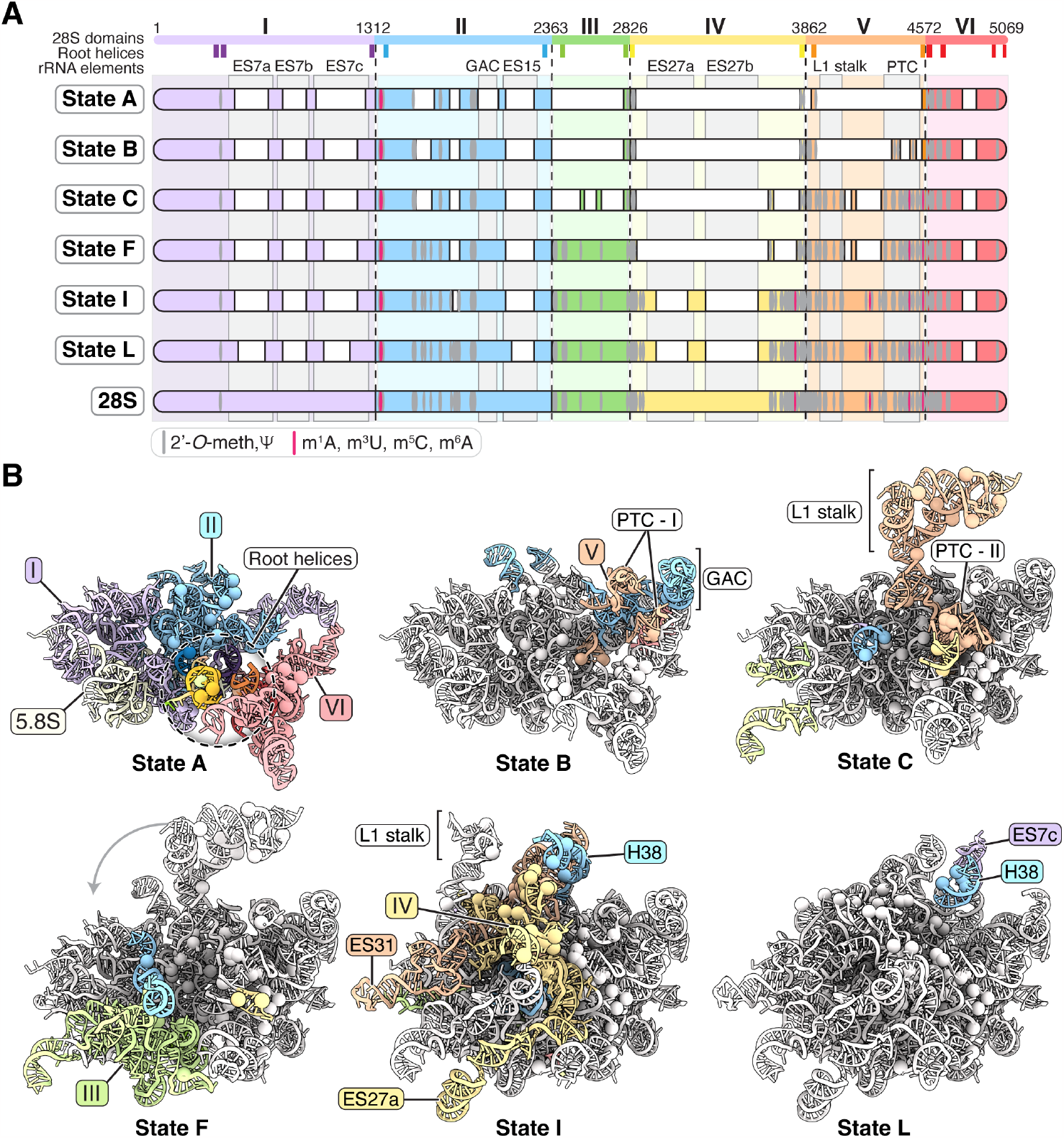
Chemical modifications of human 28*S* rRNA during assembly. **(A)** Schematic domain organization of the six color-coded domains (I-VI) of the human 28*S* rRNA. Root helices are indicated as darker blocks. Below the schematic, visible RNA elements are indicated for different states of the human pre-60*S* assembly pathway in the nucleolus (states A, B, C, F) and nucleus (states I, L). Chemical modifications are shown in grey or red as indicated. The positions of selected functional centers (PTC, GAC, L1 stalk) and RNA expansion segments (ES7, ES15, and ES27) are shown. (**B**) RNA structures of assembly intermediates shown in (A) with newly folded RNA elements color coded. Root helices and functional centers (GAC, PTC, L1 stalk) are indicated. Chemical modifications are represented as spheres.

First, during nucleolar stages, domains I-VI assemble in an order similar to what has been previously observed in yeast (*11*–*13*) with domains I, II and VI folding first before domains V, III and IV are integrated (**Fig. 7A**). Second, already in state A, all root helices, the origins of each of the six subdomains of the 28*S* rRNA are visible. Importantly, a high density of chemical modifications is observed near or within these root helices, suggesting significant prior RNA chaperoning and hence highly controlled RNA folding before the formation of state A (**Fig. 7A,B**). Third, in line with the notion that the presence of a chemical modification requires prior RNA chaperoning, we find that regions that are heavily modified form late during assembly, such as elements of domains V and These regions include the PTC that is installed in two steps via the GTPase GTPB4 in state B (PTC-I, **Fig. 7B**) and the DEAD-box RNA helicase DDX54 in state C (PTC-II, **Fig. 7B**). In the context of ribosome assembly, the presence of a modified nucleotide can therefore be seen as a means of either delaying folding in a particular region or preventing misfolding. This logic further implies that regions that need to fold quickly at the beginning of assembly, such as domain I, should have a low density of chemical modifications, which is indeed what we observe. Fourth, regions that are not conserved, disordered in our reconstructions, or not subject to extensive RNA remodeling, should lack chemical modifications. This is precisely what we observe for expansion segments such as ES7, ES15 and ES27, which our functional studies show to be dispensable for ribosome assembly (**Figs. 6, 7A**).

Together, the mapping of human chemical RNA modifications as a function of ribosome assembly in the nucleolus and nucleus provides a foundation to study both universally conserved chemical modifications as well as species-specific adaptations.

## Discussion

During nucleolar and nuclear stages of pre-60*S* assembly, functional centers need to be first assembled and subsequently matured. Based on our structural data, we now propose a simplified model for human nucleolar and nuclear pre-60*S* biogenesis (**Fig. 8**). We highlight that emerging key nucleolar themes include high-level organization and hierarchy at the pre-rRNA and protein level (**Figs. 2,8)**, the use of energy-consuming enzymes such as GTPB4 and DDX54 to drive transitions and the installation of functional centers in a unidirectional manner (**Figs. 3,4,8**). Critical nuclear themes include recognition of pre-ribosomal RNA for LAS1L-mediated cleavage followed by RNA exosome-mediated processing of ITS2 and how the human rixosome can integrate these processing steps through C-terminal protein extensions (**Figs. 5,8**).

**Fig. 8.**
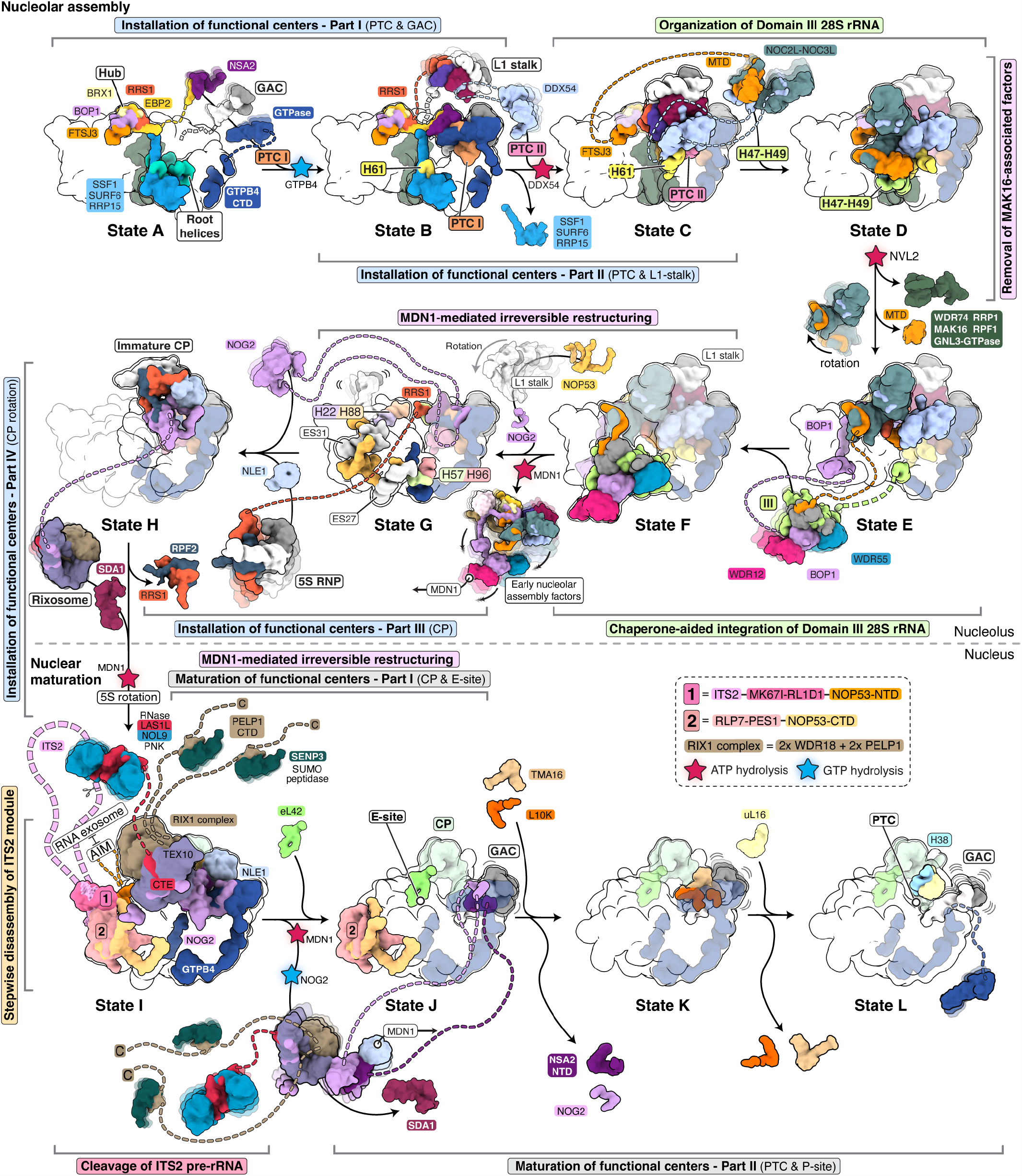
Model for human pre-60*S* biogenesis in the nucleolus and nucleus. Simplified schematic model of human large ribosomal subunit biogenesis in the nucleolus (top) and nucleus (bottom). Key nucleolar transitions such as the installation of functional centers (PTC, GAC, L1-stalk, and CP; blue highlights), the domain III integration (green highlights) and the irreversible steps involving protein removal (pink highlights) are indicated. In the nucleus, critical maturation steps such as ITS2 cleavage by LAS1L (red highlight), subsequent stepwise disassembly of ITS2 module by the RNA exosome (orange highlight) and maturation of functional centers (CP, E-site, PTC and P-site; grey highlight) are indicated. Irreversible GTP and ATP hydrolysis events are indicated with blue and red stars, respectively. Color-coded tethered protein and RNA elements are indicated with linkers and motion blur.

Beyond representing a near-complete structural inventory of nucleolar and nuclear human pre-60*S* assembly, our data now start to interweave human ribosome assembly with research fields that have so far appeared disconnected at a mechanistic level. The observation of post-translational modifications of ribosome assembly factors as observed for BOP1 (**Fig. 2, fig. S21B**) suggest a direct mechanistic role in the molecular control of human ribosome assembly and indicates how signaling pathways may integrate human ribosome assembly with cellular metabolism. In addition, our work lays the foundation for a mechanistic understanding of human diseases so that the molecular effects of point mutations in ribosomal proteins can be put into a temporal context of large ribosomal subunit assembly in patients suffering from a range of ribosomopathies (*68*–*70*).

At the nuclear level, the structural basis for the coupling between 5*S* RNP rotation and ITS2 processing as shown in state I (**Fig. 5**) highlights the importance of structural studies with endogenous complexes. Similarly, it underlines the complexity of eukaryotic ribosome assembly in which MDN1-induced 5*S* RNP rotation precedes ITS2 processing with Las1 cleavage and Nop53-mediated recruitment of the RNA exosome, events that cannot be reconstituted by using only enzymes involved in ITS2 processing (*17, 71*).

Across nucleolar and nuclear assembly, our analysis of chemical modifications of RNA indicates that chemical modifications can be used as a readout of prior RNA chaperoning during the assembly of human pre-60*S* subunits (**Fig. 7)**. Importantly, this notion is in agreement with additional functions that have been described for chemical modifications: red,

First, chemical modifications in yeast and human cells may also serve as checkpoints during assembly where they may be interrogated by assembly factors (**fig. S28B**) (*10*). Second, while chemical modifications can result in additional stabilization of the ribosome, complete ribosomes can be formed even in the presence of catalytically dead enzymes involved in pseudouridylation while the conformational freedom of subunits including inter-subunit rotation is partially affected (*72*). Third, the heterogeneity of chemical modifications as observed in the human ribosome (*73*) can be explained as the result of several redundant sites of chemical modifications that together guarantee efficient ribosome assembly without the need for selective and quantitative modification. These data suggest that in archaeal and eukaryotic systems in parallel to proteinaceous assembly factors, snoRNA-mediated modifications have evolved as a second layer of control to ensure the productive biogenesis of ribosomes.

## Materials and Methods

### MK67I-GFP^+/+^ cell line generation

A biallelically tagged MK67I-GFP cell line was generated using an in-house genome editing platform called SNEAK PEEC (*74*) (**fig. S1A**). Briefly, we designed two DNA repair templates, each encoding a dual affinity tag (TEV-cleavable HA, 3C-cleavable GFP) followed by a surface display sequence encoding a unique protein epitope (the BtuF vitamin B12 binding protein or the capsid protein p24 from HIV1)(*75, 76*) and flanked by two homology arms covering 600 bp in either direction of the Cas9 cut site in the last exon of the *NIFK* gene (coding for the MK67I protein). Both DNA repair templates were cloned into a pUC57 vector for transfection purposes. The 20-bp sgRNA target sequence (5’-GGAGAGGCGAAAATCTCAAG-3’, PAM: TGG) was cloned into a third plasmid expressing the single guide RNA (sgRNA) and the high specificity *S. pyogenes* Cas9 variant (eSpCas9(1.1)). This plasmid is a gift from Feng Zhang (*77*) (Addgene #71814). The sgRNA sequence was selected using crispor.telfor.net (*78*). Three μg of total DNA, split in equimolar concentrations of each plasmid (Repair template 1 [Btuf], Repair template 2 [p24], Cas9 + sgRNA) were reverse transfected into human HEK 293-F cells (ThermoFisher Scientific, R79007) using Lipofectamine 3000 reagent (ThermoFisher Scientific, L3000015) (1 × 10^6^ cells/ transfection/well) in 1 ml of Freestyle™ 293 Expression Medium (ThermoFisher Scientific, 12338026) supplemented with 2% heat inactivated FBS (ThermoFisher Scientific) in a 6-well plate (VWR, 10062-892). Cells were allowed to recover 24h, after which they were expanded to three new wells on the same 6-well plate for at least 5 days. Cells were harvested from the 6-well plate by gentle aspiration and stained with fluorescently labeled nanobodies (Alexa 646, APcY7), each of which specifically binding to one of the surface display epitopes. Fluorescence-assisted cell sorting (FACS) was used to first select cells expressing GFP, followed by selection of cells within this population that stained positive for both btuF and p24 surface display epitopes (**fig. S1B**). Sorted single cell clones were expanded for two weeks and then PCR-genotyped to confirm biallelic editing. Extraction of human genomic DNA was carried out using the QuickExtract DNA Extraction Solution (Lucigen, QE09050) following the standard protocol. Thirty μl of extracted genomic solution was used per screening PCR reaction (50 μl final volume). The following PCR primers were used to detect the biallelic integration of the tag:

NIFK-genomic-fwd (CAGGATTCTGTGAATCAGTGAGCTCCAGGTTTTGGCC), p24-rev (TGCTGTCATCATTTCCTCGAGCGTAGCACC), btuf-rev (GACTCCACGGGGCC AACTGTCTCAAGG).

The selected cell line was positive for both btuF and p24 surface display integration, indicating a biallelic tagging of the assembly factor MK67I with GFP (**fig. S1C,D**).

### Purification of human pre-60S assembly intermediates

Purification of human pre-60*S* assembly intermediates was carried out by sequential fractionation (**fig. S1E**). MK67I-GFP^+/+^ cells were grown as suspension cultures in Freestyle™ 293 Expression Medium. When reaching a concentration between 4-5 × 10^6^ cells/ml, cells were harvested, washed twice with ice-cold PBS and frozen into liquid nitrogen as droplets for storage at -80°C until use. Frozen cell pellets were resuspended in Buffer A (25 mM Hepes pH 7.6, 65 mM NaCl, 65 mM KCl, 3 mM MgCl_2_, 1 mM EDTA, 10 % Glycerol, 0.05 % Triton-X, 1 mM DTT, 0.5 mM PMSF, 1 mM Pepstatin, 1 mM E-64 protease inhibitor) and dounced (pestle A) 7 times on ice. The resulting lysate was centrifuged, and the supernatant discarded. The pellet was washed twice with Buffer A. The intact nuclei were then resuspended in Buffer B (25 mM Hepes pH 7.6, 10 mM KCl, 10 mM MgCl_2_, 1 mM CaCl_2_, 1 mM EDTA, 100 mM Arginine, 25 mM ATP, 1 mM Spermidine, 5 % Glycerol, 0.1 % Triton, 1 mM DTT, 0.5 mM PMSF, 1 mM Pepstatin, 1 mM E-64 protease inhibitor, 800 U RNase-free DNase I) and incubated 30 minutes on a nutator at 4°C. The insoluble fraction was removed by centrifugation at 4 °C, 25,000 g for 20 minutes. The supernatant was incubated with anti-GFP nanobody beads (Chromotek) for 4 hours at 4°C and the beads were washed 2X with Wash Buffer 1 (25 mM Hepes pH 7.6, 75 mM NaCl, 75 mM KCl, 5 mM MgCl_2_, 0.5 mM EDTA, 100 mM Arginine, 2.5 % Glycerol, 0.1 % Triton, 1 mM DTT, 0.5 mM PMSF, 1 mM Pepstatin, 1 mM E-64) and 1X with Wash Buffer 2 (25 mM Hepes pH 7.6, 75 mM NaCl, 75 mM KCl, 5 mM MgCl_2_, 0.5 mM EDTA, 100 mM Arginine, 2.5 % Glycerol, 0.05 % C12E8 (Anatrace, O330)). The complexes were eluted using 3C-protease cleavage (**fig. S1F**) and centrifuged at 25,000 g for 30 minutes to remove aggregates, immediately followed by preparation of cryo-EM grids or northern blot and SDS-PAGE analysis (**fig. S1G-I, S2**).

### Recombinant human rDNA engineering

A full copy of the human rDNA sequence was cloned into an in-house modified pUC57 plasmid (*51*). The rDNA locus includes the Pol I promoter, 5’ETS, 18*S*, ITS1, 5.8*S*, ITS2, 28*S*, 3’ETS and a shorter terminator comprising only 5 termination signals. Unique sequences were introduced into the second stem loop of ITS2 and in an expansion segment of the large subunit (ES5L). These unique sequences are not recognized within the human transcriptome and can be used to monitor the production of recombinant ribosomes in human cells transfected with the rDNA plasmid by Northern blot. To assess the requirement of different regions of the human ITS2 and expansion segments in the maturation of the large ribosomal subunit, we performed structure-based truncations into specific regions of ITS2 and 28*S* that are not immediately visible within the pre-60*S* particles (**Fig. 6 and fig. S29**). We first removed the third stem loop (Δ1: 128-228), the 5^th^ stem loop (Δ2: 411-699) and the 6^th^ stem loop (Δ3: 763-842) or combinations of these deletions (Δ1,2 and Δ1,2,3). Then, we designed a mini ITS2 containing stem loops 1, 2, a truncated stem loop 3 and the last 40 bases of ITS2; and a micro ITS2 containing only stem loops 1 and 2 and a linker of 20 bases. Finally, in the context of a rDNA locus without truncations in ITS2, we removed expansion segments 7a, 7b and 15 in one construct (Δa: 525-633, 766-888, 2137-2270), expansion segments 27a,b in a second construct (Δb: 2946-3283, 3009-3589) and a combination of both truncations in a third construct (Δa,b). All plasmids were fully sequenced using nanopore technology (Plasmidsaurus).

### Cell transfection for functional studies

Three μg of the engineered rDNA plasmids were reverse transfected into human HEK 293-F cells using Lipofectamine 3000 reagent (1 M cells/transfection/well in 6-well plates). The cells were incubated at 37 °C, 8% CO_2_. After 48h, the cells were washed once with cold PBS, gently detached from the plate with cold PBS and then pelleted at 100x g for 5 minutes at 4°C. The pellets were immediately used for isolation of recombinant ribosomes from different cellular compartments.

### Isolation of recombinant ribosomes from different cellular compartments

To isolate the pre-ribosomes and mature ribosomes from different cellular compartments, cells were first resuspended in hypotonic buffer (20 mM Tris-HCl pH 7.4, 10 mM NaCl, 3 mM MgCl_2_) supplemented with 1 mM PMSF, 1mM Pepstatin and 1 mM E-64 and incubated on ice for 15 minutes. NP-40 was added at a final concentration of 0.5% and cells were vortexed at max speed for 15 seconds. At this stage, most of the cells were lysed yielding intact nuclei. The lysate was centrifuged 5 minutes at 100 x g at 4°C, and the cytoplasmic fraction was kept for total RNA extraction and western blot analysis. The nuclei were washed 3 times with hypotonic buffer to prevent any contamination of cytoplasmic ribosomes in the subsequent nuclear extraction. Then, the nuclei were resuspended in RIPA buffer (20 mM Tris-HCl pH 7.4, 150 mM NaCl, 1 mM EDTA, NP-40 1%, 0.5% deoxycholate, 0.1% SDS) supplemented with 1 mM PMSF, 1mM Pepstatin and 1 mM E-64 and incubated 30 minutes on ice with vortexing at 10-minute intervals. The lysate was centrifuged 30 minutes at 20,000 x g at 4°C. The supernatant constituting the nuclear fraction was kept for total RNA extraction and western blot analysis.

### RNA extraction and Northern Blot analysis

The RNA was extracted using Trizol/chloroform and isopropanol precipitation following standard procedures. Different quantities of total RNA or rRNA were loaded and resolved on 1.2 % denaturing agarose gel as follows: 5 μg for whole cell extracts, 5 μg for intermediate purification fractions, 2 μg for nuclear fractions containing recombinant ribosomes, 4 μg for cytoplasmic fractions containing recombinant ribosomes and 50 ng for the purified pre-60*S* particles. After complete migration, the RNA was transferred to a positively charged nylon membrane and fixed by UV-crosslinking. The following 5′-^32^P-labeled oligonucleotide probes were used for Northern blot analysis:

Endogenous ITS2 probe (5’-CTGCGAGGGAACCCCCAGCCGCGCA -3’), Endogenous 28*S* probe (5’-ATTCGGCGC TGGGCTCTTCCCTG -3’), Unique ITS2 probe (5’-CATA AAGCCTCGATCCTCGGG -3’), Unique 28*S* probe (5’-TG AAGCTCTCGAGTGTACCT-3’)

The Millenium RNA marker (Thermo Fischer Scientific) was used as a molecular weight ladder.

### Western blotting

To assess whether the nuclear compartment was not leaking into the cytoplasm during cellular fractionation, we sought to analyze the protein content of the cytoplasmic and nuclear fractions using topoisomerase IIβ as a nuclear-specific marker. Equal amounts of total protein (10 μg) from nuclear or cytoplasmic extract containing the recombinant ribosomes were separated by SDS–PAGE and transferred to a PVDF membrane. The membrane was blocked in TBS-T supplemented with 5% BSA for 1 h at room temperature before incubating with primary antibody at 1:5000 dilution (mouse Topo IIβ Antibody: clone A2, cat. #sc-365071, Santa Cruz Biotechnology) in TBS-T with 3% BSA overnight at 4°C. The membrane was washed three times in TBS-T and then incubated for 45 minutes at room temperature with the horseradish peroxidase (HRP)-conjugated secondary antibody goat anti-mouse IgG (cat. #115-035-003, Jackson ImmunoResearch) diluted at 1:5000 in TBS-T with 3% BSA. The membrane was washed twice in TBS-T and once in TBS before imaging on a Typhoon 9400 imager (GE) using ECL Prime detection reagent (Amersham). The Precision Plus Protein (Biorad) was used as a molecular weight ladder.

### Cryo-EM grid preparation

Cryo-EM grids were prepared using a Vitrobot Mark IV robot (FEI Company) maintained at 10°C and 95 % humidity. The sample (3.5 μl) was applied four times to glow-discharged Quantifoil R3.5/1 grids coated with a layer of 2 nm ultrathin carbon (Au 400 2nm C, Electron Microscopy Sciences) with 45 seconds incubation followed by a manual blot between each application. After the last incubation, the grid was blotted by the Vitrobot for 10 seconds using a blot force of 8 and plunged into liquid ethane.

### Cryo-EM data collection

Imaging was carried out on a Titan Krios electron microscope (FEI) equipped with an energy filter (slit width 20 eV) and a K3 Summit detector (Gatan) operating at 300 kV with a nominal magnification of 64,000×. Using SerialEM (*79*), four datasets totaling 172,699 movies were collected with a defocus range of -0.5 to -2.5 μm and a super-resolution pixel size of 0.54 Å. Images with 40 subframes were collected using a dose of 30 electron/pixel/second (1.08 Å pixel size at the specimen) with an exposure time of 2.0 seconds and a total dose of 60 e^-^/Å^2^. A multi-shot 3-by-3 beam-tilt imaging strategy was employed to record 90 micrographs per stage position, yielding approximatively 10,000 micrographs per day.

### Cryo-EM data processing

Raw movies were on-the-fly gain-corrected, dose-weighted, aligned and binned to a pixel size of 1.08 Å using RELION’s own implementation of MotionCor2 (*80*). During import, movies were pooled in nine different optic groups for each dataset. The defocus value of each micrograph was estimated using Gctf (*81*). A total of 15,679,142 particles were picked from 4 datasets using the gaussian blob picker in RELION 4 (*82*). The particles from each dataset were extracted at a pixel size of 4.32 Å and were separately cleaned up with 2D classification in RELION 4 (VDAM algorithm) followed by heterogeneous refinement in CryoSPARC v3.3 (*83*) (**fig. S3**). For each dataset, the particles contributing to the reconstruction of a featured pre-60*S* intermediate were selected, re-imported into RELION 4.0 using pyEM (*84*) and re-extracted at a pixel size of 2.16 Å. To further remove junk particles, these particle stacks were subjected to one round of 3D classification without alignment in RELION 4.0. The particles belonging to good classes were merged, re-centered and re-extracted at a pixel size of 1.08Å. The final particle stack of each dataset was subjected to one round of CTF refinement and Bayesian polishing in RELION 4.0. The polished particles from each dataset were merged into a consensus particle stack of 2,474,972 particles. This particle stack was subjected to a 3D refinement in RELION 4.0 resulting in the reconstruction of a consensus human pre-60*S* intermediate at a global resolution of 2.17 Å. To separate all the different states present in the consensus reconstruction, extensive global and focused 3D classifications without alignment were performed in RELION 4. Twenty-four states, showing unique features that highlight the progression of the early human pre-60*S* assembly pathway, were isolated (**figs. S4-5**). To improve their resolution, two rounds of Global CTF refinement (beam tilt, aberrations and magnification anisotropy) were performed in CryoSPARC v3.3. After calibration, the pixel size was adjusted to 1.072Å/pix and the resolution of the reconstructions was recalculated. The global resolution for the 24 states ranges from 2.5 to 3.2Å (**figs. S4-5**).

To harness the full structural potential and overcome the flexibility present in each state, 3D variability analysis, particle signal subtraction and focused local refinements were performed in CryoSPARC v3.3 using different masks encompassing relevant modules (**figs. S6-15, tables S1-2**). Each focused map and their respective half maps were rescaled at 1.072Å/pix and postprocessed using either cryoSPARC or phenix.auto_sharpen (*85*). To facilitate model building and overall visualization of each state, the focused maps were combined into a composite map using the ‘vop max’ command in ChimeraX (*86*) (figs. S16, S18). The corresponding composite half maps were generated with phenix.combine_focused_maps in PHENIX (*87*) and were used to estimate the global and local resolution of the composite map for each state. Local resolution estimation and filtering for the overall, focused, and composite maps were performed with cryoSPARC v3.3. 3DFSC curves were calculated using the 3DFSC server (*88*).

### Model building and refinement

A combination of AlphaFold structure predictions (*89*), existing X-ray/EM structures, and *de novo* model building was used to build the different human pre-60*S* assembly intermediates (**tables S3-4**). Most of the human ribosomal protein atomic coordinates were extracted from the late pre-60*S* EM structure (PDB ID 6LSS) (*25*), rigid-body docked in our EM density and manually adjusted in COOT (*90*). All assembly factors were built and manually adjusted in COOT using an AlphaFold structure prediction as a starting model. The 5.8*S*, 28*S* and 5*S* rRNA were extracted from the late pre-60*S* EM structure (PDB ID 6LSS) (*25*), docked in our EM maps, morphed using prosmart restraints (*91*) and manually curated in COOT. The ITS2 rRNA was built manually *de novo*. The different rRNA were then rebuilt with optimized geometry using ERRASER (*92*). The final models for the 24 states were real-space refined in their respective composite maps using phenix.real_space_refine in PHENIX (*87*) with secondary structure restraints for proteins and RNAs. The model refinement statistics can be found in **table S5**. The maps and models were analyzed and visualized in ChimeraX (*86*) or PyMol (Schrödinger, LLC). Figures were generated using ChimeraX (*86*).

### AlphaFold-multimer prediction

The SENP3-PELP1 interaction was predicted using AlphaFold-Multimer (*89*) with the full-length amino acid sequences from Uniprot (SENP3: Q9H4L4; PELP1: Q8IZL8) as input.

### RNA secondary structure predictions

RNA secondary structure predictions were performed using LocARNA-P (*93*) and the Vienna RNA websuite (*94*).

### Sequence alignments

Sequence alignments were performed using Clustal Omega (*95*).

## Acknowledgments

We thank M. Ebrahim, J. Sotiris, and H. Ng for help with grid screening and data collection at the Evelyn Gruss Lipper Cryo-Electron Microscopy Resource Center, The Rockefeller University. We thank Søren Heissel for swift mass spectrometry analysis of our purified samples at the Proteomics Resource Center, The Rockefeller University. Molecular graphics and analyses were performed with UCSF ChimeraX, developed by the Resource for Biocomputing, Visualization, and Informatics at the University of California, San Francisco, with support from National Institutes of Health R01-GM129325 and the Office of Cyber Infrastructure and Computational Biology, National Institute of Allergy and Infectious Diseases. We thank members of the Klinge laboratory for critical reading of this manuscript.

## Author contributions

AVB and SK conceived of the study, designed the experiments and analyzed the data. AVB performed the genome editing, prepared the samples of human pre-60*S* and performed the cryo-EM analyses. AVB built the atomic models with help from SK. AVB engineered the rDNA plasmids and performed the functional assays. AVB and SK wrote and edited the manuscript.

## Data and materials availability

The raw unaligned multi-frame movies and the consensus pre-60*S* polished particles stack have been deposited in the Electron Microscopy Public Image Archive (EMPIAR-XXXXX). The composite cryo-EM maps and atomic models have been deposited in the Electron Microscopy Data Bank (EMDB) and the Protein Data Bank (PDB), respectively: state A1 (EMD-29252 and 8FKP), state A2 (EMD-29253 and 8FKQ), state B1 (EMD-29254 and 8FKR), state B2 (EMD-29255 and 8FKS), state C1 (EMD-29256 and 8FKT), state C2 (EMD-29257 and 8FKU), state D1 (EMD-29258 and 8FKV), state D2 (EMD-29259 and 8FKW), state E (EMD-29260 and 8FKX), state F (EMD-29261 and 8FKY), state G (EMD-29262 and 8FKZ), state H (EMD-29263 and 8FL0), state I1 (EMD-29265 and 8FL2), state I2 (EMD-29266 and 8FL3), state I3 (EMD-29267 and 8FL4), state J1 (EMD-29268 and 8FL6), state J2 (EMD-29269 and 8FL7), state J3 (EMD-29271 and 8FL9), state K1 (EMD-29272 and 8FLA), state K2 (EMD-29273 and 8FLB), state K3 (EMD-29274 and 8FLC), state L1 (EMD-29275 and 8FLD), state L2 (EMD-29276 and 8FLE), state L3 (EMD-29277 and 8FLF). All consensus and focused refinement maps have also been deposited in the EMDB (**tables S1-2**). Materials in the main text or the supplement are available from SK upon reasonable request.

