## Supplementary Materials for "Principles of human pre-60*S* biogenesis"

**This PDF file includes:**

Supplementary Text
Figs. S1 to S33
Tables S1 to S5
References

#### Supplementary Text

#### Transitions towards nuclear maturation

##### A trio of WD40 repeat proteins chaperone the integration of domain III

The parallel assembly pathways observed for states A to D merge once state D2 has formed and the departure of assembly factors at the interface of domains I and II enables further compaction of pre-rRNA (**figs. S16,17**). Subsequently the transition from state D2 to state E involves a rotation of the NOC2L-NOC3L complex which is essential for the subsequent formation of domain III. As a result of this rotation, parts of domain III (H48, H49) and the FTSJ3 MTD become disordered while parts of domain II (H32) and the RRS1 C-terminal region become stabilized (**fig. S24**). With the NOC2L-NOC3L complex now in an upward position, domain III assembly can occur during the state E-to-F transition (**fig. S24**). State F contains a fully chaperoned domain III that is held together via several tethered multiprotein complexes. Here, a peptide of the already bound FTSJ3 is involved in the recruitment of eL27, eL30 and eL34 while the tethered BOP1 WD40 domain recruits both WDR12 as well as WDR55, an assembly factor positioned near eL19 that has so far not been visualized in any ribosome assembly intermediate and whose function could be related to the yeast assembly factor Jip5 (**figs. S21B, 25**). Together with ITS2-bound assembly factors PES1 and RLP7, the three WD40 repeat proteins (BOP1, WDR12 and WDR55) act in concert to chaperone domain III helices 54, 57 and 59, allowing juxtaposition of key helices while preventing direct interactions that are only observed in state G (**fig. S25**). In state F, WDR55 performs a central role by stabilizing domain III (helices 47, 57, 59) and domain VI (H96) around eL19 while preventing the association of domain IV (helix 63). In state G, where MDN1-mediated removal of many BOP1 interactors, including WDR55, has taken place, helices of domains III, IV and VI have coalesced onto eL19 and helices 57 and 96 form an interdomain connection between domains III and VI.

The structural transitions observed from states F to G highlight that an ensemble of WD40 repeat proteins is not only involved in docking domain III, but also in its chaperoned integration with domains VI and IV.

##### Irreversible restructuring of late nucleolar particles primes nuclear maturation

With the installation of domain III in state F, the assembly of the solvent accessible regions of the human large ribosomal subunit is largely complete. However, the presence of many early nucleolar assembly factors prevents access for late nucleolar and early nuclear assembly factors that are involved in maturing the subunit interface, as described in detail below (**fig. S26**). As highlighted previously, many of these assembly factors are highly interconnected in the human nucleolar assembly pathway, allowing their concerted removal to vacate strategic positions for subsequent maturation (**Figs. 2-4, fig. S26**).

In state F, this network of assembly factors blocks key sites from further maturation: The binding of BOP1 near PES1 prevents ITS2 processing, the FTSJ3 C-terminal domain blocks the nascent PET from being interrogated by GTPB4, and the elbow region of DDX54 blocks access to GTPB4 for nuclear assembly factors such as NOG2 (**fig. S26A,B,E**). Similarly, inter-domain rRNA interactions are blocked so that domain I:IV (H22/H88) and domain III:VI (H57/96) interactions are prevented by ribosome assembly factors BRX1-EBP2 and WDR55 respectively. The transition from state F to state G is associated with the removal of BOP1-WDR12 and its interaction partners leading to the replacement of BOP1 with NOP53 to prime ITS2 maturation (**fig. S26B**). In state F, the N-terminus of BOP1 interacts with PES1 in the ITS2 region, contacting DDX18 and NOP16, before terminating in the protein-protein interaction hub with EBP2-BRX1-

FTSJ3. Through FTSJ3, this protein-protein interaction hub is further connected with NOC3L-NOC2L, DDX54 and the NOP2-NIP7 complex (**fig. S26A**). In an important irreversible step during nucleolar pre-60*S* assembly, mediated by the AAA+ ATPase MDN1, the inter-connected assembly factors that either directly or indirectly interact with the WDR12-BOP1-WDR55 complex in state F are removed.

In addition to a mutually exclusive binding site with BOP1 near PES1, the binding sites of NOP53 in state G also prevent re-association of DDX18 and NOP16 near domain I and FTSJ3 near eL8 and eL27 (**fig. S26D**). The re-association of FTSJ3 is further prevented, as the GTPB4 C-terminal domain now binds the nascent PET in a region close to where the FTSJ3 C-terminus was previously bound. With the removal of DDX54, the structured domains of GTPB4 are now available for engagement by a peptide of NOG2 to facilitate late nucleolar and early nuclear maturation (**fig. S26A,B,E**). In state H, of which we have only determined a partial reconstruction, the GTP-bound NOG2 and an immature Central Protuberance (CP) containing the 5*S* RNP have been integrated. Here RRS1 interconnects eL21 with regions of domain V (H81 & H86), RPF2, and the ubiquitin-like domain (UBL) of NLE1 with its N-terminus to prevent premature engagement of NLE1 with MDN1 (**fig. S26C,E**). Beyond the similarities to the yeast Nog2 particle (*16*), the visualized elements of the 5*S* RNP and associated ribosome assembly factors in state H highlight how the integration of this particle is organized from the very beginning of nucleolar assembly as state A1 already contains a peptide of RRS1 (**Fig. 2B, fig. S21D**). The expanded roles of RRS1 further highlight that proteins involved in peptide-like interactions can be functionally developed in the mammalian system.

Our structural insights into late stages of human nucleolar assembly as exemplified by states F to H show how a vast protein-protein interaction network is strategically placed to stall key transitions. The ATPase-dependent removal of these factors not only results in the replacement of many assembly factors by NOP53, but also provides a checkpoint and unidirectional progression as re-association of early assembly factors is prevented.

#### Transitions preceding nuclear export

##### Concerted formation of the E-site

Among the functional centers of the large ribosomal subunit, the E-site is the most variable, with archaeal E-sites containing eL42 and eukaryotic E-sites containing both eL42 and eL29 (*56*, *57*). As the binding site for de-acylated tRNAs during the translation cycle, the E-site is also targeted by several antibiotics, including cycloheximide (*58*, *59*). During the transition from states I to J, the coupled activities of the rixosome and NOG2 (*10*, *60*) trigger their own departure as well as departure of SDA1, NLE1, and CCD86 (yeast Sda1, Rsa4 and Cgr1, respectively) (**Fig. 1, figs. S31, S32**). As a result of the transition from states I to J, the central protuberance undergoes major conformational changes, which form the E-site while also initiating the final stages of PTC maturation.

In the rixosome-bound state I, the central protuberance (CP) is locked in an immature conformation by SDA1, NLE1 and CCD86 (**fig. S31A**). In this context, CCD86 acts as a stabilizer that specifically binds to the rotated immature CP with its N- and C-termini. The conserved N-terminal hook region of CCD86, which was not visualized previously, specifically recognizes RNA and protein elements of the GTPase associated center (GAC) near the assembly factor MRT4. By contrast, the C-terminal region of CCD86, which has evolved into a major protein-interacting region in human cells, now contacts not only the 5*S* RNP via extensive interactions with uL18, but also NLE1 (**fig. S31A,C**). With the removal of the rixosome and many other assembly factors – including NLE1, CCD86 and GNL3 – in state J, the CP adopts a near-mature conformation that is additionally stabilized by ES7c (**fig. S31B**).

On the opposite side of the maturing nuclear pre-60*S* particle, the conformational changes associated with the removal of the rixosome and associated factors lead to the concerted formation of the human E-site (**fig. S32**). In state I, SDA1 splays apart pre-ribosomal RNA elements of domains II and V, thereby preventing the association of eL42 and hence the formation of the E-site (**fig. S32A**). In contrast, following the departure of SDA1 in state J, the compacted RNA helices of domains II and V are stabilized by the newly integrated eL42 and eL29 (fig. S32B). Near the E-site, the C-terminal domain of eL13 (eL13 CTD) stabilizes RNA helices of domains I, II and V, rationalizing how insertions in the eL13 CTD in human patients affect ribosome function without severe effects on ribosome assembly (*61*, *62*).

Due to these major RNA conformational changes that are communicated throughout the entire pre-60*S* particle during the transition from states I to J, we further observe how tail regions of human ribosomal proteins become ordered underneath the E-site, acting as local chaperones that direct final stages of RNA folding (**fig. S32C**). Here the compaction of rRNA elements of domains II and V during the transitions from states I to J occur together with the gradual ordering of parts of universally conserved ribosomal proteins such as uL15, as well as peptides of eukaryote-specific ribosomal proteins including eL13, eL15, eL18, eL21, and eL29. The orchestrated nature with which ribosomal proteins and RNA undergo coordinated folding events underlines one of the key unidirectional drivers of ribosome assembly.

Together, our structural data on the transition from states I to J highlight how large-scale conformational changes are communicated through assembly factors and pre-rRNA so that ribosomal proteins can act as mutual stabilizers of functional centers, rigidifying otherwise flexible states during RNA folding.

**The concerted disassembly of NSA2, NOG2 and GTPB4 prepares pre-60*S* particles for nuclear export**

As described in the states A-to-B transition, the GTPase GTPB4 mediates the GTP hydrolysis-dependent installation of the peptidyl transferase center (PTC) and the GTPase associated Center (GAC) during nucleolar pre-60*S* assembly. Towards the end of nuclear maturation, GDP-bound GTPB4 is among the last assembly factors to dissociate. However, as nucleotide hydrolysis has already occurred in the nucleolus, it has remained unclear how the dissociation of GTPB4 and its associated factors (NSA2 and NOG2) is achieved. While our cryo-EM dataset captures only ITS2-associated intermediates, which represent a subpopulation of maturing pre-60*S* particles, states J to L highlight one pathway for how the remodeling of the GAC results in the dissociation of its bound assembly factors (**fig. S33**).

In state J, the GDP-bound GTPB4 is bound by the N-terminus of NSA2 as well as a peptide of NOG2 (**fig. S33A**). In the subsequent state K, we have identified a new assembly factor, called L10K (C19orf53), which has a mutually exclusive binding site with NSA2 and NOG2. As previously observed during late stages of nuclear maturation (*25*), TMA16 is bound between GTPB4 and MRT4, destabilizing the GAC (**fig. S33B**). In this context, L10K can bind between GTBP4, TMA16, and helix 89 (H89) with its helical C-terminal region. The departure of NSA2 and NOG2 and replacement by L10K results in a more flexible GAC with parts of uL11 becoming disordered. The increased flexibility in this region, together with the reduced stabilization of GTPB4, likely further destabilizes the assembly factors L10K, TMA16 and the structured domains of GTPB4. The dissociation of the structured domains of GTPB4 is required for final stages of PTC maturation that involve the installation of H38 and uL16. As the resulting state L still lacks the ribosomal protein eL40 as well as the export factor NMD3, we believe that this state precedes final nuclear export states as previously observed (*25*). Importantly, state L further indicates how mutations in the P-site loop of uL16 (R98S and Q123R) that have been associated with T-cell acute lymphoblastic leukemia (T-ALL) could impact later stages of assembly (*63*, *64*). As the corresponding residues are still disordered in state L, their mutations likely only affect subsequent stages of PTC maturation and nuclear export once the uL16 P-site loop has been completely integrated.

Together with data presented on the nucleolar role of GTPB4, our structures of late nuclear assembly intermediates of the human pre-60*S* particle provide a model for how TMA16 and the newly identified L10K can bring about the coordinated remodeling of the GAC to favor a subsequent incorporation of uL16.

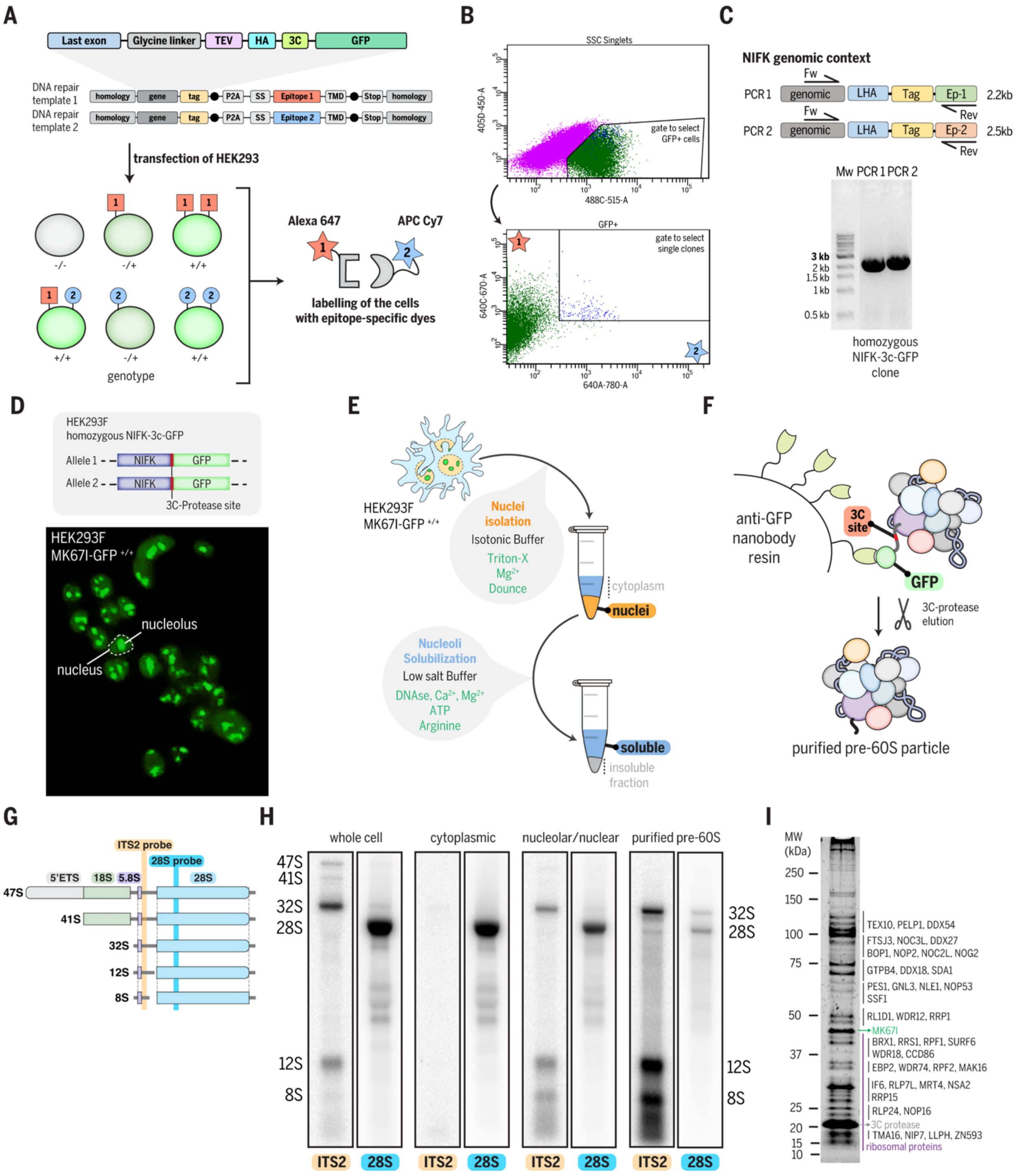

**Fig. S1. Tagging and purification of human pre-60*S* ribosomal subunits.**

(**A**) Strategy for biallelic tagging of the ribosome assembly factor MK67I (gene NIFK) using SNEAK PEEC. Each repair template contains a left homology arm, a portion of the last exon of the gene of interest (gene), a C-terminal tag (tag), a self-cleaving peptide sequence (P2A), a cell-surface display epitope (SS-epitope1/2-TMD) and a right homology arm. Transfection of human cells with the two DNA repair templates and Cas9+sgRNA can result in six different outcomes of cells either containing no edited gene or different monoallelic (-/+) or biallelic (+/+) combinations.

Cells are surface stained using epitope-specific dyes. (**B**) Cells are sorted for single cell clones expressing GFP and both surface displays which represent biallelically edited clones. (**C**) Isolated single clones are genotyped to confirm the genomic integration of both repair templates. In this case, two separate PCR reactions confirm the biallelic tagging of a MK67I-GFP single cell clone. (**D**) Fluorescence microscopy image of HEK293F MK67I-GFP$^{+/+}$ cells showing a nucleolar localization of the protein MK67I. (**E**) Schematic of the fractional lysis purification procedure. (**F**) Schematic of the one-step affinity purification strategy to isolate MK67I-GFP tagged human pre-60*S* particles. (**G**) Schematic of pre-RNAs species detected using ITS2 and 28*S* northern blot probes. (**H**) Northern blot analysis of pre-rRNAs species present in the whole cell, intermediate purification fractions and purified human pre-60*S* particles. Probes hybridizing within ITS2 and 28*S* were used and original Northern blots are depicted in fig. S2. (**I**) SYPRO Ruby stained SDS-PAGE analysis of the purified pre-60*S* sample.

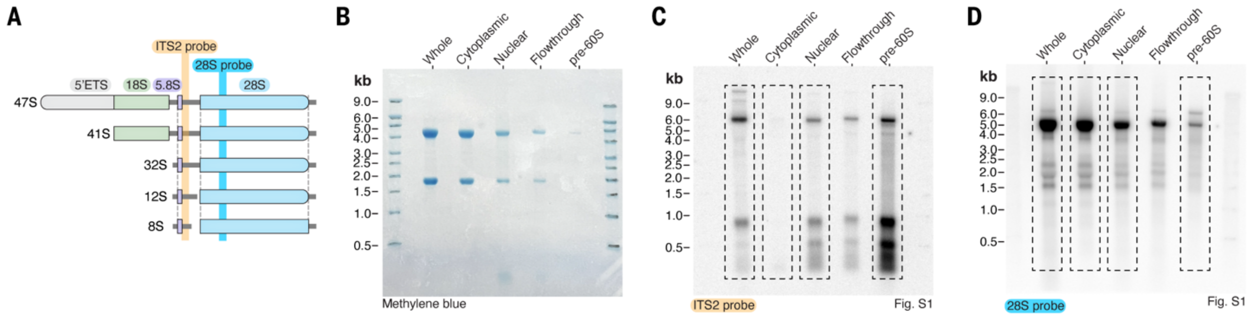


**Fig. S2. Northern blot analysis of purified human pre-60*S* particles.**

(**A**) Schematic of pre-RNAs species detected using ITS2 and 28*S* northern blot probes. (**B**) Methylene blue staining showing 18*S* and 28*S* RNA bands detected from different fractions of the pre-60*S* purification. (**C**) Northern blot analysis using the ITS2 probe showing the different pre-rRNAs species in each fraction of the pre-60*S* purification. (**D)** Northern blot analysis using the 28*S* probe showing bands corresponding to the mature 28*S* rRNA and its precursor 32*S* pre-rRNA. The dashed squares show the cropped regions used in fig. S1.

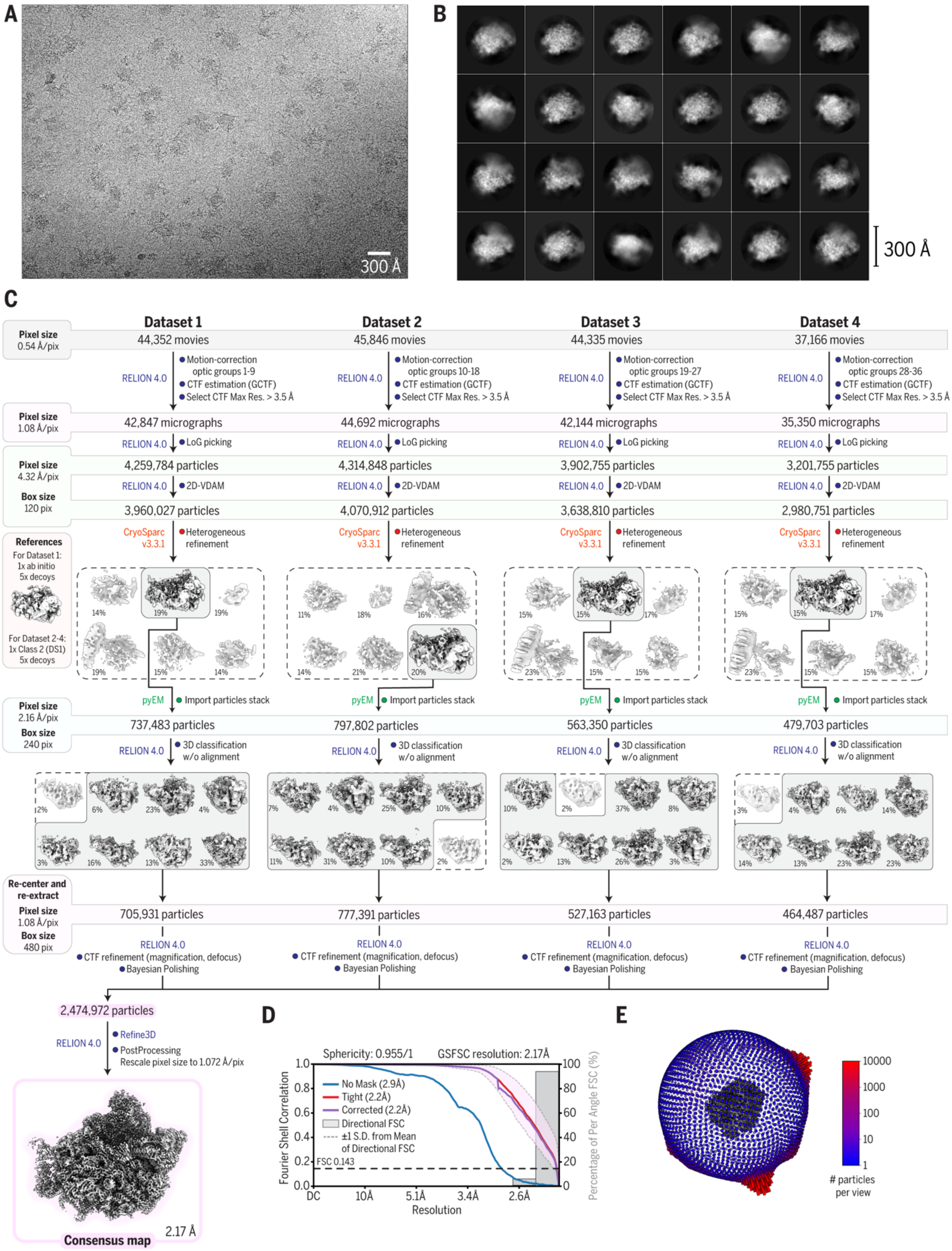
A
300 Å
B
300 Å
C
Dataset 1
Dataset 2
Dataset 3
Dataset 4
Pixel size 0.54 Å/pix
44,352 movies
45,846 movies
44,335 movies
37,166 movies
RELION 4.0
Motion-correction optic groups 1-9
CTF estimation (GCTF)
Select CTF Max Res. > 3.5 Å
Pixel size 1.08 Å/pix
42,847 micrographs
44,692 micrographs
42,144 micrographs
35,350 micrographs
LoG picking
Pixel size 4.32 Å/pix
Box size 120 pix
4,259,784 particles
4,314,848 particles
3,902,755 particles
3,201,755 particles
2D-VDAM
3,960,027 particles
4,070,912 particles
3,638,810 particles
2,980,751 particles
CryoSparc v3.3.1
Heterogeneous refinement
References
For Dataset 1: 1x ab initio 5x decoys
For Dataset 2-4: 1x Class 2 (DS1) 5x decoys
pyEM
Import particles stack
Pixel size 2.16 Å/pix
Box size 240 pix
737,483 particles
797,802 particles
563,350 particles
479,703 particles
3D classification w/o alignment
Re-center and re-extract
Pixel size 1.08 Å/pix
Box size 480 pix
705,931 particles
777,391 particles
527,163 particles
464,487 particles
CTF refinement (magnification, defocus)
Bayesian Polishing
2,474,972 particles
Refine3D
PostProcessing
Rescale pixel size to 1.072 Å/pix
Consensus map
2.17 Å
D
Sphericity: 0.955/1
GSFSC resolution: 2.17Å
No Mask (2.9Å)
Tight (2.2Å)
Corrected (2.2Å)
Directional FSC
±1 S.D. from Mean of Directional FSC
FSC 0.143
Fourier Shell Correlation
Percentage of Per Angle FSC (%)
Resolution
DC
10Å
5.1Å
3.4Å
2.6Å
E
### particles per view

**Fig. S3. Initial cryo-EM processing.**

(**A**) Representative cryo-EM micrograph. (**B**) Unsupervised 2D class averages. (**C**) Cryo-EM data processing workflow for the determination of a consensus human pre-60*S* ribosomal subunit. Software used and type of jobs are indicated. (**D**) FSC curves (no mask, tight mask, solvent corrected, 3D). The resolution is indicated for each curve and was determined at FSC-0.143. (**E**) Euler angle distribution for the consensus reconstruction.

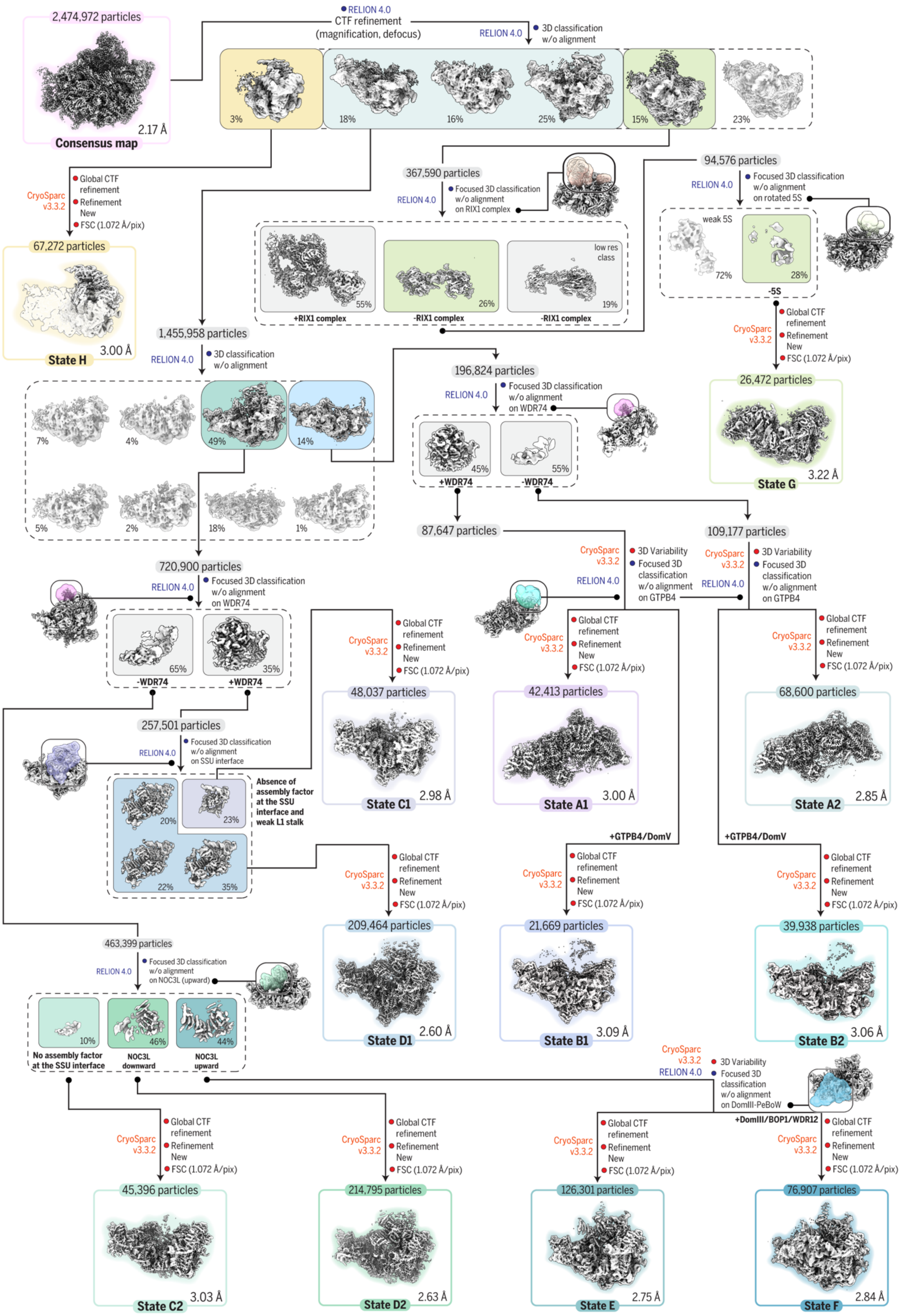

2,474,972 particles
2.17 Å
Consensus map
RELION 4.0
CTF refinement
(magnification, defocus)
RELION 4.0
3D classification
w/o alignment
3%
18%
16%
25%
15%
23%
CryoSparc v3.3.2
Global CTF refinement
Refinement New
FSC (1.072 Å/pix)
67,272 particles
3.00 Å
State H
367,590 particles
RELION 4.0
Focused 3D classification
w/o alignment
on RIX1 complex
55%
26%
19%
low res class
+RIX1 complex
-RIX1 complex
-RIX1 complex
94,576 particles
RELION 4.0
Focused 3D classification
w/o alignment
on rotated 5S
weak 5S
72%
28%
-5S
CryoSparc v3.3.2
Global CTF refinement
Refinement New
FSC (1.072 Å/pix)
26,472 particles
3.22 Å
State G
1,455,958 particles
RELION 4.0
3D classification
w/o alignment
7%
4%
49%
14%
5%
2%
18%
1%
196,824 particles
RELION 4.0
Focused 3D classification
w/o alignment
on WDR74
45%
55%
+WDR74
-WDR74
87,647 particles
109,177 particles
CryoSparc v3.3.2
RELION 4.0
3D Variability
Focused 3D classification
w/o alignment
on GTPB4
CryoSparc v3.3.2
RELION 4.0
3D Variability
Focused 3D classification
w/o alignment
on GTPB4
720,900 particles
RELION 4.0
Focused 3D classification
w/o alignment
on WDR74
65%
35%
-WDR74
+WDR74
CryoSparc v3.3.2
Global CTF refinement
Refinement New
FSC (1.072 Å/pix)
48,037 particles
2.98 Å
State C1
CryoSparc v3.3.2
Global CTF refinement
Refinement New
FSC (1.072 Å/pix)
42,413 particles
3.00 Å
State A1
CryoSparc v3.3.2
Global CTF refinement
Refinement New
FSC (1.072 Å/pix)
68,600 particles
2.85 Å
State A2
257,501 particles
RELION 4.0
Focused 3D classification
w/o alignment
on SSU interface
20%
23%
22%
35%
Absence of assembly factor at the SSU interface and weak L1 stalk
+GTPB4/DomV
+GTPB4/DomV
CryoSparc v3.3.2
Global CTF refinement
Refinement New
FSC (1.072 Å/pix)
209,464 particles
2.60 Å
State D1
CryoSparc v3.3.2
Global CTF refinement
Refinement New
FSC (1.072 Å/pix)
21,669 particles
3.09 Å
State B1
CryoSparc v3.3.2
Global CTF refinement
Refinement New
FSC (1.072 Å/pix)
39,938 particles
3.06 Å
State B2
463,399 particles
RELION 4.0
Focused 3D classification
w/o alignment
on NOC3L (upward)
10%
46%
44%
No assembly factor at the SSU interface
NOC3L downward
NOC3L upward
CryoSparc v3.3.2
RELION 4.0
3D Variability
Focused 3D classification
w/o alignment
on DomIII-PeBoW
+DomIII/BOP1/WDR12
CryoSparc v3.3.2
Global CTF refinement
Refinement New
FSC (1.072 Å/pix)
45,396 particles
3.03 Å
State C2
CryoSparc v3.3.2
Global CTF refinement
Refinement New
FSC (1.072 Å/pix)
214,795 particles
2.63 Å
State D2
CryoSparc v3.3.2
Global CTF refinement
Refinement New
FSC (1.072 Å/pix)
126,301 particles
2.75 Å
State E
CryoSparc v3.3.2
Global CTF refinement
Refinement New
FSC (1.072 Å/pix)
76,907 particles
2.84 Å
State F

**Fig. S4. Cryo-EM processing and reconstruction of human pre-60*S* nucleolar assembly intermediates.**

Cryo-EM data processing workflow for the identification and reconstruction of 12 pre-60*S* nucleolar assembly intermediates (states A1 to H). Software used and type of jobs are indicated. Map resolutions were determined at FSC-0.143.

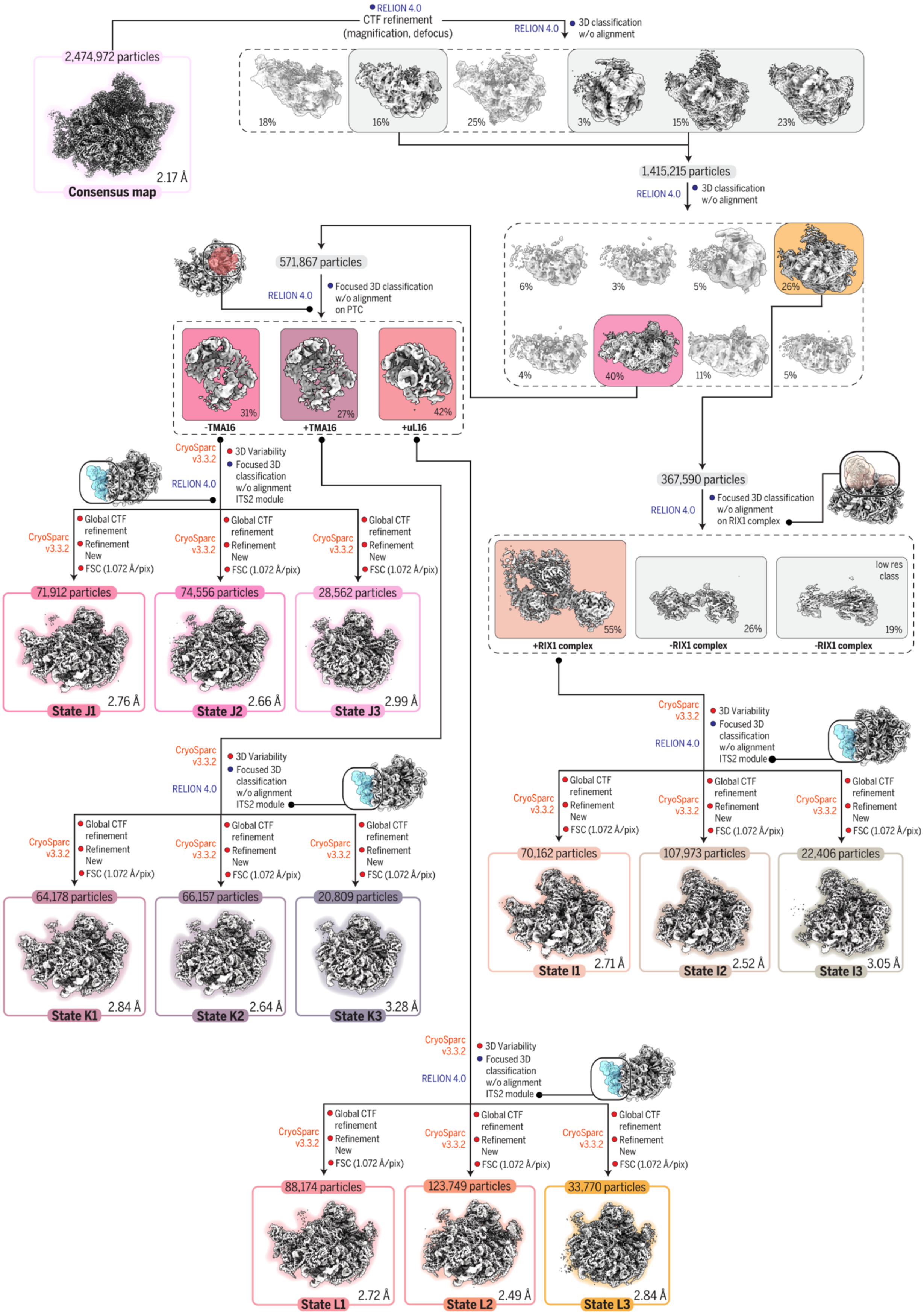

RELION 4.0
CTF refinement
(magnification, defocus)
RELION 4.0
3D classification
w/o alignment
2,474,972 particles
2.17 Å
Consensus map
18%
16%
25%
3%
15%
23%
1,415,215 particles
RELION 4.0
3D classification
w/o alignment
6%
3%
5%
26%
4%
40%
11%
5%
571,867 particles
RELION 4.0
Focused 3D classification
w/o alignment
on PTC
31%
27%
42%
-TMA16
+TMA16
+uL16
367,590 particles
RELION 4.0
Focused 3D classification
w/o alignment
on RIX1 complex
55%
26%
low res
class
19%
+RIX1 complex
-RIX1 complex
-RIX1 complex
CryoSparc
v3.3.2
3D Variability
RELION 4.0
Focused 3D
classification
w/o alignment
ITS2 module
Global CTF
refinement
Refinement
New
FSC (1.072 Å/pix)
71,912 particles
2.76 Å
State J1
74,556 particles
2.66 Å
State J2
28,562 particles
2.99 Å
State J3
64,178 particles
2.84 Å
State K1
66,157 particles
2.64 Å
State K2
20,809 particles
3.28 Å
State K3
70,162 particles
2.71 Å
State I1
107,973 particles
2.52 Å
State I2
22,406 particles
3.05 Å
State I3
88,174 particles
2.72 Å
State L1
123,749 particles
2.49 Å
State L2
33,770 particles
2.84 Å
State L3

**Fig. S5. Cryo-EM processing and reconstruction of human pre-60*S* nuclear assembly intermediates.**

Cryo-EM data processing workflow for the identification and reconstruction of 12 pre-60*S* nuclear assembly intermediates (states I1 to L3). Software used and type of jobs are indicated. Map resolutions were determined at FSC-0.143.

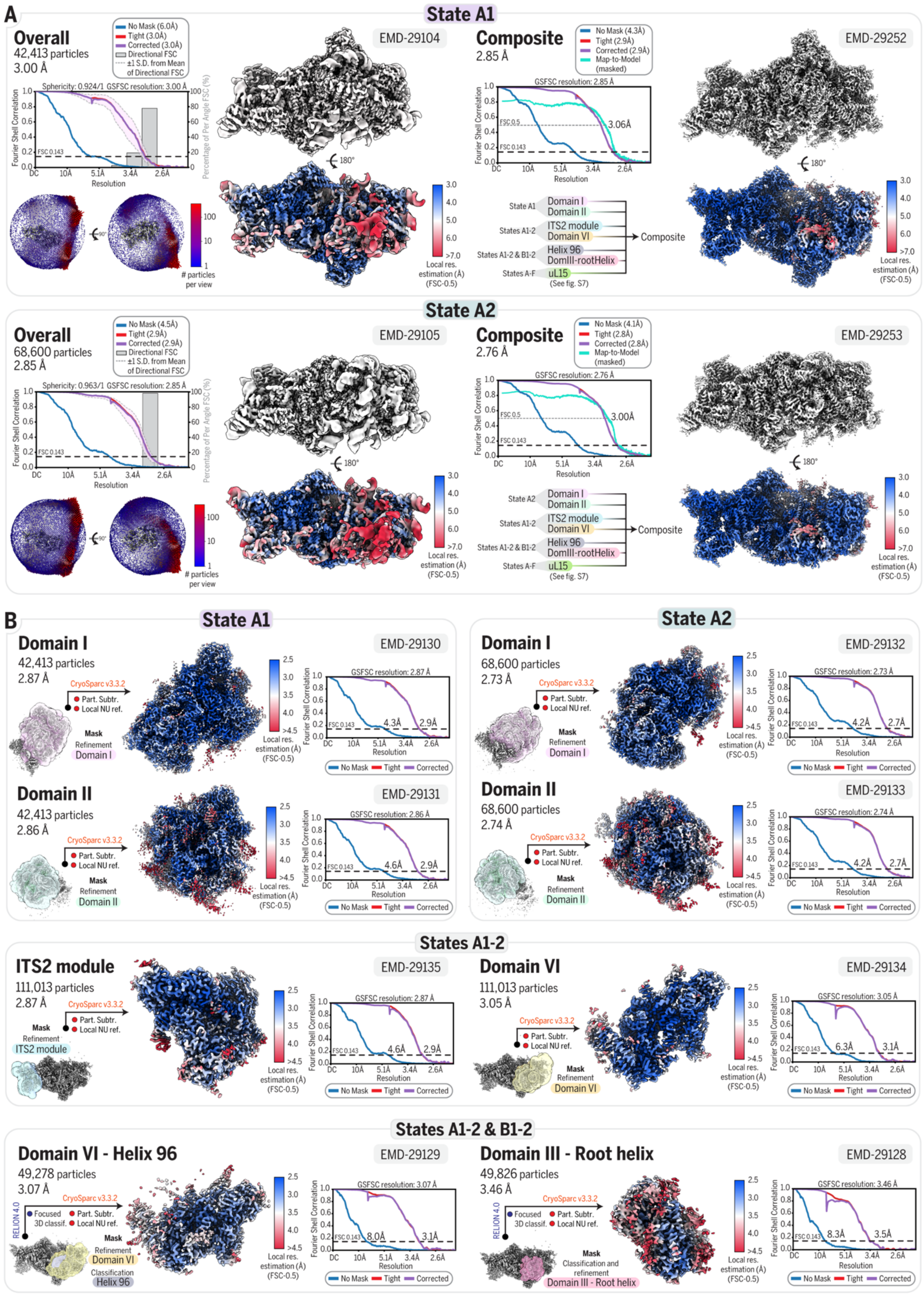

A
State A1
Overall
42,413 particles
3.00 Å
EMD-29104
Composite
2.85 Å
EMD-29252
State A2
Overall
68,600 particles
2.85 Å
EMD-29105
Composite
2.76 Å
EMD-29253
B
State A1
Domain I
42,413 particles
2.87 Å
EMD-29130
Domain II
42,413 particles
2.86 Å
EMD-29131
State A2
Domain I
68,600 particles
2.73 Å
EMD-29132
Domain II
68,600 particles
2.74 Å
EMD-29133
States A1-2
ITS2 module
111,013 particles
2.87 Å
EMD-29135
Domain VI
111,013 particles
3.05 Å
EMD-29134
States A1-2 & B1-2
Domain VI - Helix 96
49,278 particles
3.07 Å
EMD-29129
Domain III - Root helix
49,826 particles
3.46 Å
EMD-29128

**Fig. S6. Composite cryo-EM reconstruction of states A1 and A2.**

(**A**) Overall and composite cryo-EM maps of state A1 and state A2. For the overall reconstructions, FSC curves (no mask, tight mask, solvent corrected and 3D) and Euler angle distribution are displayed on the left. For the composite maps, the FSC curves (no mask, tight mask, solvent corrected, map-to-model) and a list of the selected focused maps used to generate the composite map are displayed on the left. (**B**) Focused refinements for relevant modules of state A1 and state A2. The mask used for each focus refinement is shown with respect to the reference map. The resulting focused map is colored by local resolution estimation. FSC curves (no mask, tight mask, solvent corrected) are displayed on the right. Software used and type of jobs are indicated. In some cases, when relevant, particles stack from different states are merged to improve the reconstruction of a specific module common to the different states. (**A-B**) The resolution indicated for all FSC curve was determined at FSC-0.143, except for the map-to-model FSC which was determined at FSC-0.5. The Electron Microscopy Data Bank (EMDB) accession number is indicated for each map.

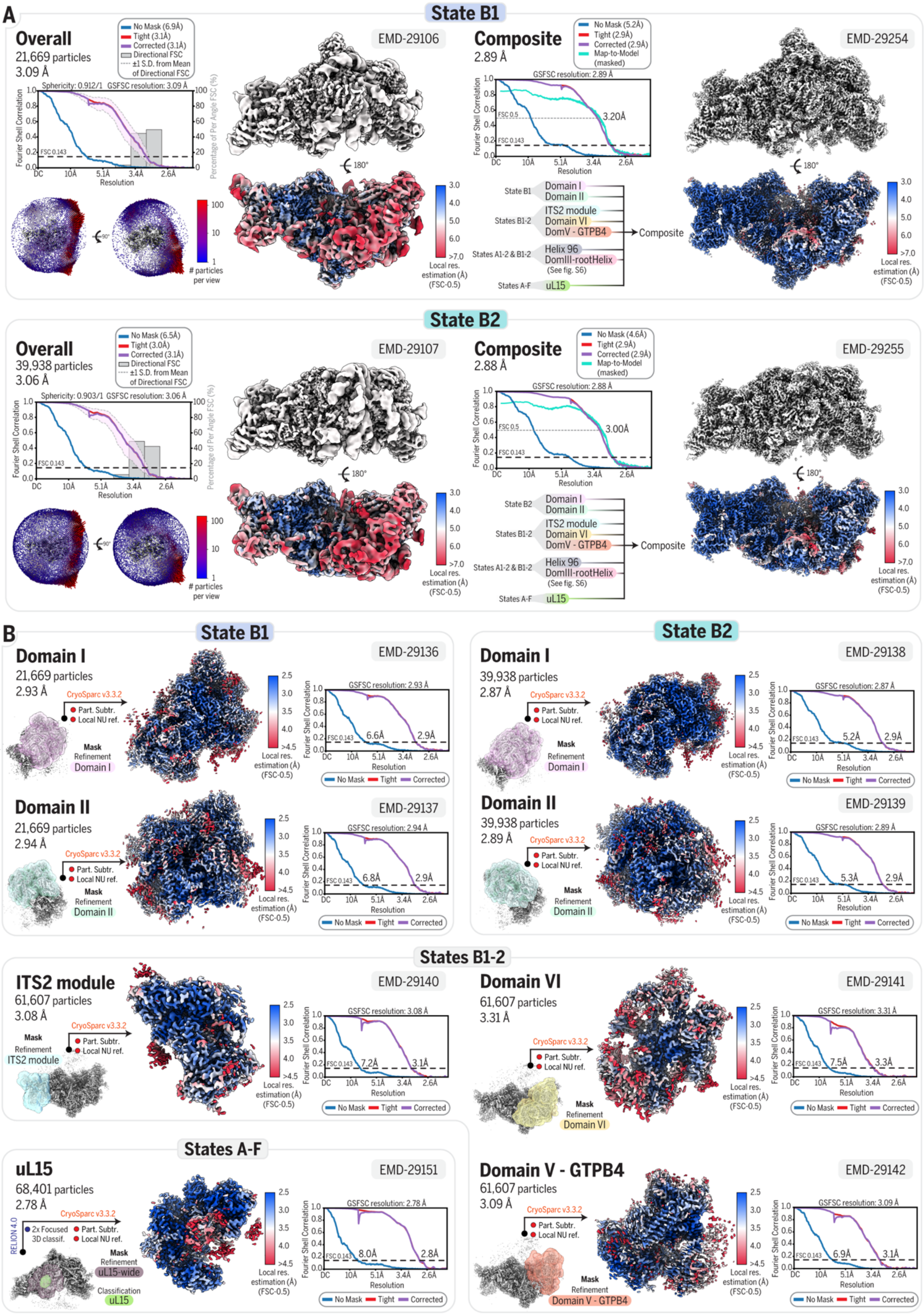
A
State B1
Overall
21,669 particles
3.09 Å
No Mask (6.9Å)
Tight (3.1Å)
Corrected (3.1Å)
Directional FSC
±1 S.D. from Mean of Directional FSC
Sphericity: 0.912/1 GSFSC resolution: 3.09 Å
Fourier Shell Correlation
Percentage of Per Angle FSC (%)
FSC 0.143
Resolution
### particles per view
EMD-29106
180°
Local res. estimation (Å) (FSC-0.5)
Composite
2.89 Å
No Mask (5.2Å)
Tight (2.9Å)
Corrected (2.9Å)
Map-to-Model (masked)
GSFSC resolution: 2.89 Å
FSC 0.5
3.20Å
State B1
Domain I
Domain II
States B1-2
ITS2 module
Domain VI
DomV - GTPB4
Composite
States A1-2 & B1-2
Helix 96
DomIII-rootHelix
(See fig. S6)
States A-F
uL15
EMD-29254
State B2
Overall
39,938 particles
3.06 Å
No Mask (6.5Å)
Tight (3.0Å)
Corrected (3.1Å)
Directional FSC
±1 S.D. from Mean of Directional FSC
Sphericity: 0.903/1 GSFSC resolution: 3.06 Å
EMD-29107
Composite
2.88 Å
No Mask (4.6Å)
Tight (2.9Å)
Corrected (2.9Å)
Map-to-Model (masked)
GSFSC resolution: 2.88 Å
3.00Å
State B2
EMD-29255
B
State B1
Domain I
21,669 particles
2.93 Å
CryoSparc v3.3.2
Part. Subtr.
Local NU ref.
Mask Refinement Domain I
EMD-29136
GSFSC resolution: 2.93 Å
6.6Å
2.9Å
No Mask Tight Corrected
Domain II
21,669 particles
2.94 Å
Mask Refinement Domain II
EMD-29137
GSFSC resolution: 2.94 Å
6.8Å
2.9Å
State B2
Domain I
39,938 particles
2.87 Å
EMD-29138
GSFSC resolution: 2.87 Å
5.2Å
2.9Å
Domain II
39,938 particles
2.89 Å
EMD-29139
GSFSC resolution: 2.89 Å
5.3Å
2.9Å
States B1-2
ITS2 module
61,607 particles
3.08 Å
Mask Refinement ITS2 module
EMD-29140
GSFSC resolution: 3.08 Å
7.2Å
3.1Å
Domain VI
61,607 particles
3.31 Å
Mask Refinement Domain VI
EMD-29141
GSFSC resolution: 3.31 Å
7.5Å
3.3Å
States A-F
uL15
68,401 particles
2.78 Å
RELION 4.0
2x Focused 3D classif.
Mask Refinement uL15-wide
Classification uL15
EMD-29151
GSFSC resolution: 2.78 Å
8.0Å
2.8Å
Domain V - GTPB4
61,607 particles
3.09 Å
Mask Refinement Domain V - GTPB4
EMD-29142
GSFSC resolution: 3.09 Å
6.9Å
3.1Å

**Fig. S7. Composite cryo-EM reconstruction of states B1 and B2.**

(**A**) Overall and composite cryo-EM maps of state B1 and state B2. For the overall reconstructions, FSC curves (no mask, tight mask, solvent corrected and 3D) and Euler angle distribution are displayed on the left. For the composite maps, the FSC curves (no mask, tight mask, solvent corrected, map-to-model) and a list of the selected focused maps used to generate the composite map are displayed on the left. (**B**) Focused refinements for relevant modules of state B1 and state B2. The mask used for each focus refinement is shown with respect to the reference map. The resulting focused map is colored by local resolution estimation. FSC curves (no mask, tight mask, solvent corrected) are displayed on the right. Software used and type of jobs are indicated. In some cases, when relevant, particles stack from different states are merged to improve the reconstruction of a specific module common to the different states. (**A-B**) The resolution indicated for all FSC curve was determined at FSC-0.143, except for the map-to-model FSC which was determined at FSC-0.5. The Electron Microscopy Data Bank (EMDB) accession number is indicated for each map.

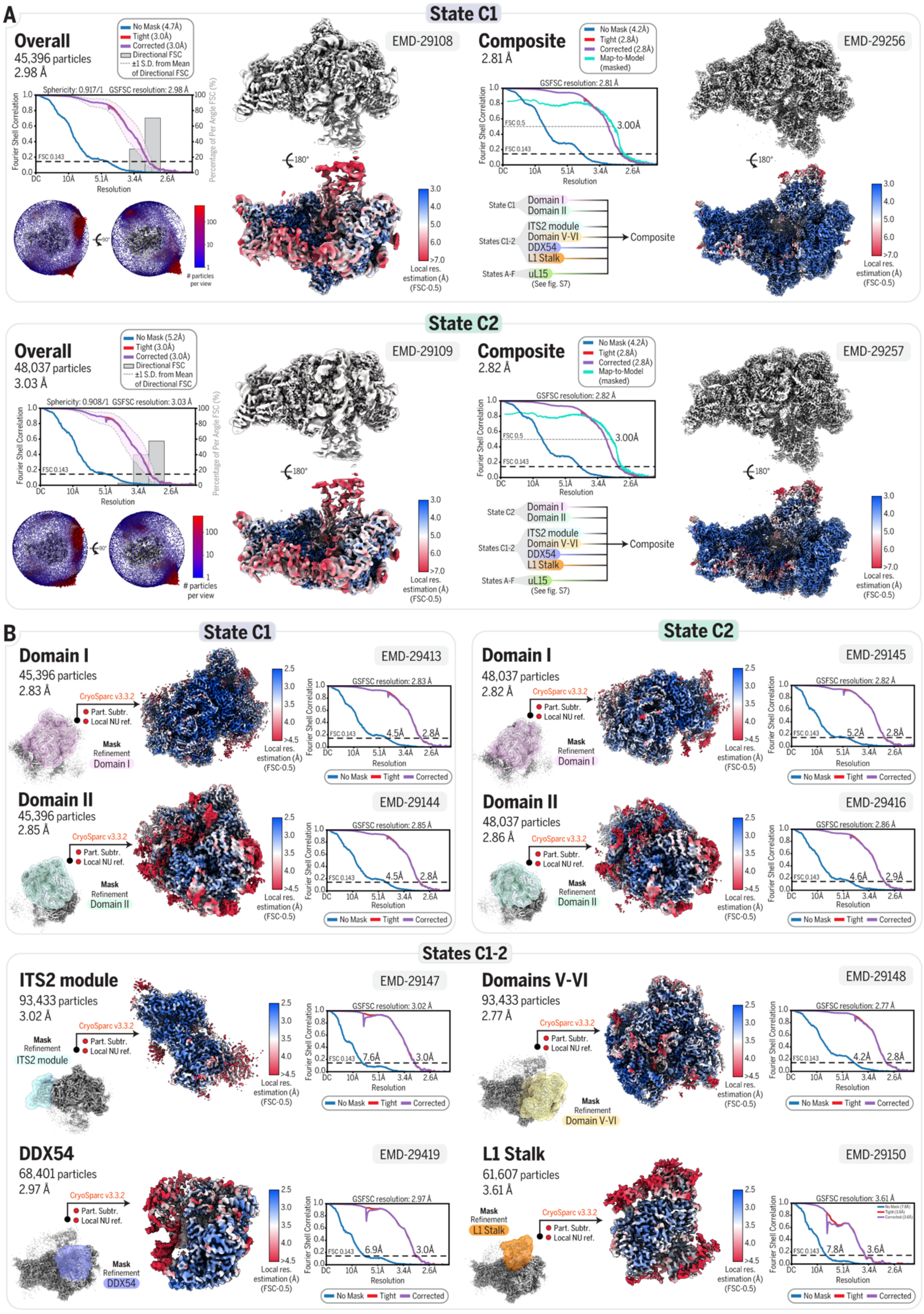
A
State C1
Overall
45,396 particles
2.98 Å
No Mask (4.7Å)
Tight (3.0Å)
Corrected (3.0Å)
Directional FSC
±1 S.D. from Mean of Directional FSC
Sphericity: 0.917/1
GSFSC resolution: 2.98 Å
Fourier Shell Correlation
Percentage of Per Angle FSC (%)
FSC 0.143
Resolution
### particles per view
EMD-29108
180°
Local res. estimation (Å) (FSC-0.5)
Composite
2.81 Å
No Mask (4.2Å)
Tight (2.8Å)
Corrected (2.8Å)
Map-to-Model (masked)
GSFSC resolution: 2.81 Å
FSC 0.5
3.00Å
EMD-29256
State C1
Domain I
Domain II
States C1-2
ITS2 module
Domain V-VI
DDX54
L1 Stalk
States A-F
uL15
(See fig. S7)
Composite
State C2
Overall
48,037 particles
3.03 Å
No Mask (5.2Å)
Tight (3.0Å)
Corrected (3.0Å)
Sphericity: 0.908/1
GSFSC resolution: 3.03 Å
EMD-29109
Composite
2.82 Å
GSFSC resolution: 2.82 Å
EMD-29257
B
State C1
Domain I
45,396 particles
2.83 Å
CryoSparc v3.3.2
Part. Subtr.
Local NU ref.
Mask Refinement
EMD-29413
GSFSC resolution: 2.83 Å
4.5Å
2.8Å
No Mask
Tight
Corrected
Domain II
45,396 particles
2.85 Å
EMD-29144
GSFSC resolution: 2.85 Å
State C2
Domain I
48,037 particles
2.82 Å
EMD-29145
GSFSC resolution: 2.82 Å
5.2Å
Domain II
48,037 particles
2.86 Å
EMD-29416
GSFSC resolution: 2.86 Å
4.6Å
2.9Å
States C1-2
ITS2 module
93,433 particles
3.02 Å
EMD-29147
GSFSC resolution: 3.02 Å
7.6Å
3.0Å
Domains V-VI
93,433 particles
2.77 Å
EMD-29148
GSFSC resolution: 2.77 Å
4.2Å
Domain V-VI
DDX54
68,401 particles
2.97 Å
EMD-29419
GSFSC resolution: 2.97 Å
6.9Å
L1 Stalk
61,607 particles
3.61 Å
EMD-29150
GSFSC resolution: 3.61 Å
7.8Å
3.6Å

**Fig. S8. Composite cryo-EM reconstruction of states C1 and C2.**

(**A**) Overall and composite cryo-EM maps of state C1 and state C2. For the overall reconstructions, FSC curves (no mask, tight mask, solvent corrected and 3D) and Euler angle distribution are displayed on the left. For the composite maps, the FSC curves (no mask, tight mask, solvent corrected, map-to-model) and a list of the selected focused maps used to generate the composite map are displayed on the left. (**B**) Focused refinements for relevant modules of state C1 and state C2. The mask used for each focus refinement is shown with respect to the reference map. The resulting focused map is colored by local resolution estimation. FSC curves (no mask, tight mask, solvent corrected) are displayed on the right. Software used and type of jobs are indicated. In some cases, when relevant, particles stack from different states are merged to improve the reconstruction of a specific module common to the different states. (**A-B**) The resolution indicated for all FSC curve was determined at FSC-0.143, except for the map-to-model FSC which was determined at FSC-0.5. The Electron Microscopy Data Bank (EMDB) accession number is indicated for each map.

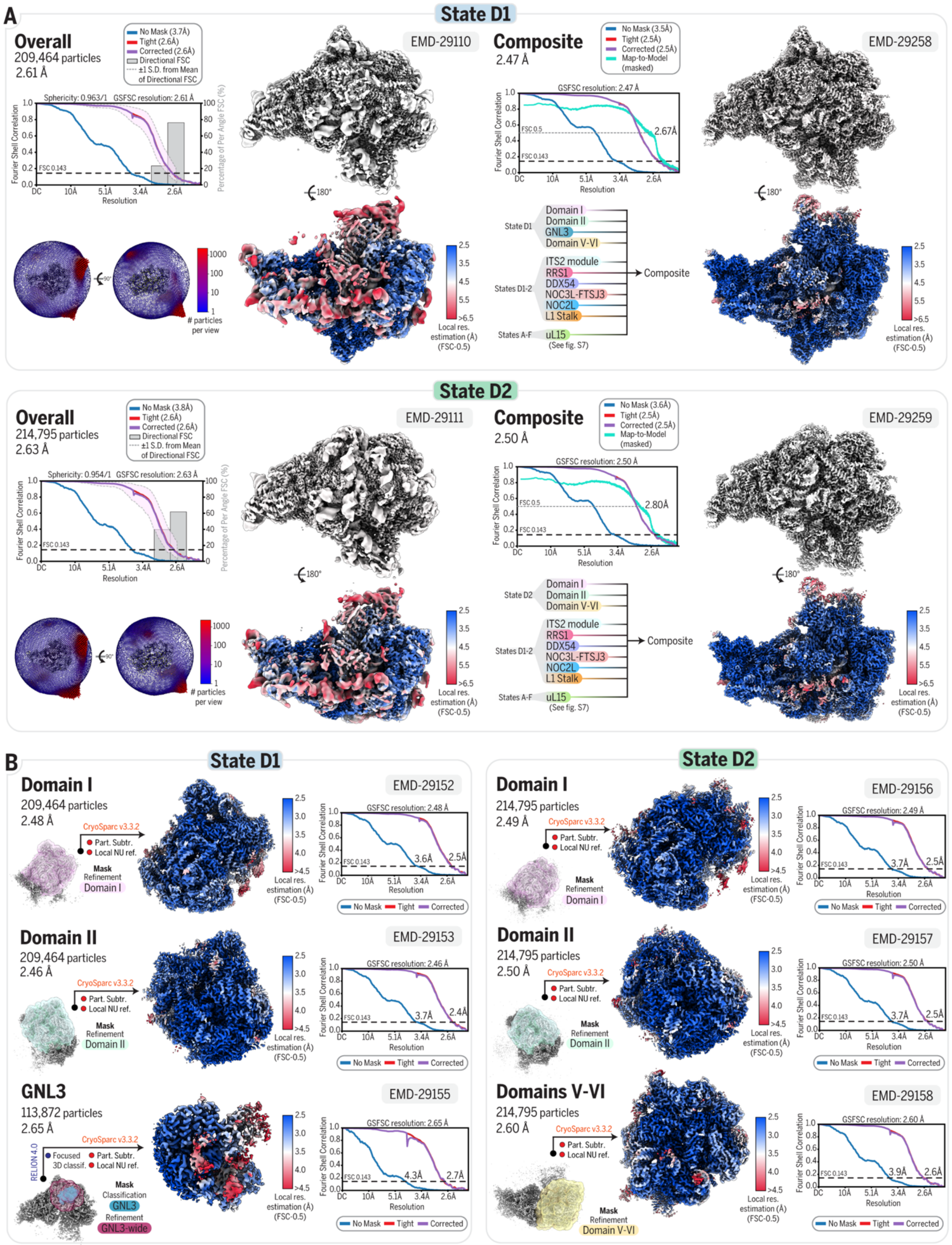

A
State D1
Overall
209,464 particles
2.61 Å
No Mask (3.7Å)
Tight (2.6Å)
Corrected (2.6Å)
Directional FSC
±1 S.D. from Mean of Directional FSC
Sphericity: 0.963/1 GSFSC resolution: 2.61 Å
Fourier Shell Correlation
Percentage of Per Angle FSC (%)
Resolution
EMD-29110
Composite
2.47 Å
No Mask (3.5Å)
Tight (2.5Å)
Corrected (2.5Å)
Map-to-Model (masked)
GSFSC resolution: 2.47 Å
2.67Å
EMD-29258
### particles per view
Local res. estimation (Å) (FSC-0.5)
State D1
Domain I
Domain II
GNL3
Domain V-VI
States D1-2
ITS2 module
RRS1
DDX54
NOC3L-FTSJ3
NOC2L
L1 Stalk
Composite
States A-F
uL15
(See fig. S7)
State D2
Overall
214,795 particles
2.63 Å
No Mask (3.8Å)
Tight (2.6Å)
Corrected (2.6Å)
Directional FSC
±1 S.D. from Mean of Directional FSC
Sphericity: 0.954/1 GSFSC resolution: 2.63 Å
EMD-29111
Composite
2.50 Å
No Mask (3.6Å)
Tight (2.5Å)
Corrected (2.5Å)
Map-to-Model (masked)
GSFSC resolution: 2.50 Å
2.80Å
EMD-29259
State D2
Domain I
Domain II
Domain V-VI
States D1-2
ITS2 module
RRS1
DDX54
NOC3L-FTSJ3
NOC2L
L1 Stalk
Composite
States A-F
uL15
(See fig. S7)
B
State D1
Domain I
209,464 particles
2.48 Å
CryoSparc v3.3.2
Part. Subtr.
Local NU ref.
Mask
Refinement
Domain I
EMD-29152
GSFSC resolution: 2.48 Å
3.6Å
2.5Å
No Mask
Tight
Corrected
Domain II
209,464 particles
2.46 Å
CryoSparc v3.3.2
Part. Subtr.
Local NU ref.
Mask
Refinement
Domain II
EMD-29153
GSFSC resolution: 2.46 Å
3.7Å
2.4Å
GNL3
113,872 particles
2.65 Å
RELION 4.0
CryoSparc v3.3.2
Focused 3D classif.
Part. Subtr.
Local NU ref.
Mask
Classification
GNL3
Refinement
GNL3-wide
EMD-29155
GSFSC resolution: 2.65 Å
4.3Å
2.7Å
State D2
Domain I
214,795 particles
2.49 Å
CryoSparc v3.3.2
Part. Subtr.
Local NU ref.
Mask
Refinement
Domain I
EMD-29156
GSFSC resolution: 2.49 Å
3.7Å
2.5Å
Domain II
214,795 particles
2.50 Å
CryoSparc v3.3.2
Part. Subtr.
Local NU ref.
Mask
Refinement
Domain II
EMD-29157
GSFSC resolution: 2.50 Å
3.7Å
2.5Å
Domains V-VI
214,795 particles
2.60 Å
CryoSparc v3.3.2
Part. Subtr.
Local NU ref.
Mask
Refinement
Domain V-VI
EMD-29158
GSFSC resolution: 2.60 Å
3.9Å
2.6Å

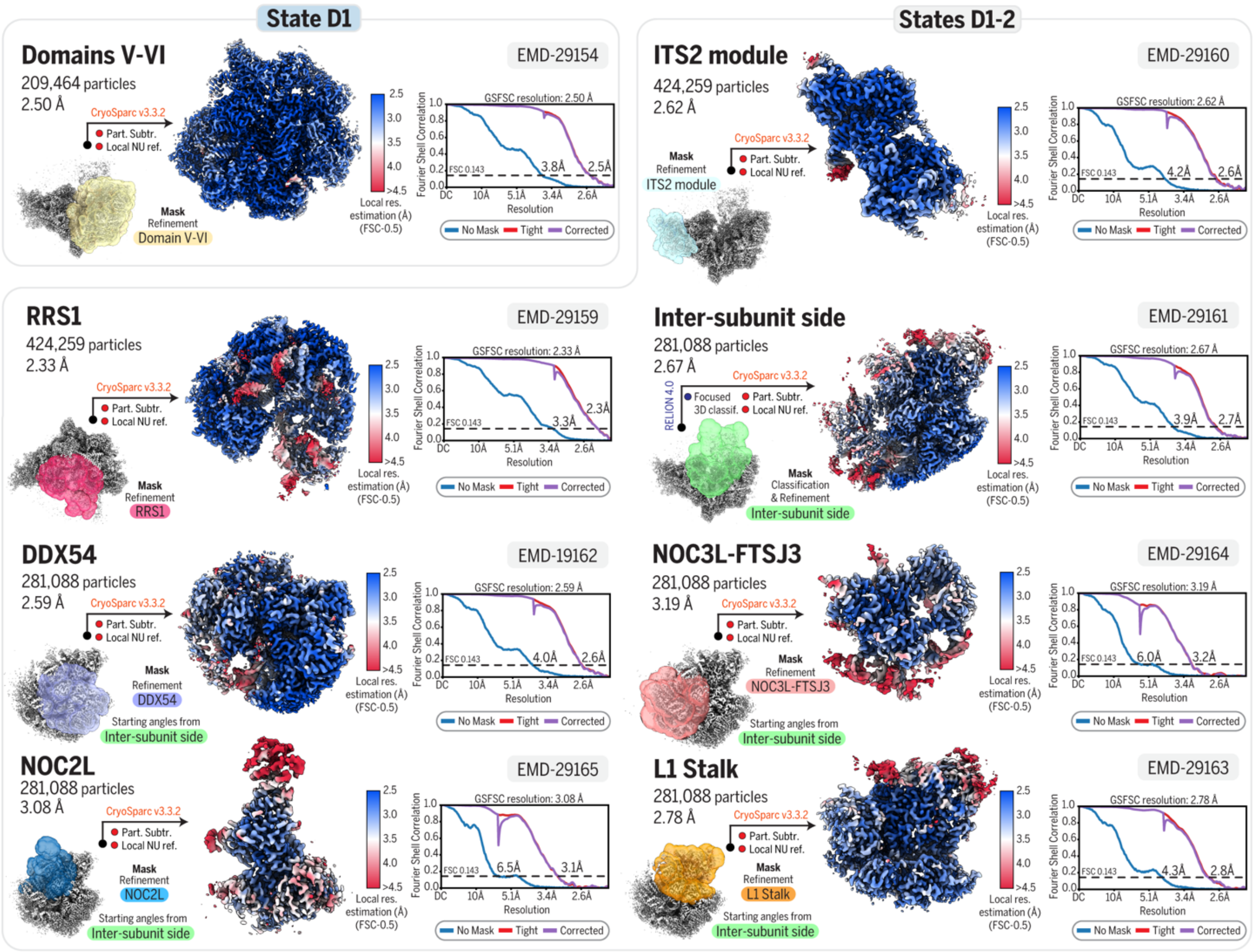


**Fig. S9. Composite cryo-EM reconstruction of states D1 and D2.**

(**A**) Overall and composite cryo-EM maps of state D1 and state D2. For the overall reconstructions, FSC curves (no mask, tight mask, solvent corrected and 3D) and Euler angle distribution are displayed on the left. For the composite maps, the FSC curves (no mask, tight mask, solvent corrected, map-to-model) and a list of the selected focused maps used to generate the composite map are displayed on the left. (**B**) Focused refinements for relevant modules of state D1 and state D2. The mask used for each focus refinement is shown with respect to the reference map. The resulting focused map is colored by local resolution estimation. FSC curves (no mask, tight mask, solvent corrected) are displayed on the right. Software used and type of jobs are indicated. In some cases, when relevant, particles stack from different states are merged to improve the reconstruction of a specific module common to the different states. (**A-B**) The resolution indicated for all FSC curve was determined at FSC-0.143, except for the map-to-model FSC which was determined at FSC-0.5. The Electron Microscopy Data Bank (EMDB) accession number is indicated for each map.

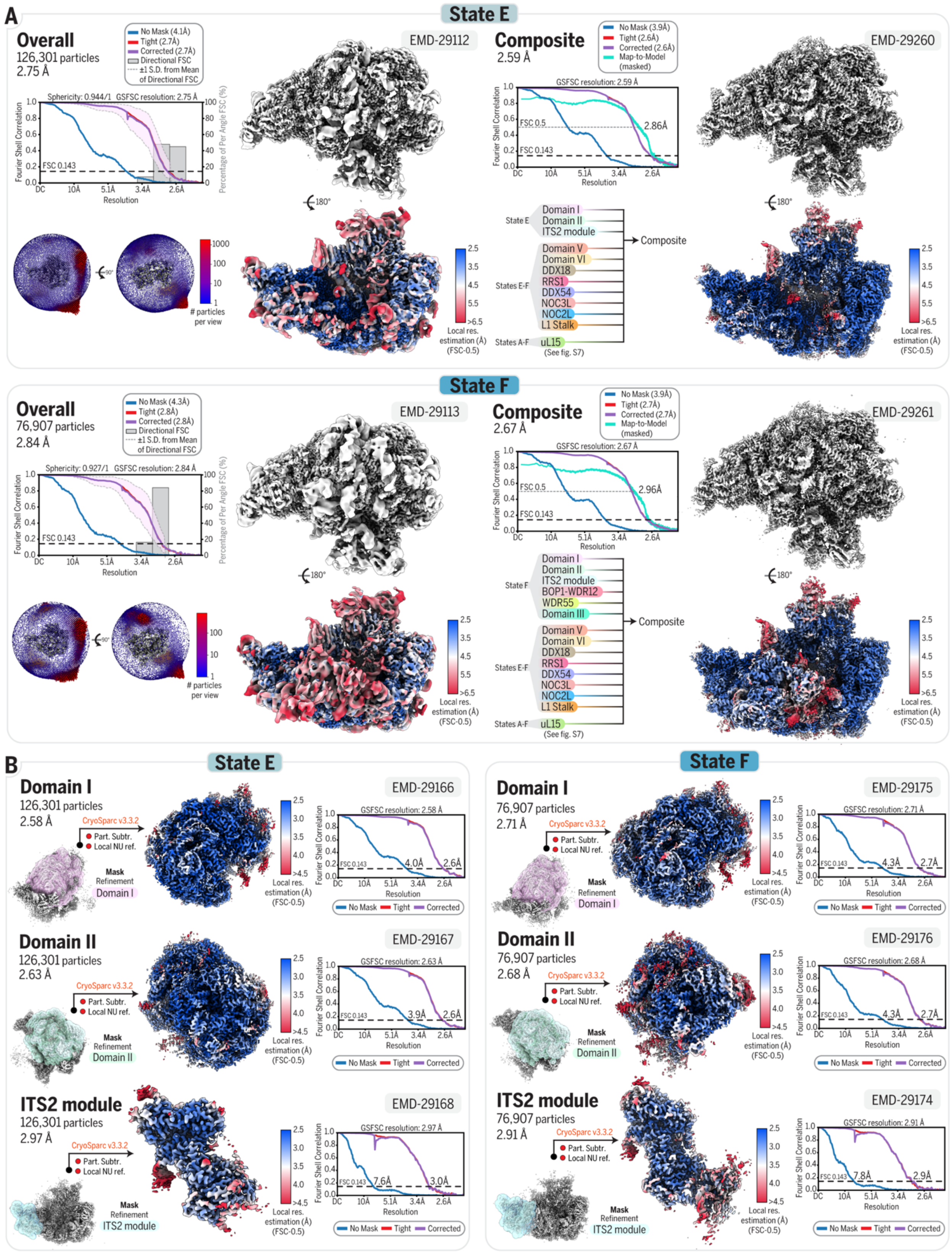

A
State E
Overall
126,301 particles
2.75 Å
EMD-29112
Composite
2.59 Å
EMD-29260
State F
Overall
76,907 particles
2.84 Å
EMD-29113
Composite
2.67 Å
EMD-29261
B
State E
Domain I
126,301 particles
2.58 Å
EMD-29166
Domain II
126,301 particles
2.63 Å
EMD-29167
ITS2 module
126,301 particles
2.97 Å
EMD-29168
State F
Domain I
76,907 particles
2.71 Å
EMD-29175
Domain II
76,907 particles
2.68 Å
EMD-29176
ITS2 module
76,907 particles
2.91 Å
EMD-29174

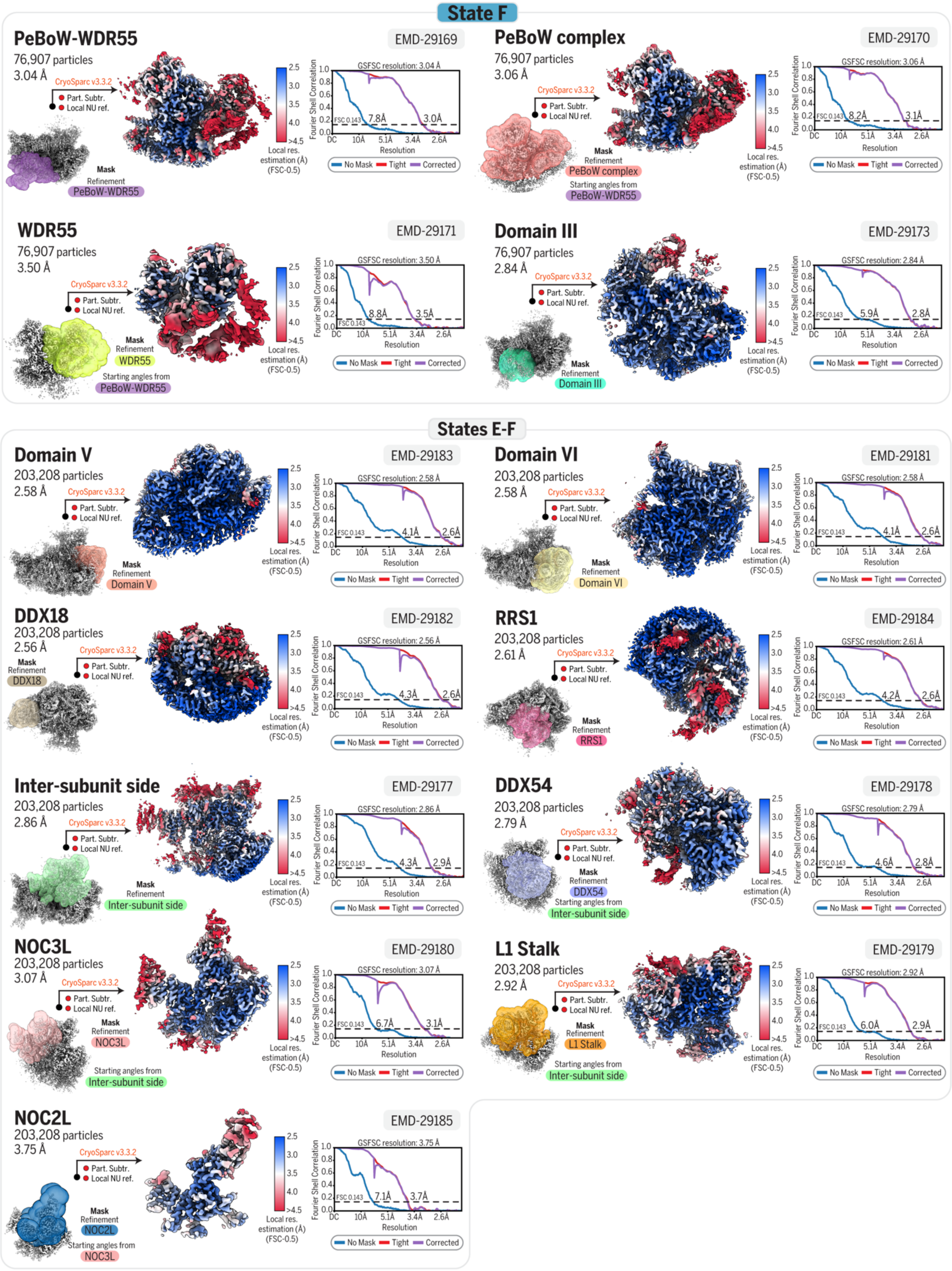
State F
PeBoW-WDR55
76,907 particles
3.04 Å
EMD-29169
PeBoW complex
76,907 particles
3.06 Å
EMD-29170
WDR55
76,907 particles
3.50 Å
EMD-29171
Domain III
76,907 particles
2.84 Å
EMD-29173
States E-F
Domain V
203,208 particles
2.58 Å
EMD-29183
Domain VI
203,208 particles
2.58 Å
EMD-29181
DDX18
203,208 particles
2.56 Å
EMD-29182
RRS1
203,208 particles
2.61 Å
EMD-29184
Inter-subunit side
203,208 particles
2.86 Å
EMD-29177
DDX54
203,208 particles
2.79 Å
EMD-29178
NOC3L
203,208 particles
3.07 Å
EMD-29180
L1 Stalk
203,208 particles
2.92 Å
EMD-29179
NOC2L
203,208 particles
3.75 Å
EMD-29185
CryoSparc v3.3.2
Part. Subtr.
Local NU ref.
Mask
Refinement
Starting angles from
Local res. estimation (Å) (FSC-0.5)
Fourier Shell Correlation
Resolution
No Mask
Tight
Corrected

**Fig. S10. Composite cryo-EM reconstruction of states E and F.**

(**A**) Overall and composite cryo-EM maps of state E and state F. For the overall reconstructions, FSC curves (no mask, tight mask, solvent corrected and 3D) and Euler angle distribution are displayed on the left. For the composite maps, the FSC curves (no mask, tight mask, solvent corrected, map-to-model) and a list of the selected focused maps used to generate the composite map are displayed on the left. (**B**) Focused refinements for relevant modules of state E and state F. The mask used for each focus refinement is shown with respect to the reference map. The resulting focused map is colored by local resolution estimation. FSC curves (no mask, tight mask, solvent corrected) are displayed on the right. Software used and type of jobs are indicated. In some cases, when relevant, particles stack from different states are merged to improve the reconstruction of a specific module common to the different states. (**A-B**) The resolution indicated for all FSC curve was determined at FSC-0.143, except for the map-to-model FSC which was determined at FSC-0.5. The Electron Microscopy Data Bank (EMDB) accession number is indicated for each map.

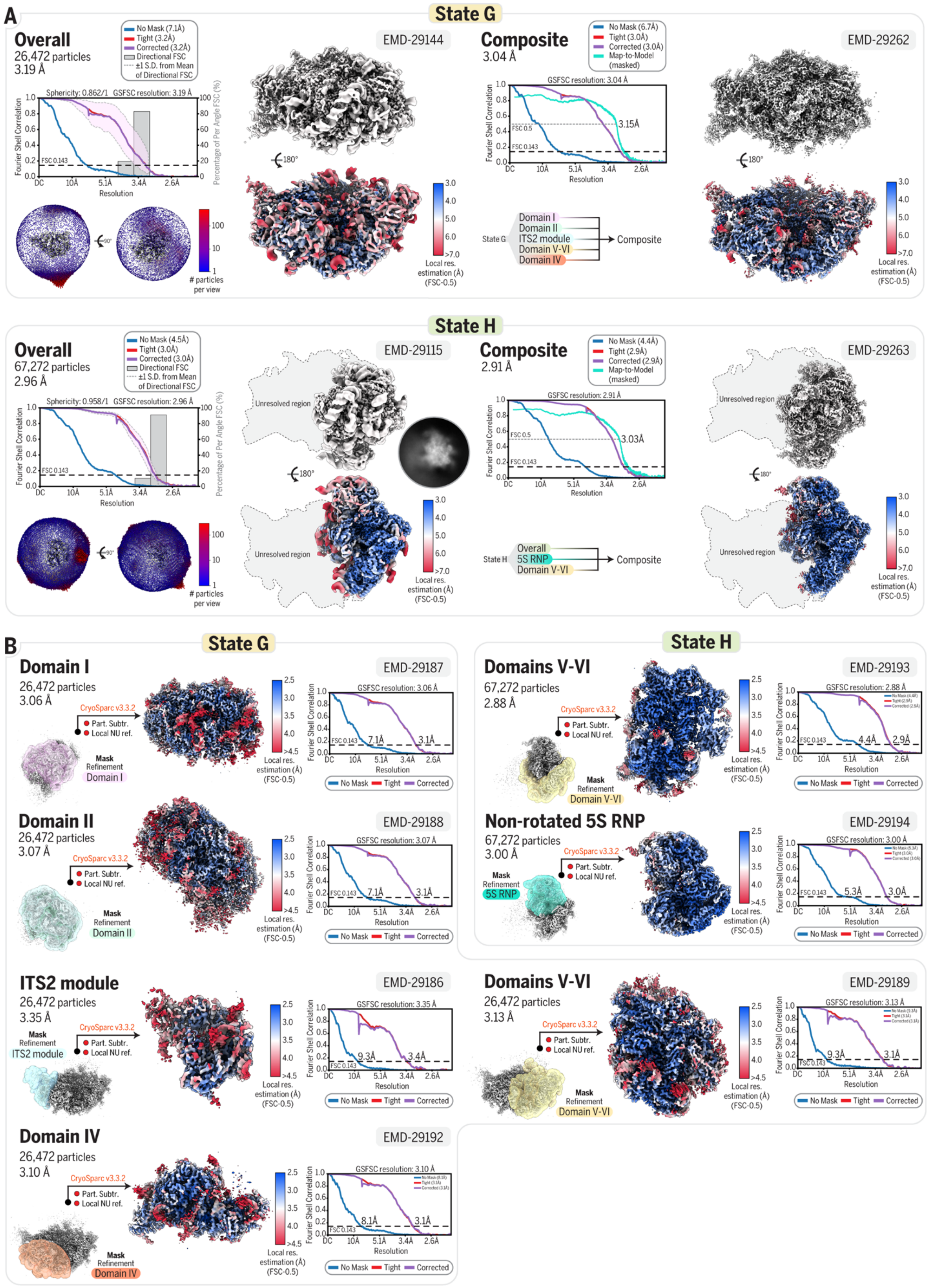

A
State G
Overall
26,472 particles
3.19 Å
EMD-29144
Composite
3.04 Å
EMD-29262
State G
Domain I
Domain II
ITS2 module
Domain V-VI
Domain IV
Composite
State H
Overall
67,272 particles
2.96 Å
EMD-29115
Composite
2.91 Å
EMD-29263
Unresolved region
State H
Overall
5S RNP
Domain V-VI
Composite
B
State G
Domain I
26,472 particles
3.06 Å
EMD-29187
Domain II
26,472 particles
3.07 Å
EMD-29188
ITS2 module
26,472 particles
3.35 Å
EMD-29186
Domain IV
26,472 particles
3.10 Å
EMD-29192
State H
Domains V-VI
67,272 particles
2.88 Å
EMD-29193
Non-rotated 5S RNP
67,272 particles
3.00 Å
EMD-29194
Domains V-VI
26,472 particles
3.13 Å
EMD-29189
CryoSparc v3.3.2
Part. Subtr.
Local NU ref.
Mask Refinement
Local res. estimation (Å) (FSC-0.5)
Fourier Shell Correlation
Resolution
No Mask
Tight
Corrected

**Fig. S11. Composite cryo-EM reconstruction of states G and H.**

(**A**) Overall and composite cryo-EM maps of state G and state H. For the overall reconstructions, FSC curves (no mask, tight mask, solvent corrected and 3D) and Euler angle distribution are displayed on the left. For the composite maps, the FSC curves (no mask, tight mask, solvent corrected, map-to-model) and a list of the selected focused maps used to generate the composite map are displayed on the left. A part of state H could not be resolved (see 2D class). An outline is displayed based on state G to help visualizing state H in context of a full pre-60*S*. (**B**) Focused refinements for relevant modules of state G and state H. The mask used for each focus refinement is shown with respect to the reference map. The resulting focused map is colored by local resolution estimation. FSC curves (no mask, tight mask, solvent corrected) are displayed on the right. Software used and type of jobs are indicated. (**A-B**) The resolution indicated for all FSC curve was determined at FSC-0.143, except for the map-to-model FSC which was determined at FSC-0.5. The Electron Microscopy Data Bank (EMDB) accession number is indicated for each map.

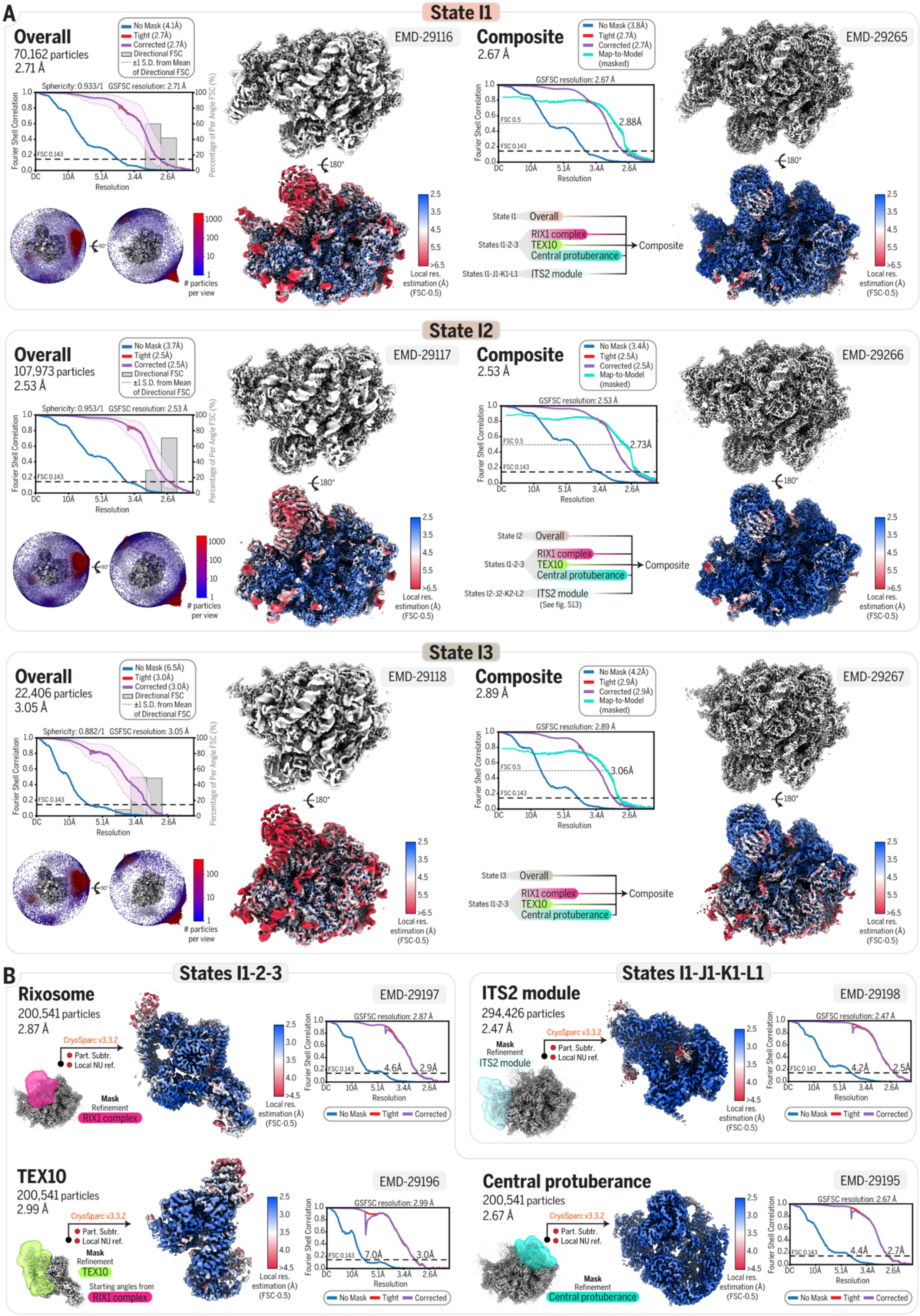
A
State I1
Overall
70,162 particles
2.71 Å
EMD-29116
Composite
2.67 Å
EMD-29265
State I2
Overall
107,973 particles
2.53 Å
EMD-29117
Composite
2.53 Å
EMD-29266
State I3
Overall
22,406 particles
3.05 Å
EMD-29118
Composite
2.89 Å
EMD-29267
B
States I1-2-3
Rixosome
200,541 particles
2.87 Å
EMD-29197
States I1-J1-K1-L1
ITS2 module
294,426 particles
2.47 Å
EMD-29198
TEX10
200,541 particles
2.99 Å
EMD-29196
Central protuberance
200,541 particles
2.67 Å
EMD-29195

**Fig. S12. Composite cryo-EM reconstruction of states I1, I2 and I3.**

(**A**) Overall and composite cryo-EM maps of states I1, I2 and I3. For the overall reconstructions, FSC curves (no mask, tight mask, solvent corrected and 3D) and Euler angle distribution are displayed on the left. For the composite maps, the FSC curves (no mask, tight mask, solvent corrected, map-to-model) and a list of the selected focused maps used to generate the composite map are displayed on the left. (**B**) Focused refinements for relevant modules of states I1, I2 and I3. The mask used for each focus refinement is shown with respect to the reference map. The resulting focused map is colored by local resolution estimation. FSC curves (no mask, tight mask, solvent corrected) are displayed on the right. Software used and type of jobs are indicated. In some cases, when relevant, particles stack from different states are merged to improve the reconstruction of a specific module common to the different states. (**A-B**) The resolution indicated for all FSC curve was determined at FSC-0.143, except for the map-to-model FSC which was determined at FSC-0.5. The Electron Microscopy Data Bank (EMDB) accession number is indicated for each map.

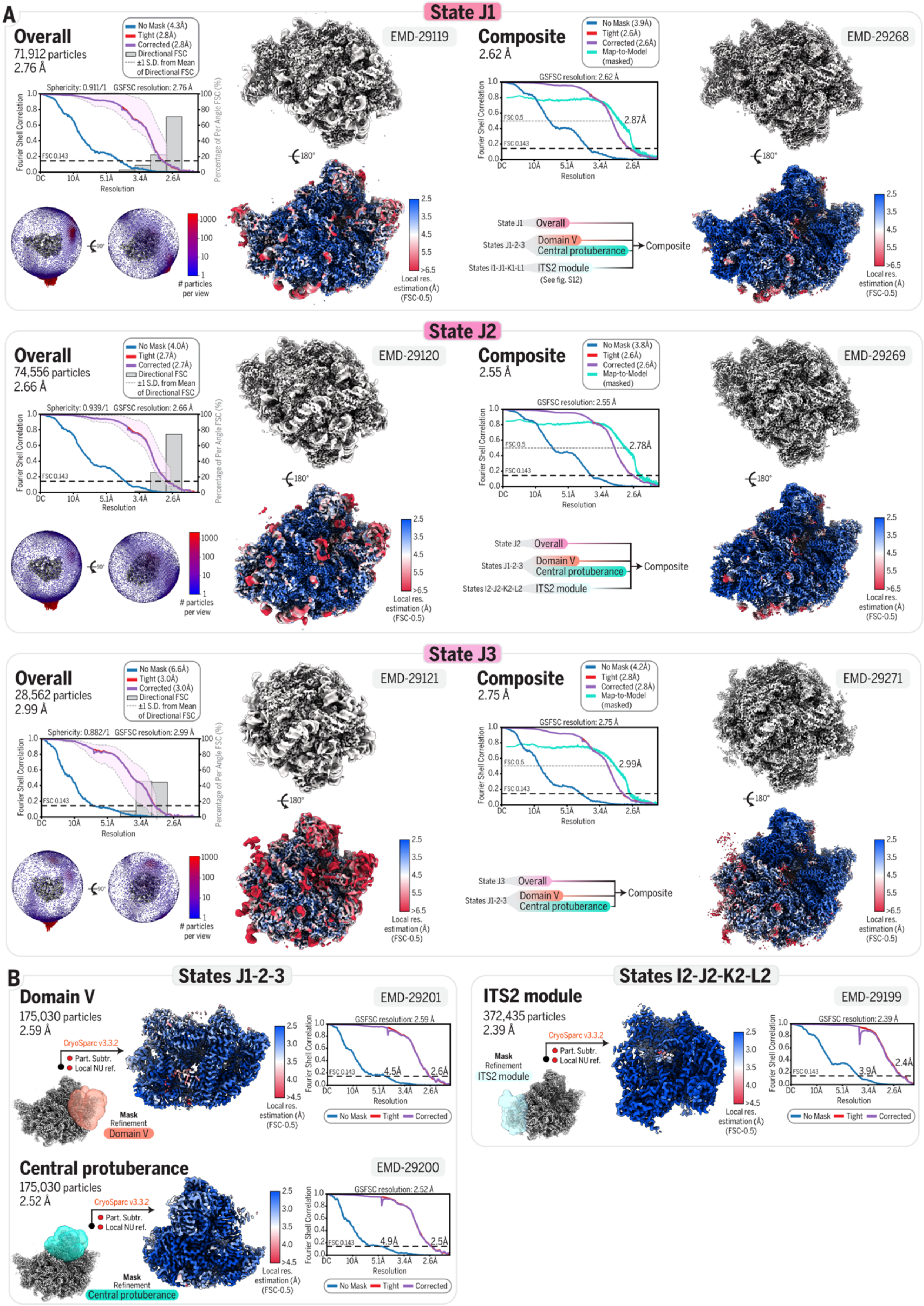

A
State J1
Overall
71,912 particles
2.76 Å
EMD-29119
Composite
2.62 Å
EMD-29268
State J1 Overall
States J1-2-3 Domain V Central protuberance
States I1-J1-K1-L1 ITS2 module
(See fig. S12)
Composite
State J2
Overall
74,556 particles
2.66 Å
EMD-29120
Composite
2.55 Å
EMD-29269
State J2 Overall
States J1-2-3 Domain V Central protuberance
States I2-J2-K2-L2 ITS2 module
Composite
State J3
Overall
28,562 particles
2.99 Å
EMD-29121
Composite
2.75 Å
EMD-29271
State J3 Overall
States J1-2-3 Domain V Central protuberance
Composite
B
States J1-2-3
Domain V
175,030 particles
2.59 Å
EMD-29201
CryoSparc v3.3.2
Part. Subtr.
Local NU ref.
Mask Refinement Domain V
States I2-J2-K2-L2
ITS2 module
372,435 particles
2.39 Å
EMD-29199
Mask Refinement ITS2 module
Central protuberance
175,030 particles
2.52 Å
EMD-29200
Mask Refinement Central protuberance
No Mask
Tight
Corrected

**Fig. S13. Composite cryo-EM reconstruction of states J1, J2 and J3.**

(**A**) Overall and composite cryo-EM maps of states J1, J2 and J3. For the overall reconstructions, FSC curves (no mask, tight mask, solvent corrected and 3D) and Euler angle distribution are displayed on the left. For the composite maps, the FSC curves (no mask, tight mask, solvent corrected, map-to-model) and a list of the selected focused maps used to generate the composite map are displayed on the left. (**B**) Focused refinements for relevant modules of states J1, J2 and J3. The mask used for each focus refinement is shown with respect to the reference map. The resulting focused map is colored by local resolution estimation. FSC curves (no mask, tight mask, solvent corrected) are displayed on the right. Software used and type of jobs are indicated. In some cases, when relevant, particles stack from different states are merged to improve the reconstruction of a specific module common to the different states. (**A-B**) The resolution indicated for all FSC curve was determined at FSC-0.143, except for the map-to-model FSC which was determined at FSC-0.5. The Electron Microscopy Data Bank (EMDB) accession number is indicated for each map.

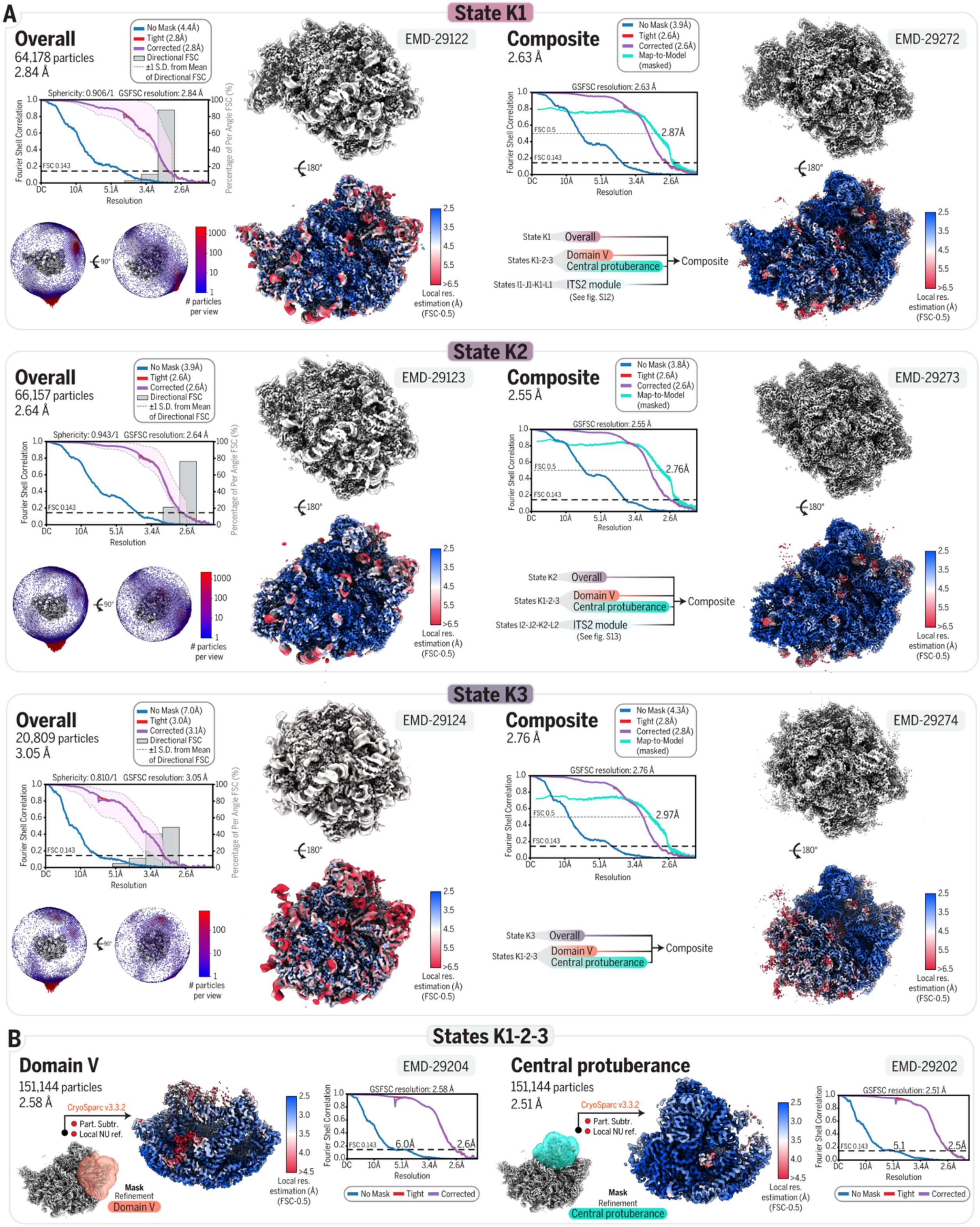
A
State K1
Overall
64,178 particles
2.84 Å
EMD-29122
Composite
2.63 Å
EMD-29272
State K2
Overall
66,157 particles
2.64 Å
EMD-29123
Composite
2.55 Å
EMD-29273
State K3
Overall
20,809 particles
3.05 Å
EMD-29124
Composite
2.76 Å
EMD-29274
B
States K1-2-3
Domain V
151,144 particles
2.58 Å
EMD-29204
Central protuberance
151,144 particles
2.51 Å
EMD-29202

**Fig. S14. Composite cryo-EM reconstruction of states K1, K2 and K3.**

(**A**) Overall and composite cryo-EM maps of states K1, K2 and K3. For the overall reconstructions, FSC curves (no mask, tight mask, solvent corrected and 3D) and Euler angle distribution are displayed on the left. For the composite maps, the FSC curves (no mask, tight mask, solvent corrected, map-to-model) and a list of the selected focused maps used to generate the composite map are displayed on the left. (**B**) Focused refinements for relevant modules of states K1, K2 and K3. The mask used for each focus refinement is shown with respect to the reference map. The resulting focused map is colored by local resolution estimation. FSC curves (no mask, tight mask, solvent corrected) are displayed on the right. Software used and type of jobs are indicated. In some cases, when relevant, particles stack from different states are merged to improve the reconstruction of a specific module common to the different states. (**A-B**) The resolution indicated for all FSC curve was determined at FSC-0.143, except for the map-to-model FSC which was determined at FSC-0.5. The Electron Microscopy Data Bank (EMDB) accession number is indicated for each map.

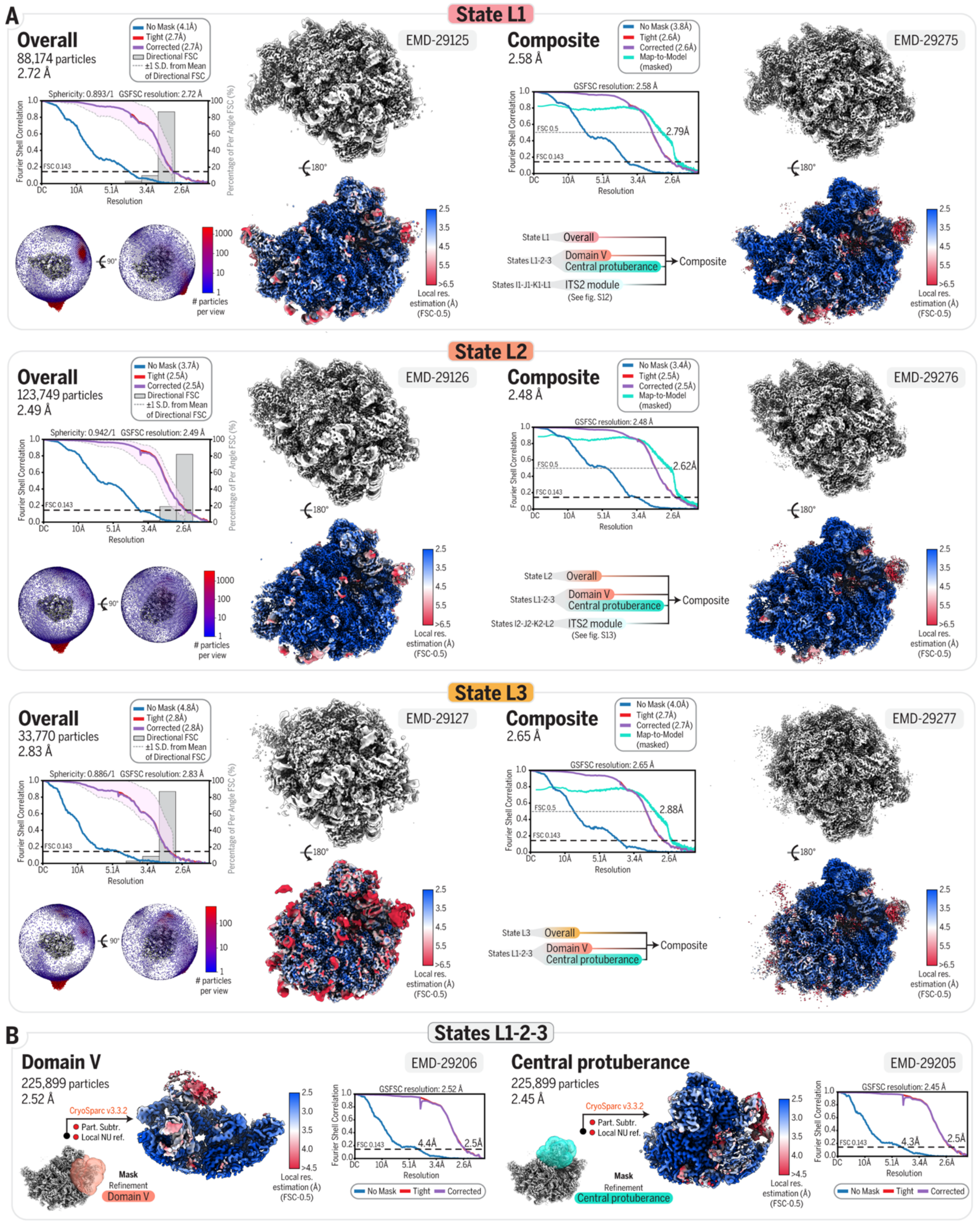
A
State L1
Overall
88,174 particles
2.72 Å
EMD-29125
Composite
2.58 Å
EMD-29275
State L2
Overall
123,749 particles
2.49 Å
EMD-29126
Composite
2.48 Å
EMD-29276
State L3
Overall
33,770 particles
2.83 Å
EMD-29127
Composite
2.65 Å
EMD-29277
B
States L1-2-3
Domain V
225,899 particles
2.52 Å
EMD-29206
Central protuberance
225,899 particles
2.45 Å
EMD-29205

**Fig. S15. Composite cryo-EM reconstruction of states L1, L2 and L3.**

(**A**) Overall and composite cryo-EM maps of states L1, L2 and L3. For the overall reconstructions, FSC curves (no mask, tight mask, solvent corrected and 3D) and Euler angle distribution are displayed on the left. For the composite maps, the FSC curves (no mask, tight mask, solvent corrected, map-to-model) and a list of the selected focused maps used to generate the composite map are displayed on the left. (**B**) Focused refinements for relevant modules of states L1, L2 and L3. The mask used for each focus refinement is shown with respect to the reference map. The resulting focused map is colored by local resolution estimation. FSC curves (no mask, tight mask, solvent corrected) are displayed on the right. Software used and type of jobs are indicated. In some cases, when relevant, particles stack from different states are merged to improve the reconstruction of a specific module common to the different states. (**A-B**) The resolution indicated for all FSC curve was determined at FSC-0.143, except for the map-to-model FSC which was determined at FSC-0.5. The Electron Microscopy Data Bank (EMDB) accession number is indicated for each map.

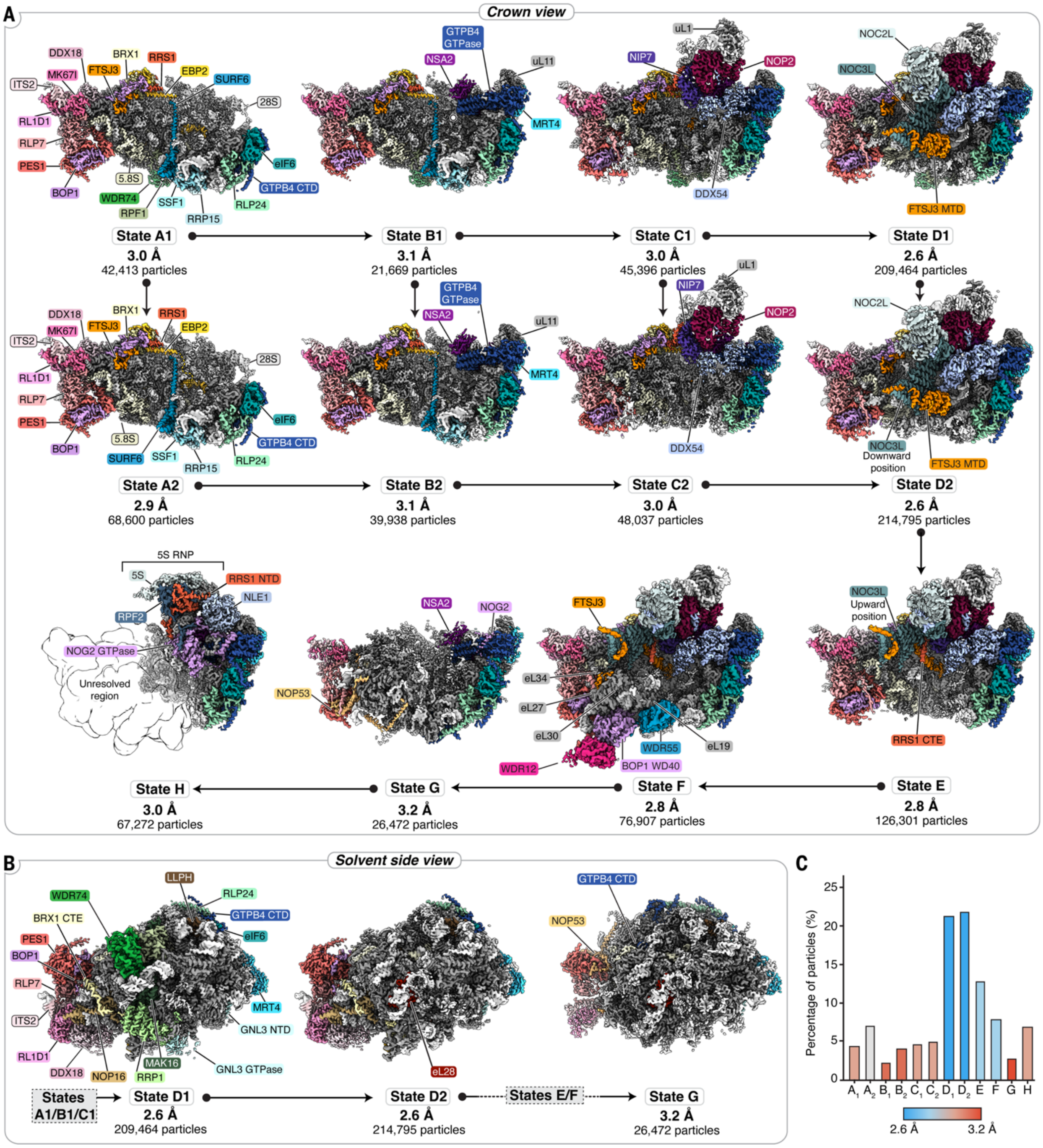


**Fig. S16. Cryo-EM structures of human pre-60*S* nucleolar assembly intermediates.**

(**A**) Crown view of cryo-EM maps of twelve nucleolar states A1 to H. Particles number used for each reconstruction and resolution are indicated below each state. (**B**) Solvent side view of states D1, D2 and G highlighting key transitions on the back of the pre-60*S* particles. (**C**) Graph showing the percentage of particles contributing to each state within the pool of nucleolar assembly intermediates.

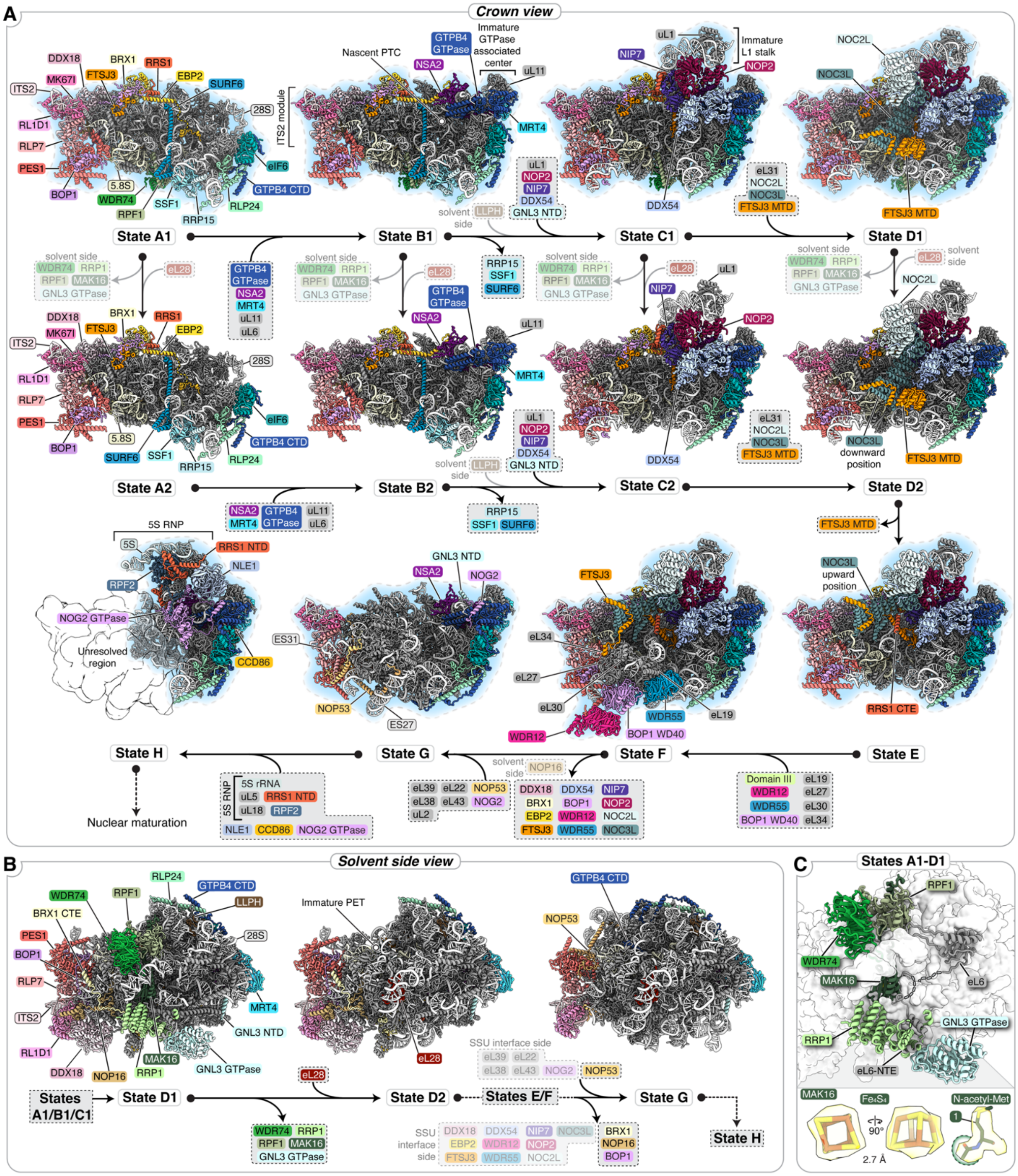


**Fig. S17. Detailed nucleolar assembly of human pre-60*S* assembly intermediates.**

(**A**) Crown view of atomic models of twelve nucleolar states A1 to H with ribosomal proteins in grey and color-coded pre-rRNAs and ribosome assembly factors. Proteins that become ordered or depart between different assembly states are indicated on arrows. Parallel pathways of assembly are illustrated for states A1-D1 and A2-D2. (**B**) Solvent side view of states D1, D2 and, G highlighting transitions on the back of the pre-60*S* particles. (**C**) Detailed view of assembly factors at the solvent-exposed side. Insets show cryo-EM maps around an iron-sulfur cluster and a post-translational modification. Models highlighted in blue were used to illustrate the main nucleolar pathway (states A-H) in Figure 1.

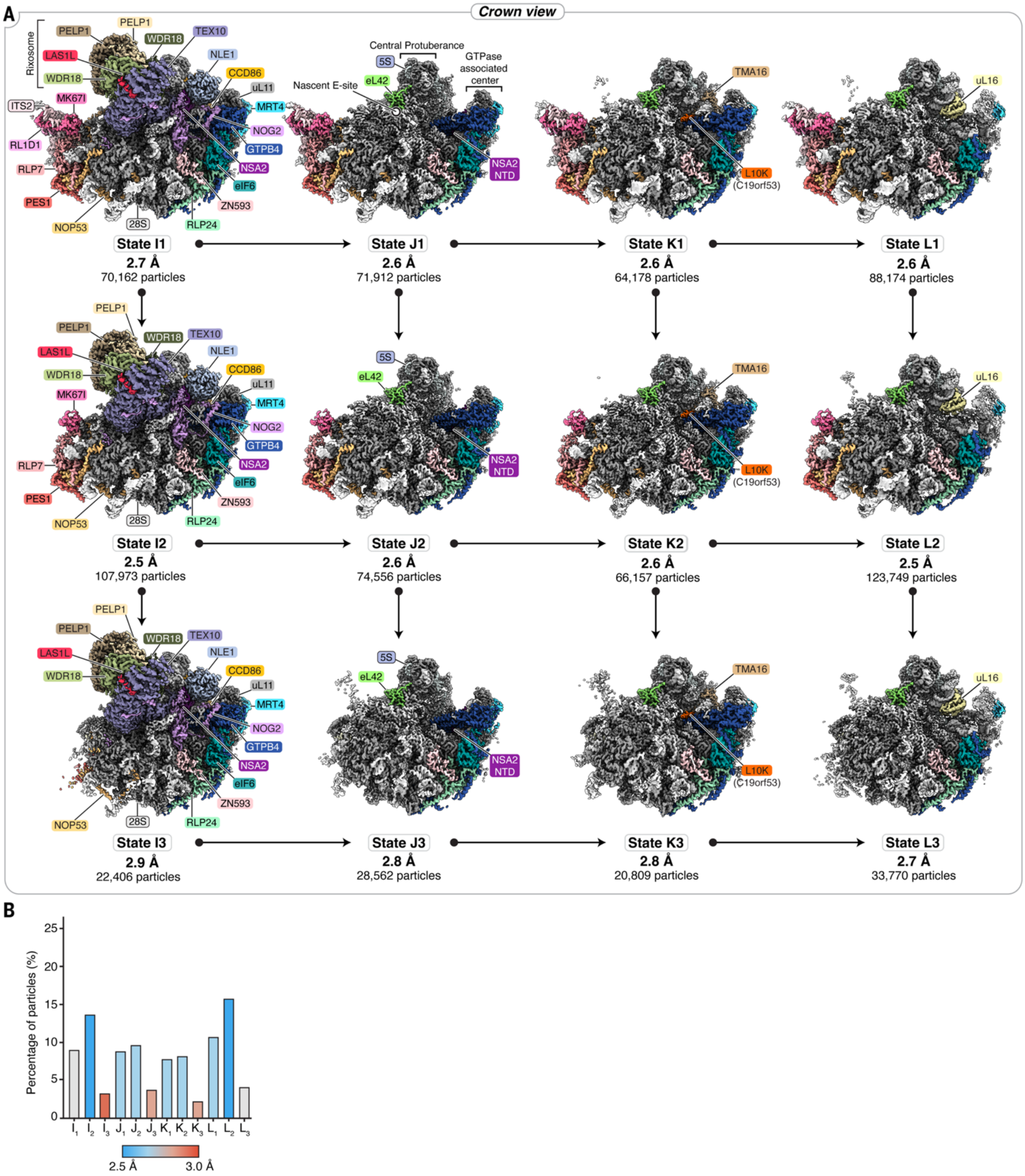


**Fig. S18. Cryo-EM structures of human pre-60*S* nuclear assembly intermediates.**

(**A**) Crown view of cryo-EM maps of twelve nuclear states I1 to L3. Particles number used for each reconstruction and resolution are indicated below each state. (**B**) Graph showing the percentage of particles contributing to each state within the pool of nuclear assembly intermediates.

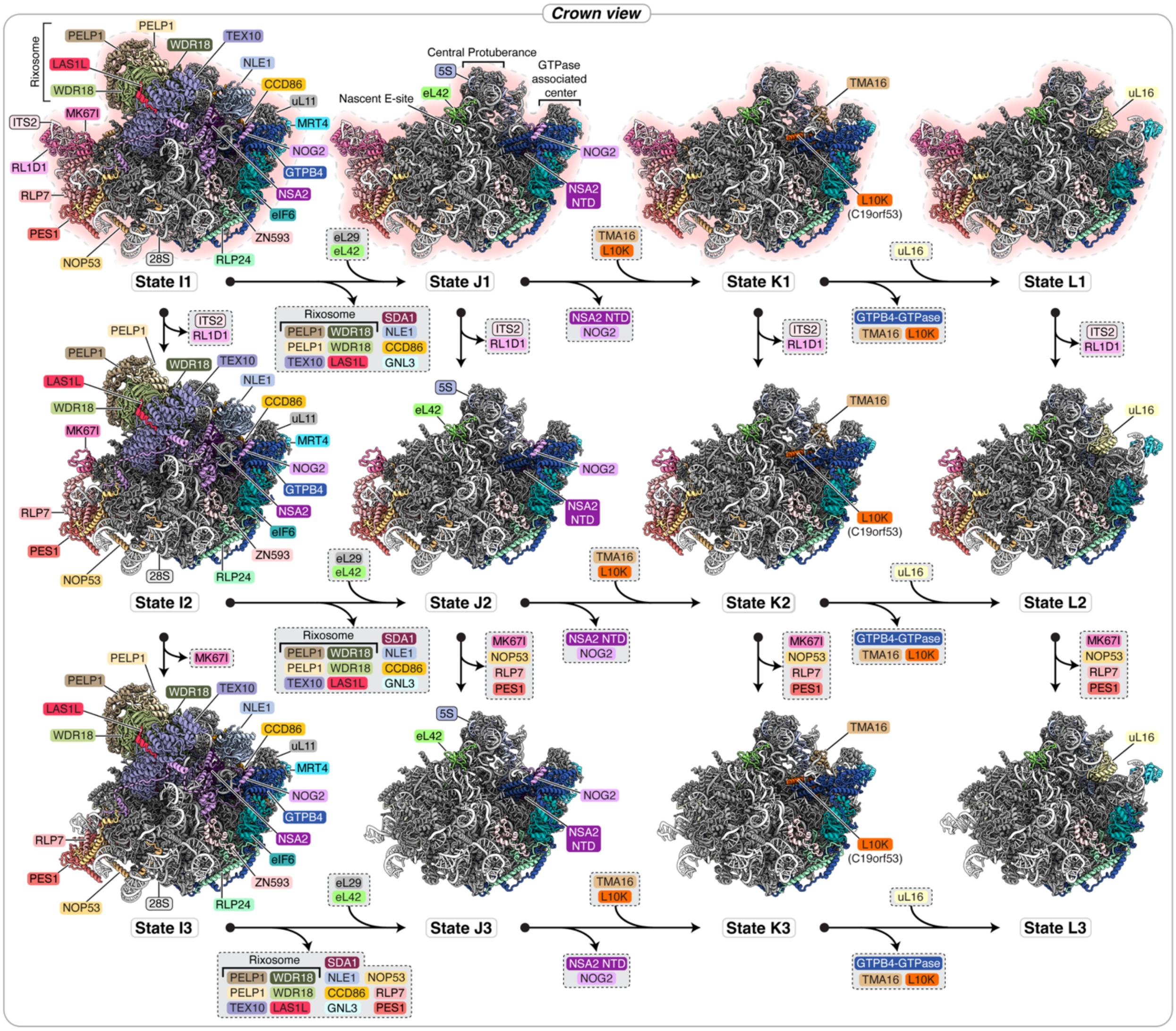


**Fig. S19. Parallel nuclear maturation pathways of human pre-60*S* assembly intermediates.** Crown view of atomic models of twelve nuclear states I1 to L3 with 28*S* rRNA colored in white, ITS2 rRNA in pink, 5*S* rRNA in blue, ribosomal proteins in grey and color-coded ribosome assembly factors. Proteins that become ordered or depart between different assembly states are indicated on arrows. Models highlighted in red were used to illustrate the main nuclear pathway (states I-L) in Figure 1.

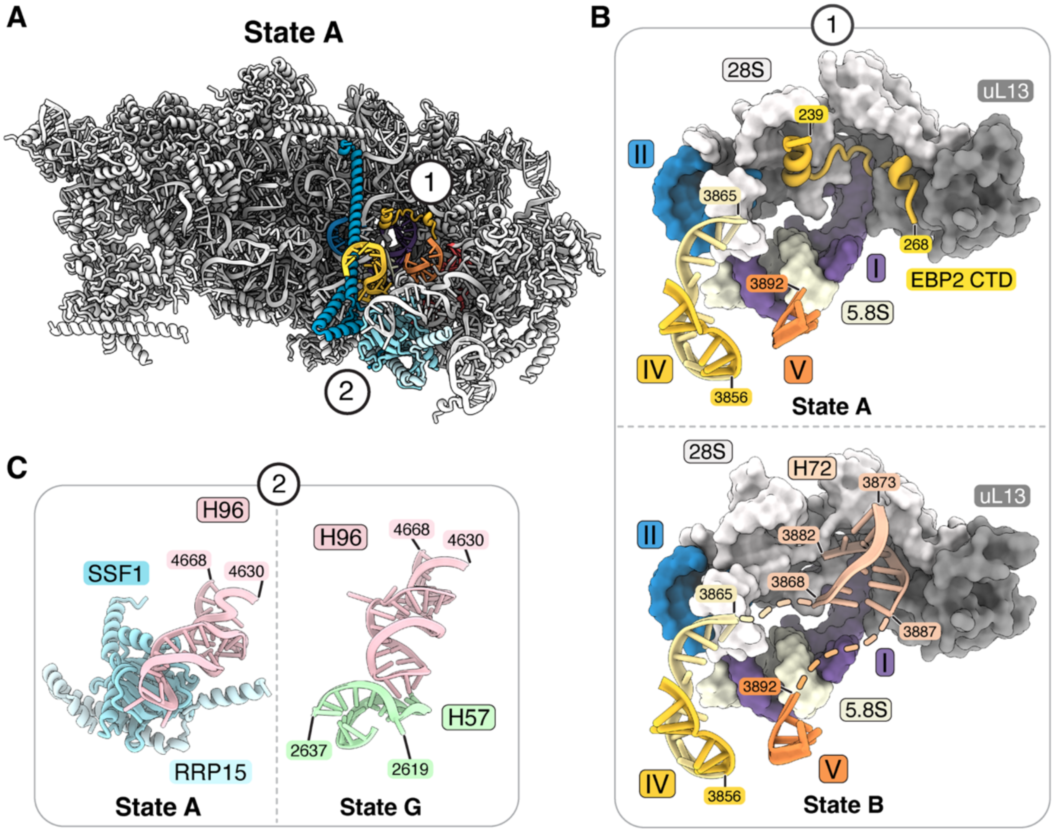


**Fig. S20. Ribosome assembly factors prevent premature rRNA folding.**

(**A**) Crown view of state A with color-coded rRNA root helices and the SURF6-SSF1-RRP15 complex shown in shades of blue. Numbers indicate regions with detailed views in other panels. (**B**) Root helices of domains IV and V near EBP2 in state A (top) compared with state B where EBP2 is replaced by helix 72 (H72) (bottom). (**C**) The SSF1-RRP15 complex binds to H96 in state A (left) whereas H57 interacts with the same region of H96 in state G (right).

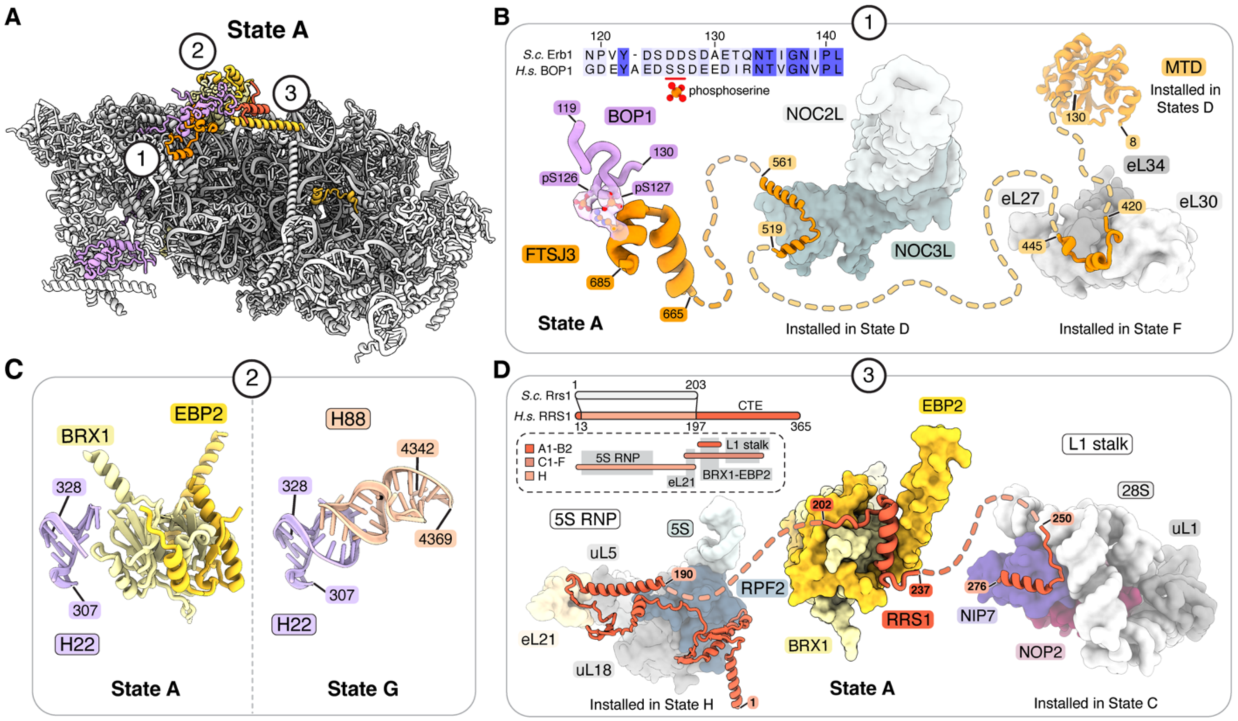


**Fig. S21. Functions of the protein interaction hub during human pre-60*S* biogenesis.**

(**A**) Crown view of state A highlighting a protein-protein interaction hub with color-coded assembly factors. Numbers indicate regions with detailed views in other panels. (**B**) Functions of FTSJ3 peptides as seen in state A (left), state D (center) and state F (right). The locations of phosphoserines in an FTSJ3-interacting BOP1 peptide are indicated in a multiple sequence alignment of yeast Erb1 and human BOP1 and highlighted with corresponding cryo-EM density maps on the left. (**C**) BRX1-EBP2 bind H22 in state A (left) whereas, in state G, H22 interacts with H88 (right). (**D**) Functions of RRS1 as seen in state A with EBP2-BRX1 (center), state H with 5*S* RNP (left) and the L1 stalk in state C (right). A schematic highlights the C-terminal extension (CTE) of human RRS1 and the elements involved in interacting with other components in states A1-B2, C1-F and H.

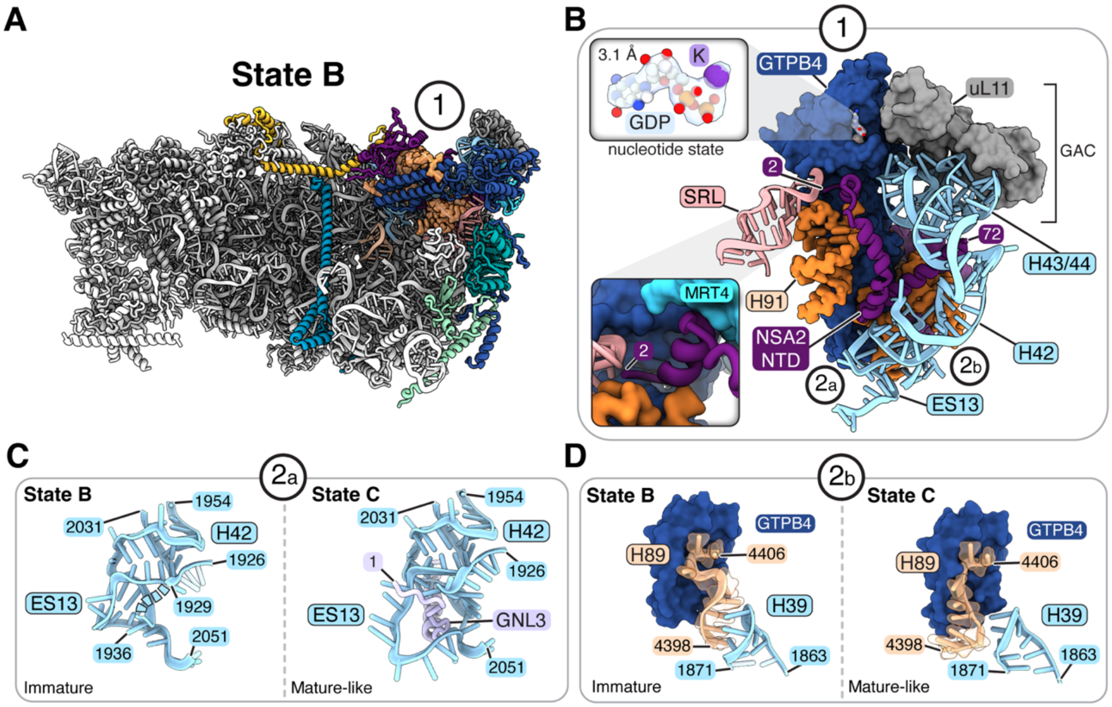


**Fig. S22. A central role for GTPB4 in early PTC installation.**

(**A**) Crown view of state B showing selected assembly factors. Assembly factors are color-coded and newly integrated PTC rRNA represented as orange surface. Numbers indicate regions with detailed views in other panels. (**B**) Detailed view of GDP-bound GTPB4 and its interacting proteins and color-coded rRNA elements from domains II, V and VI, including the sarcin-ricin loop (SRL). (**C**) Detailed views of pre-rRNA elements of domain II in an immature conformation in state B (left) or in complex with the N-terminus of GNL3 in state C (right). (**D**) Detailed views of elements of the PTC in the immature conformation in state B (left) or in a mature-like conformation in state C (bottom).

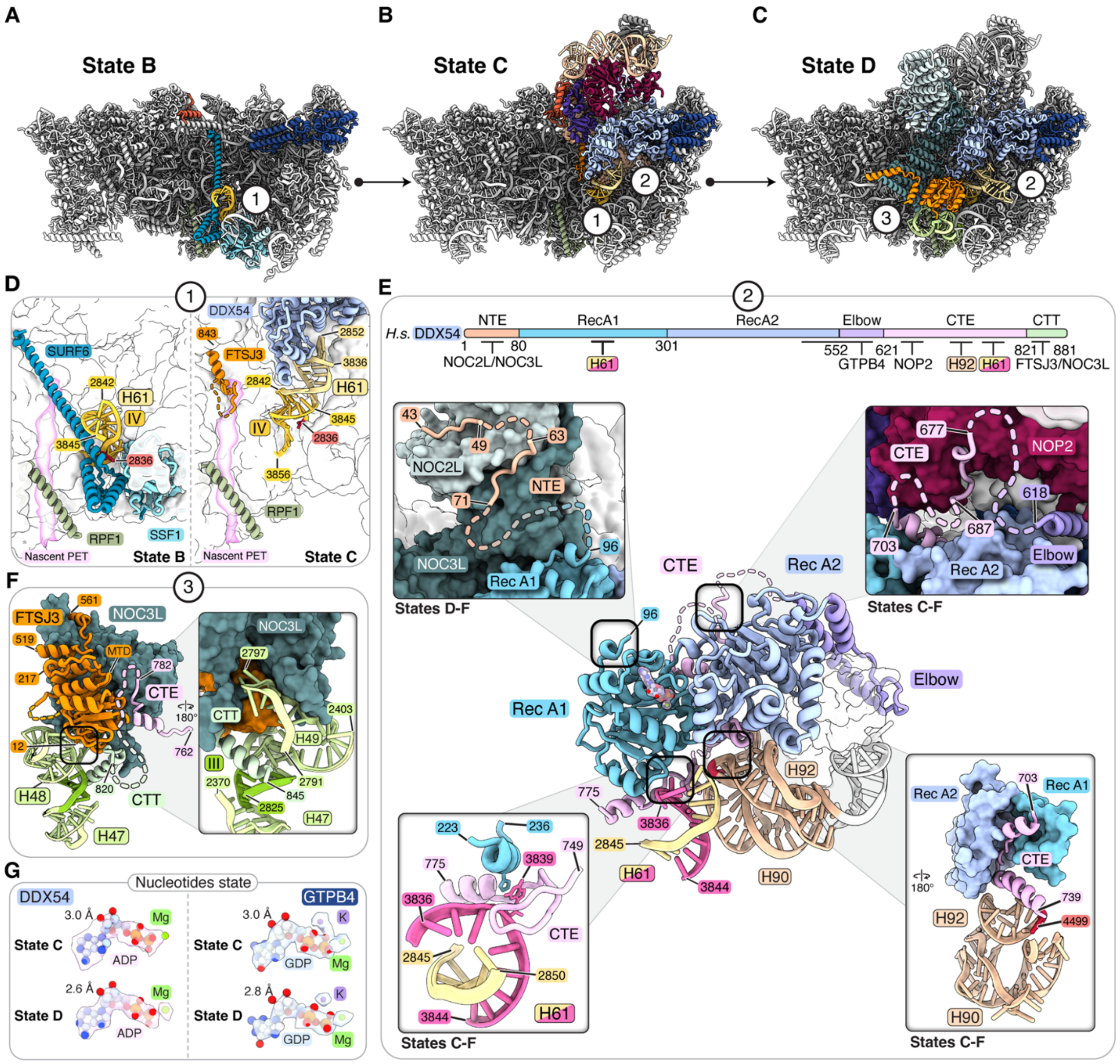


**Fig. S23. Critical roles of DDX54 during nucleolar pre-60*S* assembly.**

(**A-C**) Crown views of states B (A), C (B) and D (C) showing selected assembly factors as they enter the respective states. Numbers indicate regions with detailed views in other panels. (**D**) Root helix IV and H61 shown before (state B, left) and after (state C, right) remodeling by DDX54. The occluded nascent PET (pink) is shown with RPF1 (green) and FTSJ3 (orange). (**E**) Interactions of DDX54 in states C-F. A schematic indicates the domain organization and interaction partners of DDX54. The cartoon representation of DDX54 in context of PTC elements (orange) and helix 61 (H61, yellow/pink) shows key interactions with insets highlighting interactions of the N-terminus (top left), the elbow and C-terminal extension (CTE; top right). Key RNA interactions of DDX54 are shown in the bottom insets with H61 interactions (bottom left) and PTC interactions (bottom right). (**F**) Interactions between the FTSJ3 methyltransferase domain with NOC3L and the C-terminal extension (CTE) and C-terminal tail (CTT) of DDX54. (**G**) Nucleotide state of DDX54 (left) and GTPB4 (right) in state C (top) and state D (bottom).

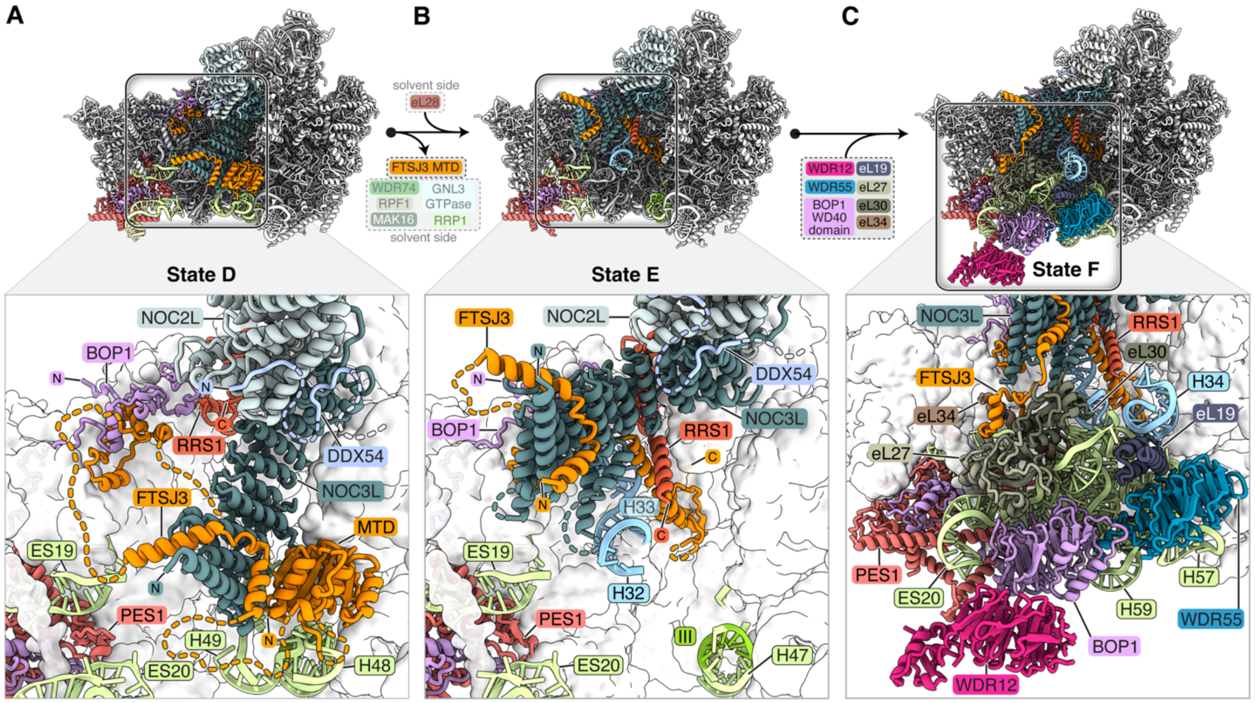


**Fig. S24. WD40 proteins chaperone the integration of domain III.**

(**A-C**) Crown views of states D (A), E (B), and F (C) showing selected assembly factors involved in domain III remodeling with the NOC2L-NOC3L complex adopting a downward conformation in state D (A) and an upward conformation in state E (B). Resulting changes in domain III rRNA elements shown in light green are completed with the formation of state F (C). Zoomed views depict changes during the integration of domain III.

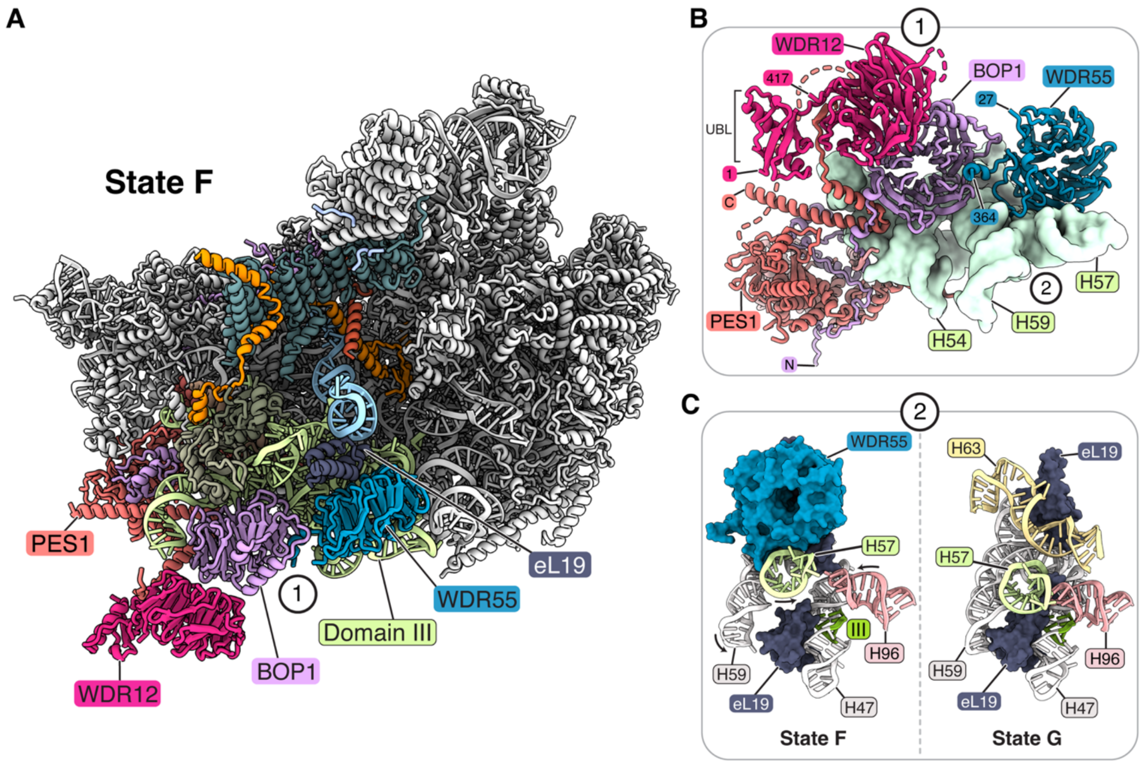


**Fig. S25. Chaperoning of domain III in state F.**

(**A**) Architecture of the fully stabilized domain III rRNA in state F with color-coded assembly factors involved. (**B**) Detailed view depicting the roles of WD40 repeat proteins involved in stabilizing domain III. (**C**) Comparison of pre-rRNA elements chaperoned and separated by WDR55 in state F (left) and interacting in state G (right).

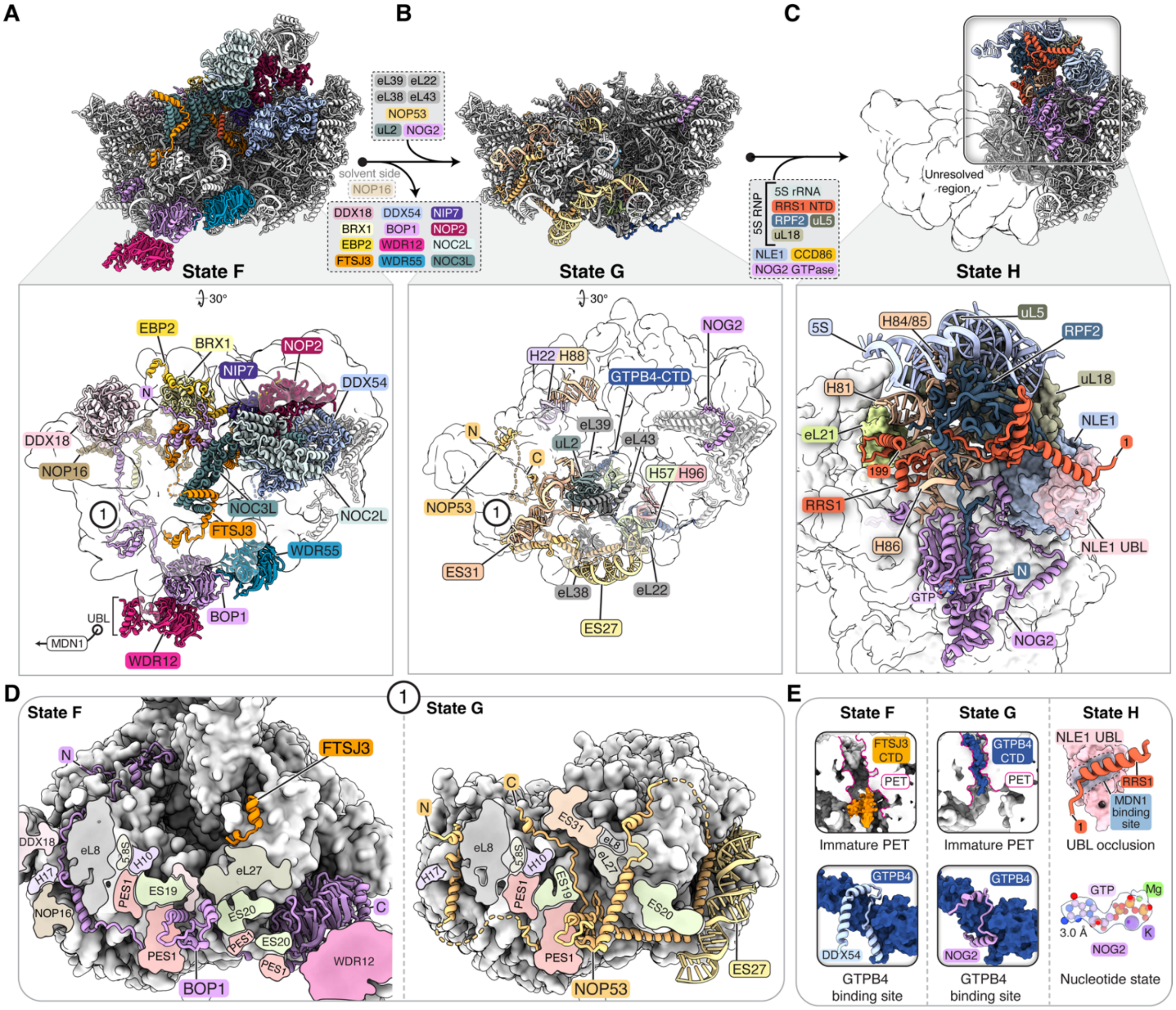


**Fig. S26. Irreversible restructuring during late nucleolar assembly primes nuclear maturation.**

(**A-C**) Crown views of state F (A), state G (B) and the partial reconstruction of state H (C) showing protein and RNA elements that are either removed from state F (A) or incorporated in state G (B) or state H (C). Zoomed views below highlight a protein-protein interaction network of factors in state F that is removed by MDN1 (A), new rRNA elements, ribosomal proteins and NOP53 observed in state G (B), and the installation of the 5*S* RNP in state H (C). (**D**) Slab views of particles with BOP1 and interacting elements in state F (left) and NOP53 and interacting elements in state G (right). (**E**) Features of states F (left) and state G (center) regarding the polypeptide exit tunnel (top) and proteins associated with GTPB4 (bottom). Features of state H (right) showing the binding of RRS1 to NLE1 that blocks MDN1 engagement (top) and the nucleotide state of NOG2 (bottom).

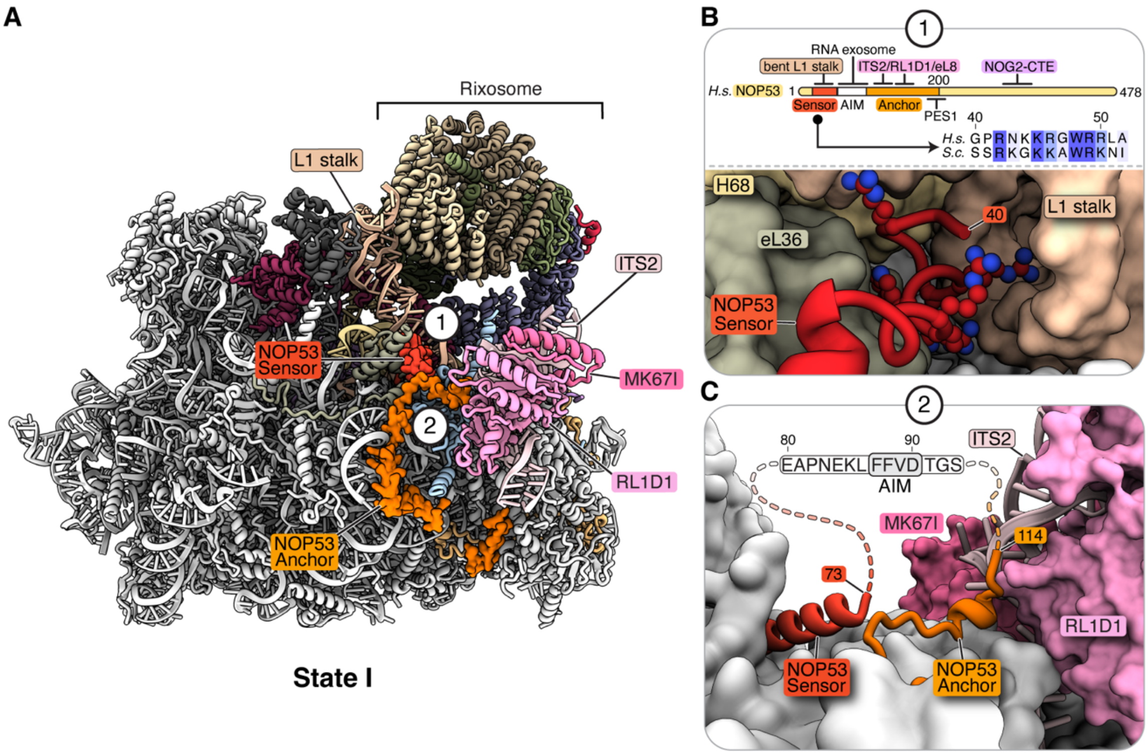


**Fig. S27. NOP53-mediated recognition of state I1.**

(**A**) View of state I1 showing selected assembly factors involved in ITS2 processing with ITS2-associated factors shown in shades of pink and NOP53 in red, orange and yellow. Numbers indicate regions with detailed views in other panels. (**B**) Schematic of the domain architecture of NOP53, its interaction partners, and the conserved N-terminal sensor domain. A zoomed view below shows the interactions of the sensor domain near the L1 stalk. Conserved residues are shown as sticks. (**C**) Detailed view of the NOP53 sensor and anchor domains near ITS2. The approximate position of the exosome recruiting arch interacting motif (AIM) is shown.

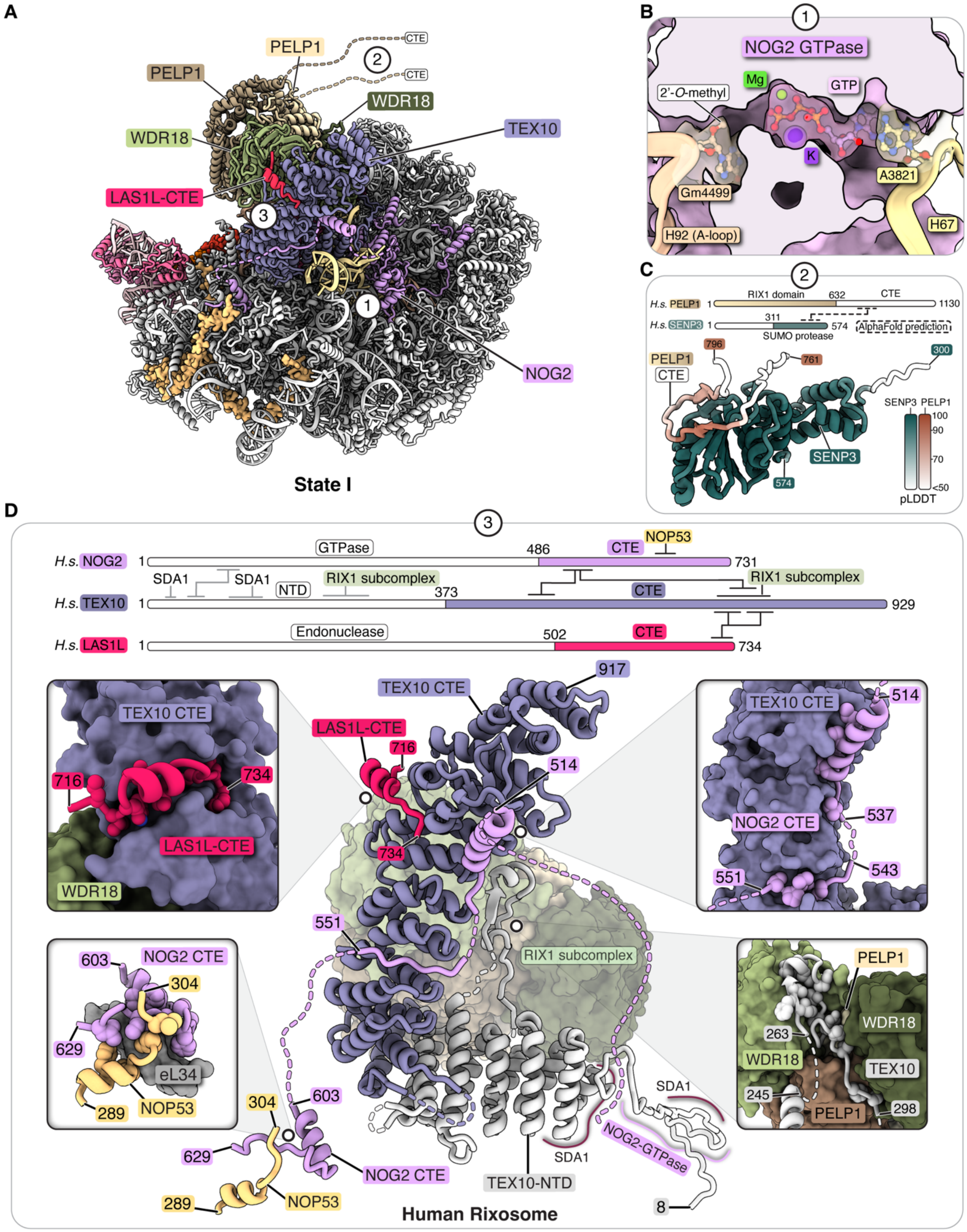


**Fig. S28. Functional architecture of the human rixosome**

(**A**) Crown view of state I with color-coded assembly factors involved in ITS2 processing. Numbers indicate regions with detailed views in other panels. (**B**) Detailed view of the NOG2

GTPase active site with flanking bases from the A-loop (Gm4499) and H67 (A3821). (**C**) AlphaFold-Multimer prediction of the interaction of a peptide of PELP1 with SENP3. Confidence levels (pLDDT) are color coded for each protein. (**D**) Schematic and detailed view showing the higher order organization of the human rixosome with C-terminal extensions (CTE). Insets show specific interactions involving LAS1L (top left), NOG2 and NOP53 (bottom left), TEX10 and NOG2 (top right) and WDR18 and TEX10 (bottom right).

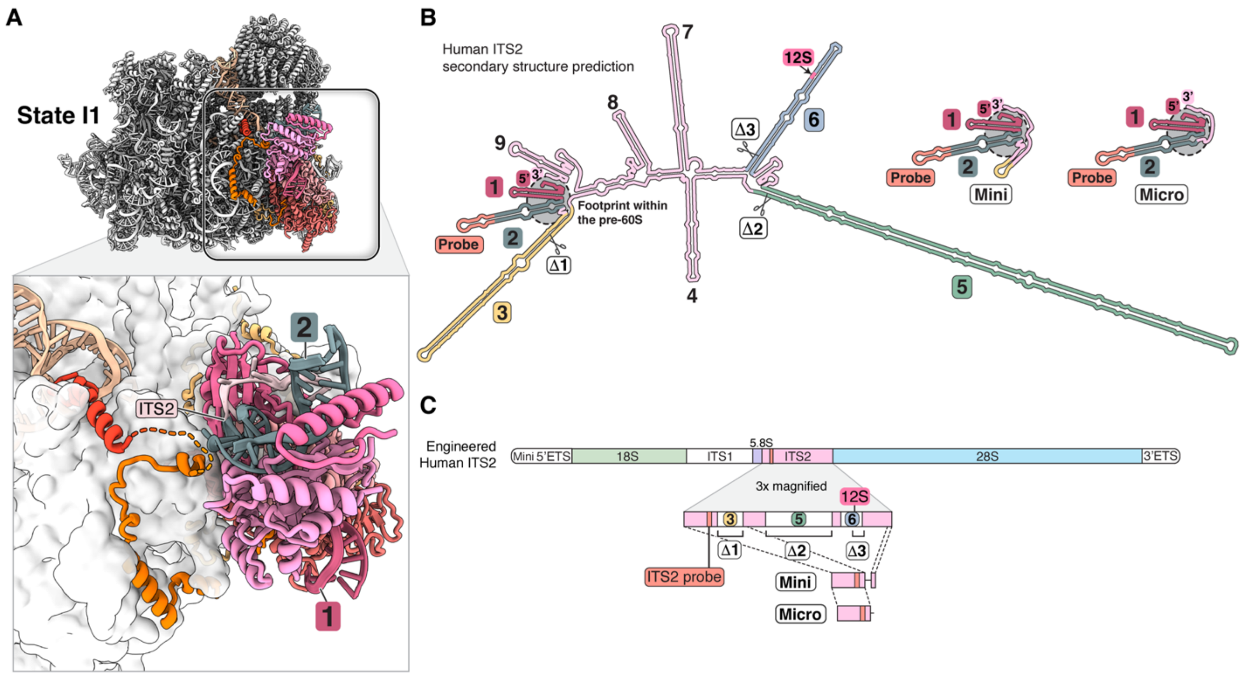


**Fig. S29. Secondary structure prediction and engineered human ITS2 variants.**

(**A**) View of state I1 with color coded assembly factors near ITS2. The zoomed view below shows domains of NOP53 around ITS2-associated assembly factors and color-coded RNA helices 1 and 2 of ITS2. (**B**) Secondary structure predictions of engineered ITS2 variants, including a wildtype (left), and two smaller designs (Mini & Micro) (center and right, respectively). ITS2 helices (1-9), designed truncations (Δ1-3), an inserted ITS2 probe for Northern blotting (Probe), the predicted LAS1L cleavage site (12S), and the footprint within the pre-60*S* particle are indicated. (**C**) Schematic representation of the locations of ITS2 truncations shown in (B) in relation to an engineered human rDNA locus. Zoomed sections indicate the position of ITS2 variants (bottom).

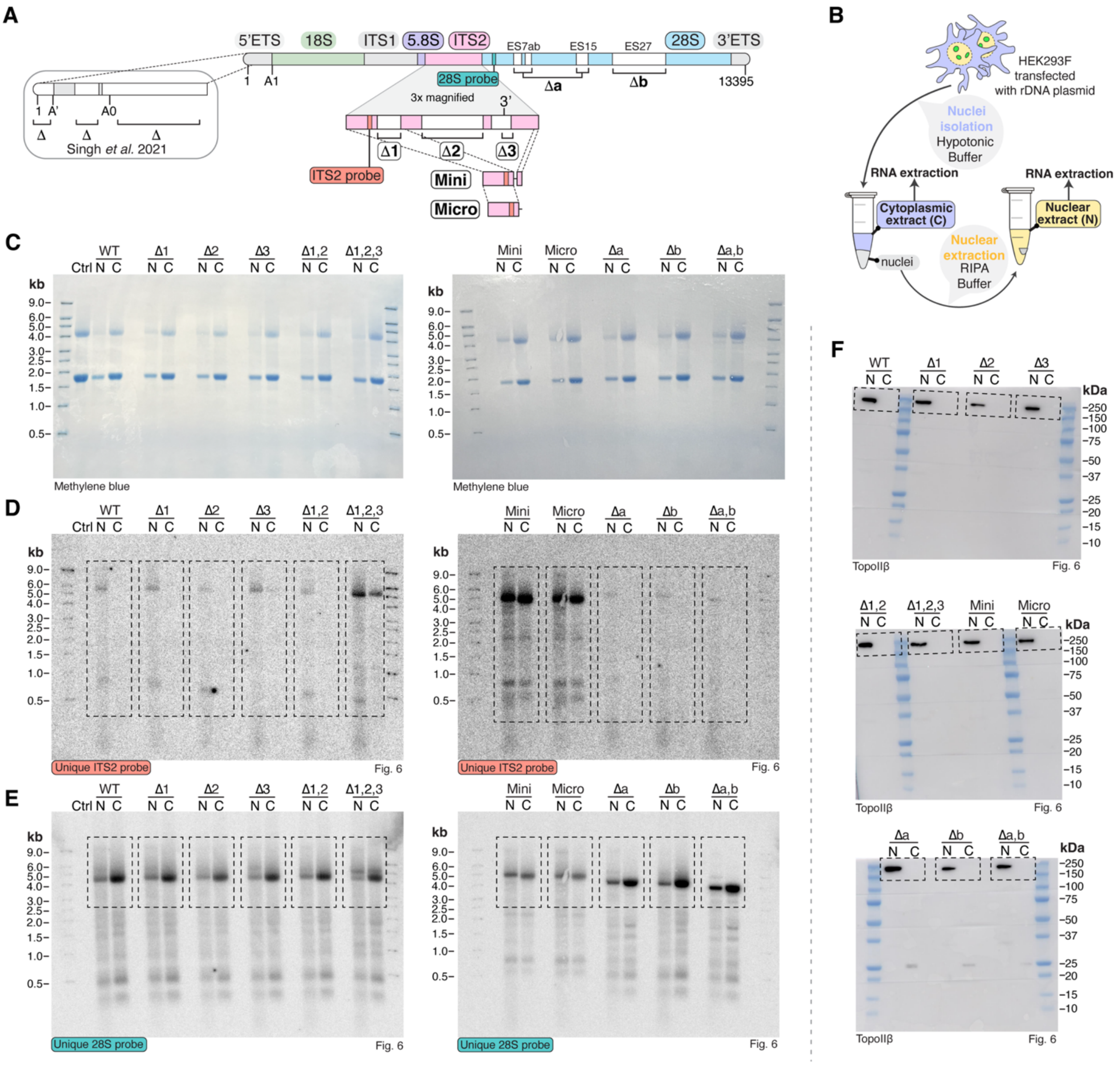


#### Fig. S30. Functional studies of recombinant human rDNA loci.

(**A**) Schematic representation of a human ribosomal RNA transcript showing 3 truncations (Δ1, Δ2, Δ3) within the internal transcribed spacer 2 (ITS2) and 2 truncations (Δa and Δb) in the 28*S*. Unique sequences inserted in the ITS2 and 28*S* transcripts can be recognized by specific ITS2 probe and 28*S* probe, respectively. All the rDNA constructs contain a miniaturized 5'ETS which was shown to have no impact on the LSU assembly (*32*). (**B**) Schematic illustration of the biochemical procedures used to obtain cytoplasmic and nuclear extracts for Northern blotting shown in (D) and (E). (**C**) Methylene blue staining showing 18*S* and 28*S* RNA bands detected from a non-transfected control (ctrl), or cells transfected with a wild-type (ITS2 and 28*S* WT) plasmid, or plasmids harboring truncations (Δ1; Δ2; Δ3; Δ1,2; Δ1,2,3; Mini; Micro; Δa; Δb and Δa,b). (**D**) Northern blot analysis using the unique ITS2 probe showing bands corresponding to the intermediate 32*S* pre-rRNA and different sizes of its cleaved product. (**E**) Northern blot analysis using the unique 28*S* probe showing bands corresponding to the mature 28*S* rRNA and its precursor 32*S* pre-rRNA. (**F**) Western blot analysis of the nuclear and cytoplasmic protein content prior to the total RNA extraction. Antibodies against TopoIIβ, a nuclear marker, were used to demonstrate that the nuclear content has not leaked into the cytoplasm during the experiment. The dashed squares show the cropped regions used in Fig. 6.

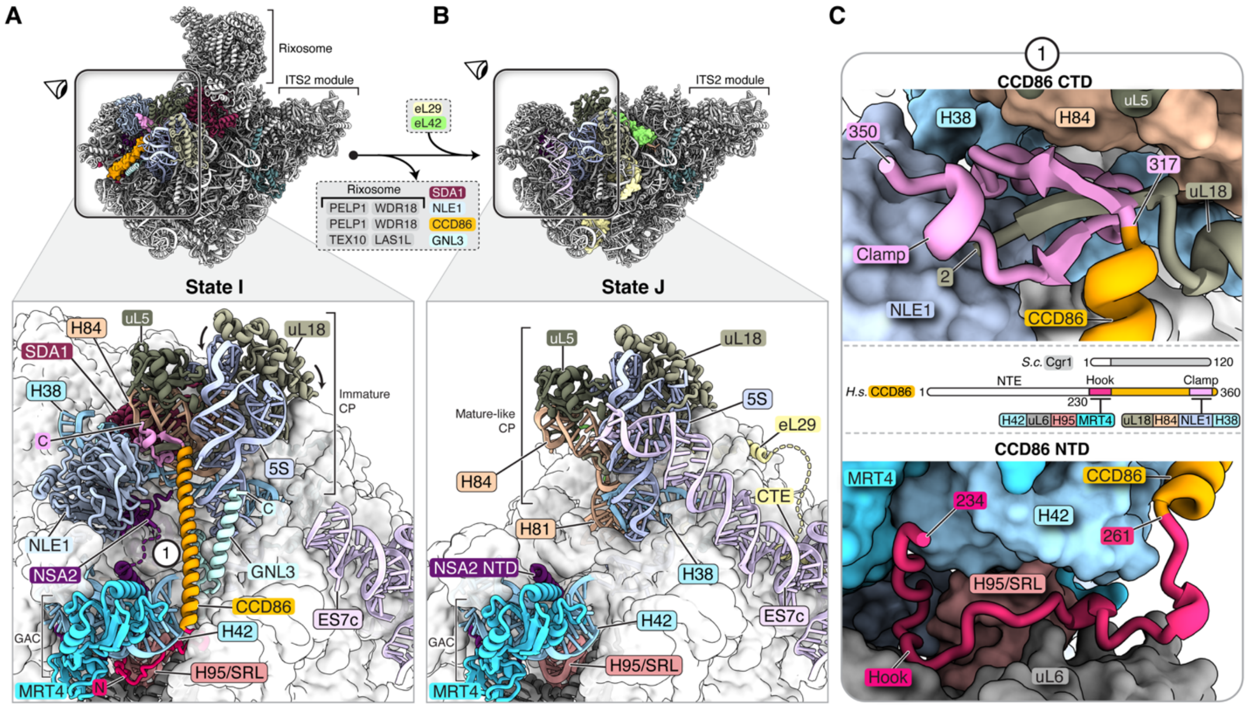


**Fig. S31. Formation of the central protuberance upon rixosome departure.**

(**A**) Top view of state I with color-coded assembly factors near the central protuberance (CP). The zoomed and rotated view below shows assembly factors including the three-domain assembly factor CCD86 (red-orange-pink) stabilizing the immature CP. (**B**) Top view of state J with color-coded assembly factors and ribosomal proteins eL29 (yellow) and eL42 (green) near the central protuberance (CP). The zoomed and rotated view below highlights the consequences of assembly factor removal from state I to J. (**C**) C-terminal domain interactions of CCD86 with NLE1 (top), N-terminal interactions of CCD86 with MRT4 and uL6 near the sarcin-ricin loop (SRL) (bottom) and domain architecture of CCD86 as compared with yeast and its interactions with other RNA and protein elements (center). Elements not visualized are colored white while visualized elements are color-coded.

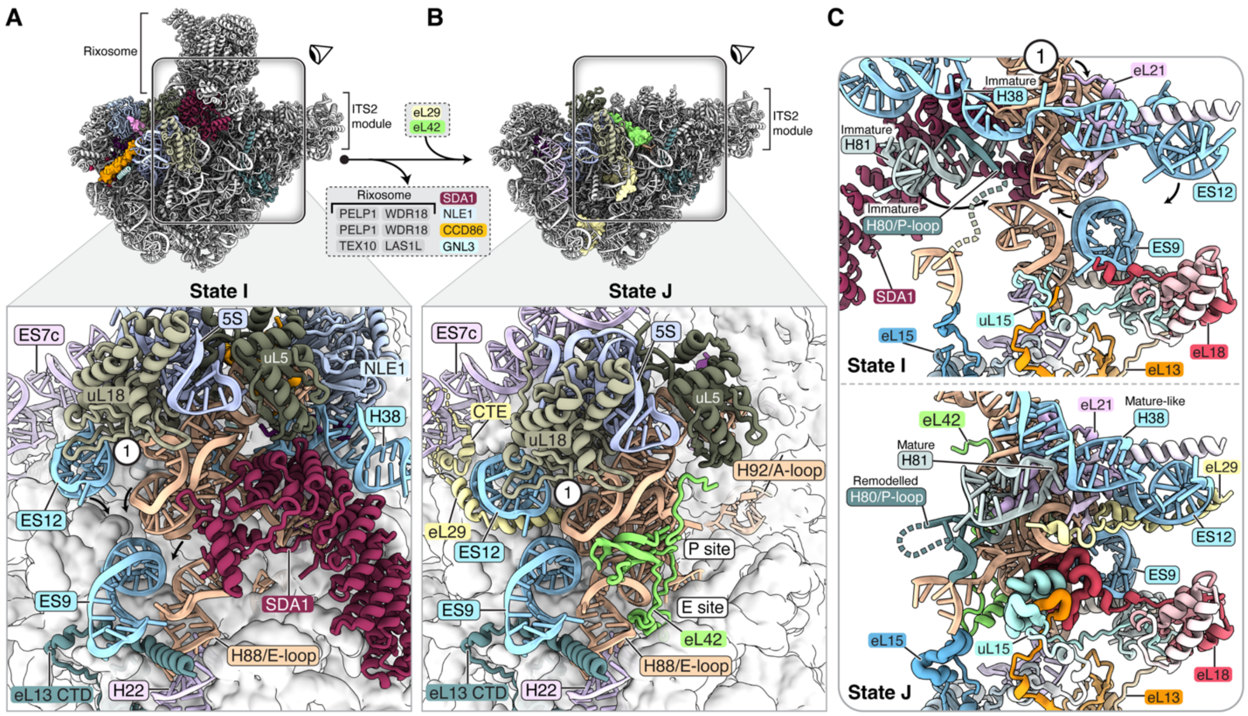


**Fig. S32. Cooperative formation of the human E-site.**

(**A**) Top view of state I with color-coded assembly factors near the immature E-site. The zoomed and rotated view below shows assembly factors and ribosomal proteins near the immature E-site. Arrows indicate movements required for E-site formation when compared to (B). (**B**) Top view of state J with color-coded ribosomal proteins near the E-site. The zoomed and rotated view below shows the mature E-site with bound eL42 (green). (**C**) Cooperative movements of pre-rRNA and ribosomal proteins required for the transition of an immature E-site in state I (top) to a mature E-site in state J (bottom). Ribosomal protein elements that only become ordered in state J1 are color-coded and shown in thick ribbon.

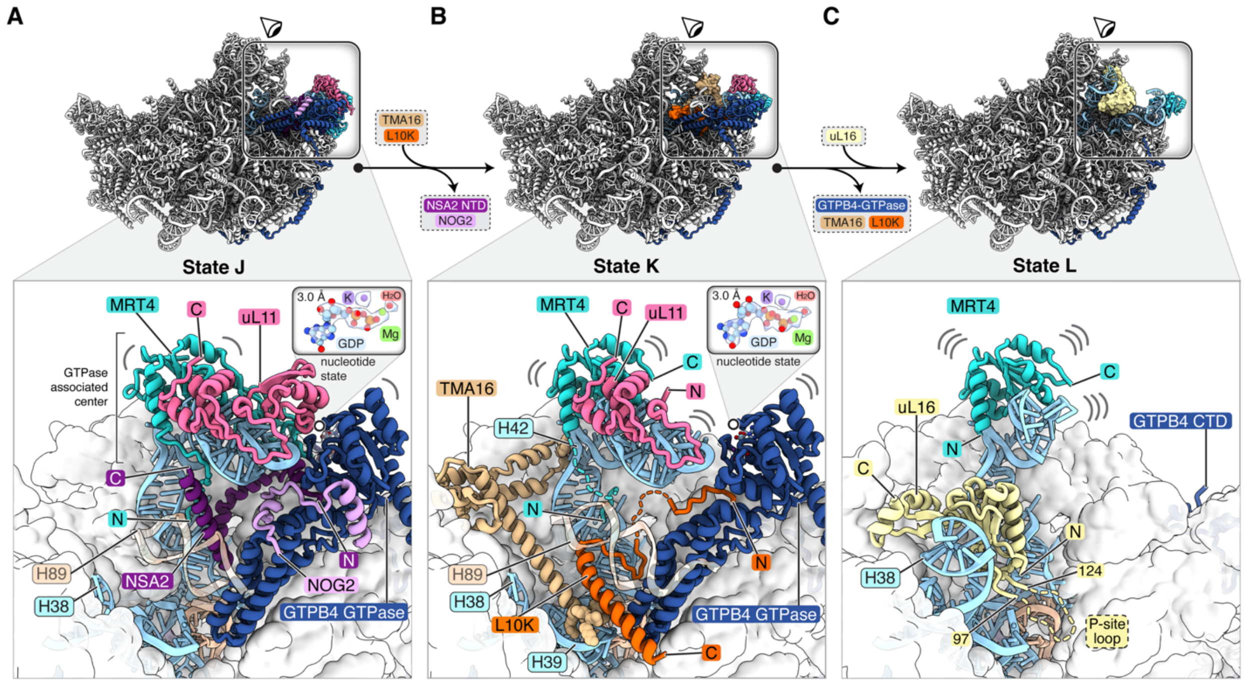


**Fig. S33. Coordinated removal of assembly factors during late nuclear maturation.**

(**A-C**) Crown views of states J (A), K (B) and L (C) with color-coded assembly factors near the GTPase associated center. Arrows with labelled assembly factors highlight transitions between the states. (A) The zoomed view below shows peptides of NSA2 and NOG2 bound to GTPB4. The inset highlights that GTPB4 is GDP-bound. (B) The zoomed view below shows L10K and TMA16 bound near GTPB4 with the inset highlighting that GTPB4 is GDP-bound. (C) The zoomed view below shows uL16 and H38 near an increasingly flexible MRT4. The disordered P-site loop of uL16 is indicated by dashes.

**Table S1. Overall and composite cryo-EM maps generated in this study.**

| Overall & composite maps | | | | | |
|---|---|---|---|---|---|
| | EMDB ID | Map | Particles no° | B-factor sharpening | Resolution (FSC-0.143) |
| State A1 | EMD-29104 | Overall | 42413 | -26.7 | 3.00 |
| | EMD-29252 | Composite | | | 2.85 |
| State A2 | EMD-29105 | Overall | 68600 | -33.5 | 2.85 |
| | EMD-29253 | Composite | | | 2.76 |
| State B1 | EMD-29106 | Overall | 21669 | -18 | 3.09 |
| | EMD-29254 | Composite | | | 2.89 |
| State B2 | EMD-29107 | Overall | 39938 | -20.2 | 3.06 |
| | EMD-29255 | Composite | | | 2.88 |
| State C1 | EMD-29108 | Overall | 45396 | -30 | 2.98 |
| | EMD-29256 | Composite | | | 2.81 |
| State C2 | EMD-29109 | Overall | 48037 | -30.8 | 3.03 |
| | EMD-29257 | Composite | | | 2.82 |
| State D1 | EMD-29110 | Overall | 209464 | -51.6 | 2.60 |
| | EMD-29258 | Composite | | | 2.47 |
| State D2 | EMD-29111 | Overall | 214795 | -52 | 2.63 |
| | EMD-29259 | Composite | | | 2.50 |
| State E | EMD-29112 | Overall | 126301 | -46 | 2.75 |
| | EMD-29260 | Composite | | | 2.59 |
| State F | EMD-29113 | Overall | 76907 | -38.7 | 2.84 |
| | EMD-29261 | Composite | | | 2.67 |
| State G | EMD-29114 | Overall | 26472 | -20.7 | 3.19 |
| | EMD-29262 | Composite | | | 3.04 |
| State H | EMD-29115 | Overall | 67272 | -43.4 | 2.96 |
| | EMD-29263 | Composite | | | 2.91 |
| State I1 | EMD-29116 | Overall | 70162 | -38.4 | 2.71 |
| | EMD-29265 | Composite | | | 2.67 |
| State I2 | EMD-29117 | Overall | 107973 | -45.5 | 2.53 |
| | EMD-29266 | Composite | | | 2.53 |
| State I3 | EMD-29118 | Overall | 22406 | -21.7 | 3.05 |
| | EMD-29267 | Composite | | | 2.89 |
| State J1 | EMD-29119 | Overall | 71912 | -38.5 | 2.76 |
| | EMD-29268 | Composite | | | 2.62 |
| State J2 | EMD-29120 | Overall | 74556 | -41.6 | 2.66 |
| | EMD-29269 | Composite | | | 2.55 |
| State J3 | EMD-29121 | Overall | 28562 | -24.7 | 2.99 |
| | EMD-29271 | Composite | | | 2.75 |
| State K1 | EMD-29122 | Overall | 64178 | -36.9 | 2.84 |
| | EMD-29272 | Composite | | | 2.63 |
| State K2 | EMD-29123 | Overall | 66157 | -42.8 | 2.64 |
| | EMD-29273 | Composite | | | 2.55 |
| State K3 | EMD-29124 | Overall | 20809 | -21.7 | 3.05 |
| | EMD-29274 | Composite | | | 2.76 |
| State L1 | **EMD-29125** | **Overall** | **88174** | **-41.9** | **2.72** |
| | EMD-29275 | Composite | | | 2.58 |
| State L2 | **EMD-29126** | **Overall** | **123749** | **-46.8** | **2.49** |
| | EMD-29276 | Composite | | | 2.48 |
| State L3 | **EMD-29127** | **Overall** | **33770** | **-29.2** | **2.83** |
| | EMD-29277 | Composite | | | 2.65 |

#### Table S2. Focused cryo-EM maps generated in this study.

| | Focused maps | | | | |
|---|---|---|---|---|---|
| | EMDB ID | Map | Particles no° | B-factor sharpening | Resolution (FSC-0.143) |
| State A1 | EMD-29130 | Domain I | 42413 | -45.5 | 2.87 |
| | EMD-29131 | Domain II | 42413 | -43.2 | 2.87 |
| State A2 | EMD-29132 | Domain I | 68600 | -50 | 2.73 |
| | EMD-29133 | Domain II | 68600 | -48.9 | 2.73 |
| States A1 & A2 | EMD-29134 | Domain VI | 111013 | -68.6 | 3.05 |
| | EMD-29135 | ITS2 | 111013 | -75.9 | 2.87 |
| State B1 | EMD-29136 | Domain I | 21669 | -35 | 2.93 |
| | EMD-29137 | Domain II | 21669 | -35.4 | 2.94 |
| State B2 | EMD-29138 | Domain I | 39938 | -39.4 | 2.87 |
| | EMD-29139 | Domain II | 39938 | -40.3 | 2.89 |
| States B1 & B2 | EMD-29140 | ITS2 | 61607 | -66.9 | 3.08 |
| | EMD-29141 | DomainVI | 61607 | -63.2 | 3.31 |
| | EMD-29142 | GTPB4-GAC | 61607 | -54.5 | 3.09 |
| States AB1 & AB2 | EMD-29128 | Root helix III | 49826 | -79.2 | 3.46 |
| | EMD-29129 | H96 | 49278 | -51.9 | 3.07 |
| State C1 | EMD-29143 | Domain I | 45396 | -46.7 | 2.83 |
| | EMD-29144 | Domain II | 45396 | -47.9 | 2.85 |
| State C2 | EMD-29145 | Domain I | 48037 | -46.8 | 2.82 |
| | EMD-29146 | Domain II | 48037 | -47.3 | 2.86 |
| States C1 & C2 | EMD-29147 | ITS2 | 93433 | -70.5 | 3.02 |
| | EMD-29148 | Domains V-VI | 93433 | -55.7 | 2.77 |
| | EMD-29149 | DDX54 | 93433 | -61.1 | 2.97 |
| | EMD-29150 | L1-stalk | 93433 | -68.4 | 3.61 |
| States A-F | EMD-29151 | uL15 | 68401 | -53.8 | 2.78 |
| State D1 | EMD-29152 | Domain I | 209464 | -56.7 | 2.48 |
| | EMD-29153 | Domain II | 209464 | -57.2 | 2.46 |
| | EMD-29154 | Domains V-VI | 209464 | -57.7 | 2.50 |
| | EMD-29155 | GNL3 | 113872 | -58 | 2.67 |
| State D2 | EMD-29156 | Domain I | 214795 | -56.9 | 2.49 |
| | EMD-29157 | Domain II | 214795 | -57.3 | 2.50 |
| | EMD-29158 | Domains V-VI | 214795 | -58.9 | 2.60 |
| States D1 & D2 | EMD-29159 | RRS1 | 424259 | -55.9 | 2.33 |
| | EMD-29160 | ITS2 | 424259 | -75.7 | 2.62 |
| | EMD-29161 | SSU interface | 281088 | -60.3 | 2.67 |
| | EMD-29162 | DDX54 | 281088 | -63.4 | 2.59 |
| | EMD-29163 | L1-stalk | 281088 | -70.7 | 2.78 |
| | EMD-29164 | NOC3L-FTSJ3 | 281088 | -74.7 | 3.19 |
| | EMD-29165 | NOC2L | 281088 | -82.3 | 3.08 |

| | Focused maps | | | | |
|---|---|---|---|---|---|
| | EMDB ID | Map | Particles no° | B-factor sharpening | Resolution (FSC-0.143) |
| State E | EMD-29166 | Domain I | 126301 | -53.3 | 2.58 |
| | EMD-29167 | Domain II | 126301 | -54.2 | 2.63 |
| | EMD-29168 | ITS2 | 126301 | -63.2 | 2.97 |
| State F | EMD-29169 | PeBoW-WDR55 | 76907 | -53.8 | 3.04 |
| | EMD-29170 | PeBoW | 76907 | -56.3 | 3.06 |
| | EMD-29171 | WDR55 | 76907 | -73.1 | 3.50 |
| | EMD-29173 | Domain III | 76907 | -49 | 2.84 |
| | EMD-29174 | ITS2 | 76907 | -59.6 | 2.91 |
| | EMD-29175 | Domain I | 76907 | -46.8 | 2.71 |
| | EMD-29176 | Domain II | 76907 | -45.3 | 2.68 |
| States E & F | EMD-29177 | SSU interface | 203208 | -60.4 | 2.86 |
| | EMD-29178 | DDX54 | 203208 | -60.2 | 2.79 |
| | EMD-29179 | L1-stalk | 203208 | -65.5 | 2.92 |
| | EMD-29180 | NOC2L-NOC3L | 203208 | -81.2 | 3.07 |
| | EMD-29181 | Domain VI | 203208 | -58.4 | 2.58 |
| | EMD-29182 | DDX18 | 203208 | -62 | 2.56 |
| | EMD-29183 | Domain V | 203208 | -57.9 | 2.58 |
| | EMD-29184 | RRS1 | 203208 | -54.7 | 2.61 |
| | EMD-29185 | NOC2L | 203208 | -99.2 | 3.75 |
| State G | EMD-29186 | ITS2 | 26472 | -38.6 | 3.35 |
| | EMD-29187 | Domain I | 26472 | -28.9 | 3.06 |
| | EMD-29188 | Domain II | 26472 | -31.4 | 3.07 |
| | EMD-29189 | Domains V-VI | 26472 | -36.6 | 3.13 |
| | EMD-29192 | Domain IV | 26472 | -31.8 | 3.10 |
| State H | EMD-29194 | Domains V-VI | 67272 | -56.2 | 2.88 |
| | EMD-29193 | 5S RNP | 67272 | -56.3 | 3.00 |
| States I1,2,3 | EMD-29195 | Central protub. | 200541 | -59.2 | 2.67 |
| | EMD-29196 | TEX10 | 200541 | -75.8 | 2.99 |
| | EMD-29197 | Rixosome | 200541 | -64.9 | 2.87 |
| States I1,J1,K1,L1 | EMD-29198 | ITS2 module | 294426 | -61.3 | 2.47 |
| States I2,J2,K2,L2 | EMD-29199 | ITS2 module | 372435 | -61.1 | 2.39 |
| States J1,2,3 | EMD-29200 | Central protub. | 175030 | -52.4 | 2.52 |
| | EMD-29201 | Domain V | 175030 | -52.8 | 2.59 |
| States K1,2,3 | EMD-29202 | Central protub. | 151144 | -52.4 | 2.51 |
| | EMD-29204 | Domain V | 151144 | -53.4 | 2.58 |
| States L1,2,3 | EMD-29205 | Central protub. | 225899 | -54.2 | 2.45 |
| | EMD-29206 | Domain V | 225899 | -54.1 | 2.52 |

#### Table S3. Model composition of the nucleolar pre-60*S* assembly intermediates.

| Subgroup | | RefSeq or Uniprot ID | Chain ID | Molecule name | Initial PDB template | State A1 | State A2 | State B1 | State B2 | State C1 | State C2 | State D1 | State D2 | State E | State F | State G | State H* |
|---|---|---|---|---|---|---|---|---|---|---|---|---|---|---|---|---|---|
| RNA | | NR_003285.3 | L1 | 5.8S | 6LSS | | | | | | | | | | | | |
| | | NR_046235.3 | L2 | ITS2 | manual | | | | | | | | | | | | |
| | | NR_003287.4 | L3 | 28S | 6LSS | | | | | | | | | | | | |
| | | NR_023369.1 | L4 | 5S | 6LSS | | | | | | | | | | | | |
| Ribosomal proteins | | P62906 | BB | RL10A_uL1 | AlphaFold | | | | | | | | | | | | |
| | | P62913 | L5 | RL11_uL5 | 6LSS | | | | | | | | | | | | |
| | | P30050 | BA | RL12_uL11 | AlphaFold | | | | | | | | | | | | |
| | | P26373 | L6 | RL13_eL13 | 6LSS | | | | | | | | | | | | |
| | | P40429 | L7 | RL13A_uL13 | 6LSS | | | | | | | | | | | | |
| | | P50914 | L8 | RL14_eL14 | 6LSS | | | | | | | | | | | | |
| | | P61313 | L9 | RL15_eL15 | 6LSS | | | | | | | | | | | | |
| | | P18621 | LA | RL17_uL22 | 6LSS | | | | | | | | | | | | |
| | | Q07020 | LB | RL18_eL18 | 6LSS | | | | | | | | | | | | |
| | | Q02543 | LC | RL18A_eL20 | 6LSS | | | | | | | | | | | | |
| | | P84098 | LD | RL19_eL19 | 6LSS | | | | | | | | | | | | |
| | | P46778 | LE | RL21_eL21 | 6LSS | | | | | | | | | | | | |
| | | P35268 | LF | RL22_eL22 | 6LSS | | | | | | | | | | | | |
| | | P62829 | LG | RL23_uL14 | 6LSS | | | | | | | | | | | | |
| | | P62750 | LH | RL23A_uL23 | 6LSS | | | | | | | | | | | | |
| | | P61254 | LI | RL26_uL24 | 6LSS | | | | | | | | | | | | |
| | | P61353 | LJ | RL27_eL27 | 6LSS | | | | | | | | | | | | |
| | | P46776 | LK | RL27A_uL15 | 6LSS | | | | | | | | | | | | |
| | | P46779 | LL | RL28_eL28 | 6LSS | | | | | | | | | | | | |
| | | P39023 | LN | RL3_uL3 | 6LSS | | | | | | | | | | | | |
| | | P62888 | LO | RL30_eL30 | 6LSS | | | | | | | | | | | | |
| | | P62899 | LP | RL31_eL31 | 6LSS | | | | | | | | | | | | |
| | | P62910 | LQ | RL32_eL32 | 6LSS | | | | | | | | | | | | |
| | | P49207 | LR | RL34_eL34 | 6LSS | | | | | | | | | | | | |
| | | P42766 | LS | RL35_uL29 | 6LSS | | | | | | | | | | | | |
| | | P18077 | LT | RL35A_eL33 | 6LSS | | | | | | | | | | | | |
| | | Q9Y3U8 | LU | RL36_eL36 | 6LSS | | | | | | | | | | | | |
| | | P61927 | LW | RL37_eL37 | 6LSS | | | | | | | | | | | | |
| | | P61513 | LX | RL37A_eL43 | 6LSS | | | | | | | | | | | | |
| | | P63173 | LY | RL38_eL38 | 6LSS | | | | | | | | | | | | |
| | | P62891 | LZ | RL39_eL39 | 6LSS | | | | | | | | | | | | |
| | | P36578 | SA | RL4_uL4 | 6LSS | | | | | | | | | | | | |
| | | P46777 | SB | RL5_uL18 | 6LSS | | | | | | | | | | | | |
| | | Q02878 | SC | RL6_eL6 | 6LSS | | | | | | | | | | | | |
| | | P18124 | SD | RL7_uL30 | 6LSS | | | | | | | | | | | | |
| | | P62424 | SE | RL7A_eL8 | 6LSS | | | | | | | | | | | | |
| | | P62917 | SF | RL8_uL2 | 6LSS | | | | | | | | | | | | |
| | | P32969 | SG | RL9_uL6 | 6LSS | | | | | | | | | | | | |
| Assembly factors | ITS2 module | Q9BYG3 | SH | MK67I | AlphaFold | | | | | | | | | | | | |
| | | Q6DKI1 | SI | RLP7 | AlphaFold | | | | | | | | | | | | |
| | | O76021 | SL | RL1D1 | AlphaFold | | | | | | | | | | | | |
| | | O00541 | SM | PES1 | AlphaFold | | | | | | | | | | | | |
| | | Q9NZM5 | NL | NOP53 | AlphaFold | | | | | | | | | | | | |
| | Ubiquitous | P56537 | SK | eIF6 | AlphaFold | | | | | | | | | | | | |
| | | Q9UHA3 | SV | RLP24 | AlphaFold | | | | | | | | | | | | |
| | Early nucleolar | Q99848 | SN | EBP2 | AlphaFold | | | | | | | | | | | | |
| | | Q8TDN6 | SO | BRX1 | AlphaFold | | | | | | | | | | | | |
| | | Q8IY81 | SJ | FTSJ3 | AlphaFold | | | | | | | | | | | | |
| | | Q14137 | SS | BOP1 | AlphaFold | | | | | | | | | | | | |
| | | Q9NVP1 | SW | DDX18 | AlphaFold | | | | | | | | | | | | |
| | | Q9Y3C1 | SZ | NOP16 | AlphaFold | | | | | | | | | | | | |
| | | Q9BVP2 | NB | GNL3 | AlphaFold | | | | | | | | | | | | |
| | Solvent side | Q9BXY0 | NM | MAK16 | AlphaFold | | | | | | | | | | | | |
| | | P56182 | NO | RRP1 | AlphaFold | | | | | | | | | | | | |
| | | Q6RFH5 | NQ | WDR74 | AlphaFold | | | | | | | | | | | | |
| | | Q9H9Y2 | NS | RPF1 | AlphaFold | | | | | | | | | | | | |
| | Domain VI | Q9NQ55 | NN | SSF1 | AlphaFold | | | | | | | | | | | | |
| | | O75683 | NE | SURF6 | AlphaFold | | | | | | | | | | | | |
| | | Q9Y3B9 | NG | RRP15 | AlphaFold | | | | | | | | | | | | |
| | PTC/GAC | Q9BZE4 | SR | GTPB4 | AlphaFold | | | | | | | | | | | | |
| | | Q9UKD2 | SQ | MRT4 | AlphaFold | | | | | | | | | | | | |
| | | O95478 | NF | NSA2 | AlphaFold | | | | | | | | | | | | |
| | Late nucleolar | Q8WTT2 | SU | NOC3L | AlphaFold | | | | | | | | | | | | |
| | | Q9Y3T9 | NA | NOC2L | AlphaFold | | | | | | | | | | | | |
| | | P46087 | SY | NOP2 | AlphaFold | | | | | | | | | | | | |
| | | Q9Y221 | NH | NIP7 | AlphaFold | | | | | | | | | | | | |
| | | Q13823 | NC | NOG2 | AlphaFold | | | | | | | | | | | | |
| | | Q8TDD1 | NI | DDX54 | AlphaFold | | | | | | | | | | | | |
| | | Q9BRT6 | NK | LLPH | AlphaFold | | | | | | | | | | | | |
| | | Q9NVX2 | NJ | NLE1 | AlphaFold | | | | | | | | | | | | |
| | | Q9H6F5 | NZ | CCD86 | AlphaFold | | | | | | | | | | | | |
| | Domain III | Q9GZL7 | NP | WDR12 | AlphaFold | | | | | | | | | | | | |
| | | Q9H6Y2 | BC | WDR55 | 7KQQ | | | | | | | | | | | | |
| | 5S RNP | Q9H7B2 | ND | RPF2 | AlphaFold | | | | | | | | | | | | |
| | | Q15050 | ST | RRS1 | AlphaFold | | | | | | | | | | | | |

*As state H is a partial EM reconstruction, the model composition is not complete.

### Table S4. Model composition the nuclear pre-60*S* assembly intermediates.

| Subgroup | | RefSeq or Uniprot ID | Chain ID | Molecule name | Initial PDB template | State I1 | State I2 | State I3 | State J1 | State J2 | State J3 | State K1 | State K2 | State K3 | State L1 | State L2 | State L3 |
|---|---|---|---|---|---|---|---|---|---|---|---|---|---|---|---|---|---|
| RNA | | NR_003285.3 | L1 | 5.8S | 6LSS | | | | | | | | | | | | |
| | | NR_046235.3 | L2 | ITS2 | manual | | | | | | | | | | | | |
| | | NR_003287.4 | L3 | 28S | 6LSS | | | | | | | | | | | | |
| | | NR_023369.1 | L4 | 5S | 6LSS | | | | | | | | | | | | |
| Ribosomal proteins | | P27635 | BE | RL10_uL16 | AlphaFold | | | | | | | | | | | | |
| | | P62906 | BB | RL10A_uL1 | AlphaFold | | | | | | | | | | | | |
| | | P62913 | L5 | RL11_uL5 | 6LSS | | | | | | | | | | | | |
| | | P30050 | BA | RL12_uL11 | AlphaFold | | | | | | | | | | | | |
| | | P26373 | L6 | RL13_eL13 | 6LSS | | | | | | | | | | | | |
| | | P40429 | L7 | RL13A_uL13 | 6LSS | | | | | | | | | | | | |
| | | P50914 | L8 | RL14_eL14 | 6LSS | | | | | | | | | | | | |
| | | P61313 | L9 | RL15_eL15 | 6LSS | | | | | | | | | | | | |
| | | P18621 | LA | RL17_uL22 | 6LSS | | | | | | | | | | | | |
| | | Q07020 | LB | RL18_eL18 | 6LSS | | | | | | | | | | | | |
| | | Q02543 | LC | RL18A_eL20 | 6LSS | | | | | | | | | | | | |
| | | P84098 | LD | RL19_eL19 | 6LSS | | | | | | | | | | | | |
| | | P46778 | LE | RL21_eL21 | 6LSS | | | | | | | | | | | | |
| | | P35268 | LF | RL22_eL22 | 6LSS | | | | | | | | | | | | |
| | | P62829 | LG | RL23_uL14 | 6LSS | | | | | | | | | | | | |
| | | P62750 | LH | RL23A_uL23 | 6LSS | | | | | | | | | | | | |
| | | P61254 | LI | RL26_uL24 | 6LSS | | | | | | | | | | | | |
| | | P61353 | LJ | RL27_eL27 | 6LSS | | | | | | | | | | | | |
| | | P46776 | LK | RL27A_uL15 | 6LSS | | | | | | | | | | | | |
| | | P46779 | LL | RL28_eL28 | 6LSS | | | | | | | | | | | | |
| | | P47914 | LM | RL29_eL29 | 6LSS | | | | | | | | | | | | |
| | | P39023 | LN | RL3_uL3 | 6LSS | | | | | | | | | | | | |
| | | P62888 | LO | RL30_eL30 | 6LSS | | | | | | | | | | | | |
| | | P62899 | LP | RL31_eL31 | 6LSS | | | | | | | | | | | | |
| | | P62910 | LQ | RL32_eL32 | 6LSS | | | | | | | | | | | | |
| | | P49207 | LR | RL34_eL34 | 6LSS | | | | | | | | | | | | |
| | | P42766 | LS | RL35_uL29 | 6LSS | | | | | | | | | | | | |
| | | P18077 | LT | RL35A_eL33 | 6LSS | | | | | | | | | | | | |
| | | Q9Y3U8 | LU | RL36_eL36 | 6LSS | | | | | | | | | | | | |
| | | P83881 | LV | RL36A_eL42 | 6LSS | | | | | | | | | | | | |
| | | P61927 | LW | RL37_eL37 | 6LSS | | | | | | | | | | | | |
| | | P61513 | LX | RL37A_eL43 | 6LSS | | | | | | | | | | | | |
| | | P63173 | LY | RL38_eL38 | 6LSS | | | | | | | | | | | | |
| | | P62891 | LZ | RL39_eL39 | 6LSS | | | | | | | | | | | | |
| | | P36578 | SA | RL4_uL4 | 6LSS | | | | | | | | | | | | |
| | | P46777 | SB | RL5_uL18 | 6LSS | | | | | | | | | | | | |
| | | Q02878 | SC | RL6_eL6 | 6LSS | | | | | | | | | | | | |
| | | P18124 | SD | RL7_uL30 | 6LSS | | | | | | | | | | | | |
| | | P62424 | SE | RL7A_eL8 | 6LSS | | | | | | | | | | | | |
| | | P62917 | SF | RL8_uL2 | 6LSS | | | | | | | | | | | | |
| | | P32969 | SG | RL9_uL6 | 6LSS | | | | | | | | | | | | |
| Assembly factors | ITS2 module | Q9BYG3 | SH | MK67I | AlphaFold | | | | | | | | | | | | |
| | | Q6DKI1 | SI | RLP7 | AlphaFold | | | | | | | | | | | | |
| | | O76021 | SL | RL1D1 | AlphaFold | | | | | | | | | | | | |
| | | O00541 | SM | PES1 | AlphaFold | | | | | | | | | | | | |
| | | Q9NZM5 | NL | NOP53 | AlphaFold | | | | | | | | | | | | |
| | Ubiquitous | P56537 | SK | eIF6 | AlphaFold | | | | | | | | | | | | |
| | | Q9UHA3 | SV | RLP24 | AlphaFold | | | | | | | | | | | | |
| | Nucleolar | Q9BVP2 | NB | GNL3 | AlphaFold | | | | | | | | | | | | |
| | | Q9BRT6 | NK | LLPH | AlphaFold | | | | | | | | | | | | |
| | | Q13823 | NC | NOG2 | AlphaFold | | | | | | | | | | | | |
| | | Q9BZE4 | SR | GTPB4 | AlphaFold | | | | | | | | | | | | |
| | | Q9UKD2 | SQ | MRT4 | AlphaFold | | | | | | | | | | | | |
| | | O95478 | NF | NSA2 | AlphaFold | | | | | | | | | | | | |
| | Nuclear | Q9NVX2 | NJ | NLE1 | AlphaFold | | | | | | | | | | | | |
| | | O00488 | NP | ZN593 | AlphaFold | | | | | | | | | | | | |
| | | Q96EY4 | NR | TMA16 | AlphaFold | | | | | | | | | | | | |
| | | Q9H6F5 | NZ | CCD86 | AlphaFold | | | | | | | | | | | | |
| | | Q9UNZ5 | VB | L10K/C19orf53 | Manual | | | | | | | | | | | | |
| | Rixosome | Q9NVU7 | NT | SDA1 | AlphaFold | | | | | | | | | | | | |
| | | Q9NXF1 | NU | TEX10 | AlphaFold | | | | | | | | | | | | |
| | | Q9BV38 | NV | WDR18.1 | AlphaFold | | | | | | | | | | | | |
| | | Q9BV38 | NW | WDR18.2 | AlphaFold | | | | | | | | | | | | |
| | | Q8IZL8 | NX | PELP1.1 | AlphaFold | | | | | | | | | | | | |
| | | Q8IZL8 | NY | PELP1.2 | AlphaFold | | | | | | | | | | | | |
| | | Q9Y4W2 | BD | LAS1L | AlphaFold | | | | | | | | | | | | |

#### Table S5. Cryo-EM data collection and refinement statistics.

| | **State A1** PDB 8FKP EMD-29252 | **State A2** PDB 8FKQ EMD-29253 | **State B1** PDB 8FKR EMD-29254 | **State B2** PDB 8FKS EMD-29255 | **State C1** PDB 8FKT EMD-29256 | **State C2** PDB 8FKU EMD-29257 |
|---|---|---|---|---|---|---|
| **Data collection and processing** | | | | | | |
| Microscope | Titan Krios | Titan Krios | Titan Krios | Titan Krios | Titan Krios | Titan Krios |
| Voltage (keV) | 300 | 300 | 300 | 300 | 300 | 300 |
| Camera | K3 | K3 | K3 | K3 | K3 | K3 |
| Magnification | 64,000 | 64,000 | 64,000 | 64,000 | 64,000 | 64,000 |
| Pixel size at detector (Å/pixel) | 1.072 | 1.072 | 1.072 | 1.072 | 1.072 | 1.072 |
| Total electron exposure (e-/Å2) | 60 | 60 | 60 | 60 | 60 | 60 |
| Exposure rate (e-/pixel/sec) | 30 | 30 | 30 | 30 | 30 | 30 |
| Number of frames collected (no.) | 40 | 40 | 40 | 40 | 40 | 40 |
| Defocus range (μm) | -0.5 to -2.5 | -0.5 to -2.5 | -0.5 to -2.5 | -0.5 to -2.5 | -0.5 to -2.5 | -0.5 to -2.5 |
| Energy filter slit width (V) | 20 | 20 | 20 | 20 | 20 | 20 |
| Automation software | SerialEM | SerialEM | SerialEM | SerialEM | SerialEM | SerialEM |
| Micrographs collected (no.) | 172,699 | 172,699 | 172,699 | 172,699 | 172,699 | 172,699 |
| Total extracted particles (no.) | 15,679,142 | 15,679,142 | 15,679,142 | 15,679,142 | 15,679,142 | 15,679,142 |
| Final particle images (no.) | 42413 | 68,600 | 21,669 | 39,938 | 45,396 | 48,037 |
| Point-group symmetry | C1 | C1 | C1 | C1 | C1 | C1 |
| Resolution (global at FSC-0.143, Å) | | | | | | |
| Overall map (masked) | 3.00 | 2.85 | 3.09 | 3.06 | 2.98 | 2.82 |
| Composite map (masked)* | 2.85 | 2.76 | 2.89 | 2.88 | 2.81 | 3.03 |
| Map sharpening *B* factor ($Å^2$) | | | | | | |
| Overall map | -26.7 | -33.5 | -18 | -20.2 | -30 | -30.8 |
| Focused maps | -43.2 to -79.2 | -48.9 to -79.2 | -35.0 to -79.2 | -39.4 to -79.2 | -45.3 to -70.5 | -46.8 to -70.5 |
| **Model composition** | | | | | | |
| Non-hydrogen atoms | 101,451 | 93,805 | 116,287 | 108,693 | 134,916 | 125,840 |
| Chains (proteins/RNA) | 44/3 | 40/3 | 48/3 | 44/3 | 50/3 | 47/3 |
| Protein residues (non-modified/modified) | 7959/4 | 6879/3 | 9087/4 | 8023/3 | 10,794/3 | 9433/2 |
| RNA (non-modified/modified) | 1728/0 | 1784/0 | 1990/0 | 2045/0 | 2346/0 | 2406/0 |
| Ligands ($Mg^{2+}$/$Zn^{2+}$/ $K^{+}$/$Fe_4$-$S_4$/ GDP/ADP/GTP) | 54/2/0/1/ 0/0/0 | 51/2/0/0/ 0/0/0 | 55/2/0/1/ 1/0/0 | 62/2/0/0/ 1/0/0 | 64/2/0/1/ 1/0/0 | 58/2/0/0/ 1/0/0 |
| **Model refinement** | | | | | | |
| Refinement package (real space) | Phenix 1.19.1 | Phenix 1.19.1 | Phenix 1.19.1 | Phenix 1.19.1 | Phenix 1.19.1 | Phenix 1.19.1 |
| Initial models used (PDB code) | 6LSS | 6LSS | 6LSS | 6LSS | 6LSS | 6LSS |
| Model resolution cutoff (Å) | 3.4 | 3.3 | 3.5 | 3.5 | 3.4 | 3.4 |
| Model-Map CC ($CC_{mask}$/$CC_{box}$/ $CC_{peaks}$/$CC_{volume}$) | 0.83/0.76/ 0.74/0.82 | 0.82/0.77/ 0.75/0.82 | 0.82/0.75/ 0.73/0.81 | 0.81/0.75/ 0.72/0.80 | 0.79/0.73/ 0.70/0.79 | 0.82/0.75/ 0.73/0.81 |
| Model-to-map FSC, threshold 0.50 (masked/unmasked, Å) | 3.06/3.06 | 3.00/3.00 | 3.20/3.20 | 3.14/3.14 | 3.00/3.00 | 3.00/3.00 |
| **RMSDs** | | | | | | |
| Bond lengths (Å) | 0.005 | 0.004 | 0.005 | 0.003 | 0.005 | 0.006 |
| Bond angles (°) | 0.842 | 0.781 | 0.840 | 0.776 | 0.829 | 0.863 |
| **B-factors** | | | | | | |
| Average B-factors ($A^2$) | | | | | | |
| Protein | 46.47 | 36.72 | 49.33 | 46.71 | 34.21 | 36.19 |
| RNA | 68.83 | 57.33 | 67.82 | 70.40 | 45.69 | 55.03 |
| Ligand | 24.41 | 20.36 | 27.82 | 28.09 | 24.54 | 20.75 |
| **Validation** | | | | | | |
| Clash score | 2.11 | 1.93 | 2.05 | 2.19 | 1.95 | 2.05 |
| MolProbity score | 0.98 | 0.96 | 0.98 | 0.99 | 0.96 | 0.98 |
| CaBLAM outliers | 0.72 | 0.80 | 0.75 | 0.74 | 0.67 | 0.80 |
| Poor rotamers (%) | 0.00 | 0.02 | 0.04 | 0.01 | 0.02 | 0.01 |
| C-beta deviations | 0.00 | 0.00 | 0.00 | 0.00 | 0.00 | 0.00 |
| EMRinger score | 3.45 | 3.34 | 3.31 | 3.17 | 3.68 | 3.77 |
| **Ramachandran plot** | | | | | | |
| Favored (%) | 98.83 | 98.86 | 98.56 | 99.06 | 98.75 | 98.36 |
| Allowed (%) | 1.17 | 1.14 | 1.44 | 0.94 | 1.25 | 1.64 |
| Outliers (%) | 0.00 | 0.00 | 0.00 | 0.00 | 0.00 | 0.00 |
| **RNA validation** | | | | | | |
| Average suiteness (%) | 57.2 | 60.0 | 54.0 | 58.2 | 56.9 | 55.1 |
| Good sugar puckers (%) | 99.88 | 99.66 | 99.89 | 99.95 | 99.82 | 99.75 |

*Fourier shell correlation calculated between composite half maps.

| | State D1<br>PDB 8FKV<br>EMD-29258 | State D2<br>PDB 8FKW<br>EMD-29259 | State E<br>PDB 8FKX<br>EMD-29260 | State F<br>PDB 8FKY<br>EMD-29261 | State G<br>PDB 8FKZ<br>EMD-29262 | State H<br>PDB 8FL0<br>EMD-29263 |
|---|---|---|---|---|---|---|
| **Data collection and processing** | | | | | | |
| Microscope | Titan Krios | Titan Krios | Titan Krios | Titan Krios | Titan Krios | Titan Krios |
| Voltage (keV) | 300 | 300 | 300 | 300 | 300 | 300 |
| Camera | K3 | K3 | K3 | K3 | K3 | K3 |
| Magnification | 64,000 | 64,000 | 64,000 | 64,000 | 64,000 | 64,000 |
| Pixel size at detector (Å/pixel) | 1.072 | 1.072 | 1.072 | 1.072 | 1.072 | 1.072 |
| Total electron exposure (e-/Å2) | 60 | 60 | 60 | 60 | 60 | 60 |
| Exposure rate (e-/pixel/sec) | 30 | 30 | 30 | 30 | 30 | 30 |
| Number of frames collected (no.) | 40 | 40 | 40 | 40 | 40 | 40 |
| Defocus range (µm) | -0.5 to -2.5 | -0.5 to -2.5 | -0.5 to -2.5 | -0.5 to -2.5 | -0.5 to -2.5 | -0.5 to -2.5 |
| Energy filter slit width (V) | 20 | 20 | 20 | 20 | 20 | 20 |
| Automation software | SerialEM | SerialEM | SerialEM | SerialEM | SerialEM | SerialEM |
| Micrographs collected (no.) | 172,699 | 172,699 | 172,699 | 172,699 | 172,699 | 172,699 |
| Total extracted particles (no.) | 15,679,142 | 15,679,142 | 15,679,142 | 15,679,142 | 15,679,142 | 15,679,142 |
| Final particle images (no.) | 209,464 | 214,795 | 126,301 | 76,907 | 26,472 | 67,272 |
| Point-group symmetry | C1 | C1 | C1 | C1 | C1 | C1 |
| Resolution (global at FSC-0.143, Å) | | | | | | |
| Overall map (masked) | 2.60 | 2.63 | 2.75 | 2.84 | 3.19 | 2.96 |
| Composite map (masked)* | 2.47 | 2.50 | 2.59 | 2.67 | 3.04 | 2.91 |
| Map sharpening *B* factor (Å$^2$) | | | | | | |
| Overall map | -51.6 | -52.0 | -46.0 | -38.7 | -20.7 | -43.4 |
| Focused maps | -55.9 to -82.3 | -55.9 to -82.3 | -53.3 to -99.2 | -45.3 to -99.2 | -28.9 to -38.6 | -56.2 to -56.7 |
| **Model composition** | | | | | | |
| Non-hydrogen atoms | 148,974 | 139,782 | 138,642 | 158,283 | 129,997 | 83,376 |
| Chains (proteins/RNA) | 53/3 | 50/3 | 50/3 | 56/3 | 48/3 | 29/2 |
| Protein residues (non-modified/modified) | 12,244/4 | 10,862/2 | 10,791/2 | 12,400/2 | 8097/0 | 5867/0 |
| RNA (non-modified/modified) | 2434/0 | 2498/0 | 2471/0 | 2809/0 | 3055/0 | 1678/0 |
| Ligands ($Mg^{2+}$/$Zn^{2+}$/ $K^+$/$Fe_4$-$S_4$/ GDP/ADP/GTP) | 65/2/0/1/ 1/1/0 | 59/2/0/0/ 1/1/0 | 61/2/0/0/ 1/1/0 | 70/3/0/0/ 1/1/0 | 64/4/0/0/ 1/0/0 | 22/1/1/0/ 1/0/1 |
| **Model refinement** | | | | | | |
| Refinement package (real space) | Phenix 1.19.1 | Phenix 1.19.1 | Phenix 1.19.1 | Phenix 1.19.1 | Phenix 1.19.1 | Phenix 1.19.1 |
| Initial models used (PDB code) | 6LSS | 6LSS | 6LSS | 6LSS/7KQQ | 6LSS | 6LSS |
| Model resolution cutoff (Å) | 3.0 | 3.0 | 3.2 | 3.2 | 3.6 | 3.2 |
| Model-Map CC ($CC_{mask}$/$CC_{box}$/ $CC_{peaks}$/$CC_{volume}$) | 0.84/0.81/ 0.79/0.84 | 0.82/0.80/ 0.78/0.82 | 0.82/0.78/ 0.77/0.82 | 0.79/0.77/ 0.75/0.79 | 0.83/0.75/ 0.70/0.82 | 0.84/0.79/ 0.73/0.82 |
| Model-to-map FSC, threshold 0.50 (masked/unmasked, Å) | 2.67/2.67 | 2.80/2.80 | 2.86/2.86 | 2.96/2.96 | 3.15/3.15 | 3.03/3.03 |
| **RMSDs** | | | | | | |
| Bond lengths (Å) | 0.004 | 0.004 | 0.004 | 0.003 | 0.003 | 0.003 |
| Bond angles (°) | 0.803 | 0.782 | 0.793 | 0.777 | 0.740 | 0.747 |
| **B-factors** | | | | | | |
| Average B-factors (A$^2$) | | | | | | |
| Protein | 39.36 | 36.30 | 38.02 | 39.14 | 67.89 | 52.62 |
| RNA | 49.50 | 50.04 | 56.15 | 54.14 | 78.66 | 85.2 |
| Ligand | 24.83 | 28.08 | 34.91 | 34.83 | 42.68 | 37.71 |
| **Validation** | | | | | | |
| Clash score | 1.85 | 1.60 | 1.79 | 1.74 | 2.21 | 1.96 |
| MolProbity score | 0.95 | 0.91 | 0.94 | 0.93 | 1.00 | 0.96 |
| CaBLAM outliers | 0.60 | 0.66 | 0.69 | 0.71 | 0.82 | 0.90 |
| Poor rotamers (%) | 0.00 | 0.00 | 0.02 | 0.01 | 0.03 | 0.02 |
| C-beta deviations | 0.00 | 0.00 | 0.00 | 0.00 | 0.00 | 0.00 |
| EMRinger score | 3.66 | 3.73 | 3.55 | 3.34 | 2.98 | 3.17 |
| **Ramachandran plot** | | | | | | |
| Favored (%) | 98.94 | 98.91 | 98.75 | 98.99 | 99.02 | 98.89 |
| Allowed (%) | 1.06 | 1.09 | 1.25 | 1.01 | 0.98 | 1.11 |
| Outliers (%) | 0.00 | 0.00 | 0.00 | 0.00 | 0.00 | 0.00 |
| **RNA validation** | | | | | | |
| Average suiteness (%) | 58.8 | 59.3 | 59.0 | 59.6 | 59.1 | 61.1 |
| Good sugar puckers (%) | 99.67 | 99.76 | 99.75 | 99.64 | 99.70 | 99.52 |

*Fourier shell correlation calculated between composite half maps.

| | **State I1**<br>PDB 8FL2<br>EMD-29265 | **State I2**<br>PDB 8FL3<br>EMD-29266 | **State I3**<br>PDB 8FL4<br>EMD-29267 | **State J1**<br>PDB 8FL6<br>EMD-29268 | **State J2**<br>PDB 8FL7<br>EMD-29269 | **State J3**<br>PDB 8FL9<br>EMD-29271 |
|---|---|---|---|---|---|---|
| **Data collection and processing** | | | | | | |
| Microscope | Titan Krios | Titan Krios | Titan Krios | Titan Krios | Titan Krios | Titan Krios |
| Voltage (keV) | 300 | 300 | 300 | 300 | 300 | 300 |
| Camera | K3 | K3 | K3 | K3 | K3 | K3 |
| Magnification | 64,000 | 64,000 | 64,000 | 64,000 | 64,000 | 64,000 |
| Pixel size at detector (Å/pixel) | 1.072 | 1.072 | 1.072 | 1.072 | 1.072 | 1.072 |
| Total electron exposure (e-/Å2) | 60 | 60 | 60 | 60 | 60 | 60 |
| Exposure rate (e-/pixel/sec) | 30 | 30 | 30 | 30 | 30 | 30 |
| Number of frames collected (no.) | 40 | 40 | 40 | 40 | 40 | 40 |
| Defocus range (μm) | -0.5 to -2.5 | -0.5 to -2.5 | -0.5 to -2.5 | -0.5 to -2.5 | -0.5 to -2.5 | -0.5 to -2.5 |
| Energy filter slit width (V) | 20 | 20 | 20 | 20 | 20 | 20 |
| Automation software | SerialEM | SerialEM | SerialEM | SerialEM | SerialEM | SerialEM |
| Micrographs collected (no.) | 172,699 | 172,699 | 172,699 | 172,699 | 172,699 | 172,699 |
| Total extracted particles (no.) | 15,679,142 | 15,679,142 | 15,679,142 | 15,679,142 | 15,679,142 | 15,679,142 |
| Final particle images (no.) | 70,162 | 107,973 | 22,406 | 71,912 | 74,556 | 28,562 |
| Point-group symmetry | C1 | C1 | C1 | C1 | C1 | C1 |
| Resolution (global at FSC-0.143, Å) | | | | | | |
| Overall map (masked) | 2.71 | 2.53 | 3.05 | 2.76 | 2.66 | 2.99 |
| Composite map (masked)* | 2.67 | 2.53 | 2.89 | 2.62 | 2.55 | 2.75 |
| Map sharpening *B* factor (Å$^2$) | | | | | | |
| Overall map | -38.4 | -45.5 | -21.7 | -38.5 | -41.6 | -24.7 |
| Focused maps | -59.2 to -75.8 | 59.2 to -75.8 | 59.2 to -75.8 | -52.4 to -61.3 | -52.4 to -61.1 | -52.4 to -52.8 |
| **Model composition** | | | | | | |
| Non-hydrogen atoms | 189,620 | 185,200 | 180,499 | 153,983 | 149,783 | 140,156 |
| Chains (proteins/RNA) | 61/4 | 60/3 | 59/3 | 52/4 | 51/3 | 47/3 |
| Protein residues (non-modified/modified) | 13,356/1 | 13,005/1 | 12,628/1 | 9065/1 | 8710/1 | 7689/1 |
| RNA (non-modified/modified) | 3838/131 | 3634/131 | 3626/131 | 3657/107 | 3589/107 | 3541/107 |
| Ligands ($Mg^{2+}$/$Zn^{2+}$/ $K^+$/GDP/GTP) | 100/5/1/1/1 | 100/5/1/1/1 | 100/5/1/1/1 | 88/6/0/1/0 | 88/6/0/1/0 | 88/6/0/1/0 |
| **Model refinement** | | | | | | |
| Refinement package (real space) | Phenix 1.19.1 | Phenix 1.19.1 | Phenix 1.19.1 | Phenix 1.19.1 | Phenix 1.19.1 | Phenix 1.19.1 |
| Initial models used (PDB code) | 6LSS | 6LSS | 6LSS | 6LSS | 6LSS | 6LSS |
| Model resolution cutoff (Å) | 3.0 | 2.8 | 3.4 | 3.1 | 3.0 | 3.3 |
| Model-Map CC ($CC_{mask}$/$CC_{box}$/ $CC_{peaks}$/$CC_{volume}$) | 0.85/0.84/ 0.79/0.84 | 0.85/0.85/ 0.79/0.85 | 0.81/081/ 0.73/0.81 | 0.85/0.85/ 0.79/0.84 | 0.87/0.86 0.82/0.86 | 0.83/0.83/ 0.76/0.82 |
| Model-to-map FSC, threshold 0.50 (masked/unmasked, Å) | 2.88/2.88 | 2.73/2.73 | 3.06/3.06 | 2.87/2.87 | 2.78/2.78 | 2.99/2.99 |
| **RMSDs** | | | | | | |
| Bond lengths (Å) | 0.005 | 0.004 | 0.003 | 0.003 | 0.004 | 0.006 |
| Bond angles (°) | 0.809 | 0.775 | 0.779 | 0.741 | 0.760 | 0.798 |
| **B-factors** | | | | | | |
| Average B-factors (A$^2$) | | | | | | |
| Protein | 57.52 | 62.64 | 61.42 | 44.55 | 40.72 | 57.19 |
| RNA | 68.74 | 73.01 | 75.34 | 55.99 | 51.73 | 73.23 |
| Ligand | 41.83 | 51.02 | 45.05 | 28.99 | 26.16 | 36.73 |
| **Validation** | | | | | | |
| Clash score | 2.59 | 2.26 | 2.76 | 2.34 | 2.19 | 2.66 |
| MolProbity score | 1.05 | 1.00 | 1.06 | 1.01 | 0.99 | 1.10 |
| CaBLAM outliers | 0.84 | 0.79 | 0.80 | 1.14 | 1.06 | 1.33 |
| Poor rotamers (%) | 0.00 | 0.01 | 0.00 | 0.00 | 0.01 | 0.03 |
| C-beta deviations | 0.00 | 0.00 | 0.00 | 0.00 | 0.00 | 0.00 |
| EMRinger score | 3.74 | 3.84 | 3.21 | 3.86 | 4.11 | 3.70 |
| **Ramachandran plot** | | | | | | |
| Favored (%) | 98.34 | 98.68 | 98.42 | 98.59 | 98.47 | 97.81 |
| Allowed (%) | 1.66 | 1.32 | 1.58 | 1.41 | 1.53 | 2.19 |
| Outliers (%) | 0.00 | 0.00 | 0.00 | 0.00 | 0.00 | 0.00 |
| **RNA validation** | | | | | | |
| Average suiteness (%) | 56.8 | 59.0 | 58.1 | 58.7 | 58.5 | 55.4 |
| Good sugar puckers (%) | 99.59 | 99.61 | 99.58 | 99.97 | 99.83 | 99.88 |

*Fourier shell correlation calculated between composite half maps.

| | State K1<br>PDB 8FLA<br>EMD-29272 | State K2<br>PDB 8FLB<br>EMD-29273 | State K3<br>PDB 8FLC<br>EMD-29274 | State L1<br>PDB 8FLD<br>EMD-29275 | State L2<br>PDB 8FLE<br>EMD-29276 | State L3<br>PDB 8FLF<br>EMD-29277 |
|---|---|---|---|---|---|---|
| **Data collection and processing** | | | | | | |
| Microscope | Titan Krios | Titan Krios | Titan Krios | Titan Krios | Titan Krios | Titan Krios |
| Voltage (keV) | 300 | 300 | 300 | 300 | 300 | 300 |
| Camera | K3 | K3 | K3 | K3 | K3 | K3 |
| Magnification | 64,000 | 64,000 | 64,000 | 64,000 | 64,000 | 64,000 |
| Pixel size at detector (Å/pixel) | 1.072 | 1.072 | 1.072 | 1.072 | 1.072 | 1.072 |
| Total electron exposure (e-/Å2) | 60 | 60 | 60 | 60 | 60 | 60 |
| Exposure rate (e-/pixel/sec) | 30 | 30 | 30 | 30 | 30 | 30 |
| Number of frames collected (no.) | 40 | 40 | 40 | 40 | 40 | 40 |
| Defocus range (μm) | -0.5 to -2.5 | -0.5 to -2.5 | -0.5 to -2.5 | -0.5 to -2.5 | -0.5 to -2.5 | -0.5 to -2.5 |
| Energy filter slit width (V) | 20 | 20 | 20 | 20 | 20 | 20 |
| Automation software | SerialEM | SerialEM | SerialEM | SerialEM | SerialEM | SerialEM |
| Micrographs collected (no.) | 172,699 | 172,699 | 172,699 | 172,699 | 172,699 | 172,699 |
| Total extracted particles (no.) | 15,679,142 | 15,679,142 | 15,679,142 | 15,679,142 | 15,679,142 | 15,679,142 |
| Final particle images (no.) | 64,178 | 66,157 | 20,809 | 88,174 | 123,749 | 33,770 |
| Point-group symmetry | C1 | C1 | C1 | C1 | C1 | C1 |
| Resolution (global at FSC-0.143, Å) | | | | | | |
| Overall map (masked) | 2.84 | 2.64 | 3.05 | 2.72 | 2.49 | 2.83 |
| Composite map (masked)* | 2.63 | 2.55 | 2.76 | 2.58 | 2.48 | 2.65 |
| Map sharpening $B$ factor (Å$^2$) | | | | | | |
| Overall map | -36.9 | -42.8 | -21.7 | -41.9 | -46.8 | -29.2 |
| Focused maps | -52.4 to -61.3 | -52.4 to -61.1 | -52.4 to 53.4 | -54.1 to -61.3 | -54.1 to -61.1 | -54.1 to -54.2 |
| **Model composition** | | | | | | |
| Non-hydrogen atoms | 153,587 | 149,312 | 139,873 | 148,845 | 144,415 | 135,006 |
| Chains (proteins/RNA) | 52/4 | 51/3 | 47/3 | 50/4 | 49/3 | 45/3 |
| Protein residues (non-modified/modified) | 9000/2 | 8645/2 | 7624/2 | 8504/1 | 8149/1 | 7132/1 |
| RNA (non-modified/modified) | 3670/109 | 3598/109 | 3551/109 | 3620/111 | 3548/111 | 3501/111 |
| Ligands ($Mg^{2+}$/$Zn^{2+}$/ $K^+$/GDP/GTP) | 88/6/0/1/0 | 88/6/0/1/0 | 88/6/0/1/0 | 86/6/0/0/0 | 86/6/0/0/0 | 86/6/0/0/0 |
| **Model refinement** | | | | | | |
| Refinement package (real space) | Phenix 1.19.1 | Phenix 1.19.1 | Phenix 1.19.1 | Phenix 1.19.1 | Phenix 1.19.1 | Phenix 1.19.1 |
| Initial models used (PDB code) | 6LSS | 6LSS | 6LSS | 6LSS | 6LSS | 6LSS |
| Model resolution cutoff (Å) | 3.10 | 2.90 | 3.40 | 3.00 | 2.80 | 3.10 |
| Model-Map CC ($CC_{mask}$/$CC_{box}$/ $CC_{peaks}$/$CC_{volume}$) | 0.85/0.85/ 0.79/0.84 | 0.86/0.84/ 0.81/0.86 | 0.82/0.81/ 0.74/0.81 | 0.86/0.86/ 0.82/0.86 | 0.88/0.87/ 0.84/0.88 | 0.84/0.82/ 0.77/0.83 |
| Model-to-map FSC, threshold 0.50 (masked/unmasked, Å) | 2.87/2.87 | 2.76/2.76 | 2.97/2.97 | 2.79/2.79 | 2.62/2.62 | 2.88/2.88 |
| **RMSDs** | | | | | | |
| Bond lengths (Å) | 0.003 | 0.005 | 0.004 | 0.005 | 0.005 | 0.003 |
| Bond angles (°) | 0.726 | 0.792 | 0.764 | 0.774 | 0.783 | 0.719 |
| **B-factors** | | | | | | |
| Average B-factors (A$^2$) | | | | | | |
| Protein | 50.13 | 23.84 | 54.24 | 41.09 | 40.25 | 37.49 |
| RNA | 62.70 | 33.89 | 69.81 | 52.19 | 50.74 | 49.23 |
| Ligand | 35.10 | 15.28 | 38.44 | 28.78 | 28.00 | 26.71 |
| **Validation** | | | | | | |
| Clash score | 2.17 | 2.04 | 2.70 | 2.24 | 1.96 | 2.05 |
| MolProbity score | 0.99 | 0.97 | 1.06 | 1.00 | 0.96 | 0.97 |
| CaBLAM outliers | 1.10 | 1.16 | 1.16 | 1.19 | 1.09 | 1.19 |
| Poor rotamers (%) | 0.01 | 0.03 | 0.00 | 0.00 | 0.01 | 0.02 |
| C-beta deviations | 0.00 | 0.00 | 0.00 | 0.00 | 0.00 | 0.00 |
| EMRinger score | 3.74 | 4.33 | 3.72 | 4.26 | 4.43 | 4.03 |
| **Ramachandran plot** | | | | | | |
| Favored (%) | 98.81 | 98.38 | 97.97 | 98.18 | 98.37 | 98.63 |
| Allowed (%) | 1.19 | 1.62 | 2.03 | 1.88 | 1.63 | 1.37 |
| Outliers (%) | 0.00 | 0.00 | 0.00 | 0.00 | 0.00 | 0.00 |
| **RNA validation** | | | | | | |
| Average suiteness (%) | 59.7 | 57.2 | 57.0 | 57.5 | 57.8 | 59.5 |
| Good sugar puckers (%) | 99.80 | 99.83 | 99.77 | 99.88 | 99.99 | 99.88 |

*Fourier shell correlation calculated between composite half maps.

87. D. Liebschner, P. V. Afonine, M. L. Baker, G. Bunkóczi, V. B. Chen, T. I. Croll, B. Hintze, L.-W. Hung, S. Jain, A. J. McCoy, N. W. Moriarty, R. D. Oeffner, B. K. Poon, M. G. Prisant, R. J. Read, J. S. Richardson, D. C. Richardson, M. D. Sammito, O. V. Sobolev, D. H. Stockwell, T. C. Terwilliger, A. G. Urzhumtsev, L. L. Videau, C. J. Williams, P. D. Adams, Macromolecular

structure determination using X-rays, neutrons and electrons: recent developments in Phenix. *Acta Crystallogr Sect D*. 75, 861–877 (2019).
